# Multi-modal skin atlas identifies a multicellular immune-stromal community associated with altered cornification and specific T cell expansion in atopic dermatitis

**DOI:** 10.1101/2023.10.29.563503

**Authors:** Evgenij Fiskin, Gökcen Eraslan, Maria B Alora-Palli, Juan Manuel Leyva-Castillo, Sean Kim, Heather Choe, Caleb A Lareau, Helena Lau, Emily P Finan, Isabella Teixeira-Soldano, Brenna LaBere, Anne Chu, Brian Woods, Janet Chou, Michal Slyper, Julia Waldman, Sabina Islam, Lynda Schneider, Wanda Phipatanakul, Craig Platt, Orit Rozenblatt-Rosen, Toni M Delorey, Jacques Deguine, Gideon P Smith, Raif Geha, Aviv Regev, Ramnik Xavier

## Abstract

In healthy skin, a cutaneous immune system maintains the balance between tolerance towards innocuous environmental antigens and immune responses against pathological agents. In atopic dermatitis (AD), barrier and immune dysfunction result in chronic tissue inflammation. Our understanding of the skin tissue ecosystem in AD remains incomplete with regard to the hallmarks of pathological barrier formation, and cellular state and clonal composition of disease-promoting cells. Here, we generated a multi-modal cell census of 310,691 cells spanning 86 cell subsets from whole skin tissue of 19 adult individuals, including non-lesional and lesional skin from 11 AD patients, and integrated it with 396,321 cells from four studies into a comprehensive human skin cell atlas in health and disease. Reconstruction of human keratinocyte differentiation from basal to cornified layers revealed a disrupted cornification trajectory in AD. This disrupted epithelial differentiation was associated with signals from a unique immune and stromal multicellular community comprised of *MMP12*^+^ dendritic cells (DCs), mature migratory DCs, cycling ILCs, NK cells, inflammatory *CCL19*^+^ *IL4I1*^+^ fibroblasts, and clonally expanded *IL13*^+^*IL22*^+^*IL26*^+^ T cells with overlapping type 2 and type 17 characteristics. Cell subsets within this immune and stromal multicellular community were connected by multiple inter-cellular positive feedback loops predicted to impact community assembly and maintenance. AD GWAS gene expression was enriched both in disrupted cornified keratinocytes and in cell subsets from the lesional immune and stromal multicellular community including *IL13*^+^*IL22*^+^*IL26*^+^ T cells and ILCs, suggesting that epithelial or immune dysfunction in the context of the observed cellular communication network can initiate and then converge towards AD. Our work highlights specific, disease-associated cell subsets and interactions as potential targets in progression and resolution of chronic inflammation.

## Main

Dysregulation of barrier and immune functions in the skin can result in a chronically inflamed tissue ecosystem. Atopic dermatitis (AD) is a prevalent skin inflammatory pathology[1], characterized by immune infiltrates in the upper dermis, epidermal barrier defects[2], and the presence of a core type 2 polarized immune response that can be accompanied by type 22, type 17 and type 1 response signatures[3]. Topical treatment and therapies targeting the type 2 response, such as antagonistic IL4RA antibodies are effective in only ∼60% of AD patients[4] and have to be given continuously[5], highlighting the unmet need to identify distinct subtypes of the disease and suitable interventions.

Despite extensive studies into the genetic and molecular features of the complex interplay between barrier dysfunction and immune activation in AD pathogenesis[6], many interactions remain only partly understood. These include aberrant barrier function and terminal keratinocyte differentiation of cornified keratinocytes, cell-cell interactions between keratinocytes and immune cells associated with pathological inflammation, and the precise states of disease-promoting T cells, their TCR clonotype composition and antigen specificity, and associations with other cell types.

Single cell RNA-seq (scRNA-seq) has been instrumental in defining the cellular landscape in human health and disease[7], providing crucial insights into homeostasis and pathology in different tissues, including the healthy and inflamed human skin. In the skin, scRNA-seq studies have provided an initial cell census and nominated disease-related cellular components, but have been limited to either small patient cohorts[8–10] (n=4 or 5), gene expression profiles, or specific tissue sub-compartments[11–15]. Thus, our understanding of the skin cellular ecosystem and its remodeling upon inflammation remains incomplete.

Here, we generated a large-scale, high resolution, multi-modal single cell atlas of RNA, chromatin, and TCR clonotype profiles in healthy and inflamed human skin. We collected 310,691 scRNA-seq profiles from 43 samples from 19 individuals, including non-lesional and lesional skin from 11 AD patients with mild to severe disease. We combined our analysis with 396,321 cells from four re-annotated public skin scRNA-seq datasets[8,10–12] into an integrated atlas of 707,012 cells, along with deconvoluted bulk RNA-seq profiles from an additional cohort of 27 AD patients[16]. We annotated 86 cell subsets, including multiple previously unknown and disease-specific cellular subsets. Computational inference of the full keratinocyte (KC) differentiation trajectory from basal to cornified layers revealed disruption in AD of the normal cornification process. Joint scRNA-seq and T cell receptor sequencing (TCR-seq) of lesional skin T cells identified a unique, clonally expanded, T cell state with overlapping type 2 and type 17 characteristics that co-expresses the key cytokines *IL13, IL22* and *IL26*. Both cornified keratinocytes and *IL13*^+^*IL22*^+^*IL26*^+^ T cells were enriched for atopic dermatitis risk genes nominated by GWAS. In AD, a DC activation trajectory was shifted towards a unique mature migratory DC (mmDC) state, predicted to promote the pathological T cell response. In AD, *IL13*^+^*IL22*^+^*IL26*^+^ T cells and mmDCs, along with lesion-specific stromal cell subsets, and select neuronal subsets formed a multicellular community through multiple inter-cellular positive feedback loops, generating signals that can impact aberrant KC differentiation and skin pathology in AD.

## Results

### An integrated, multi-modal human single cell atlas of healthy and AD skin

To construct a multi-modal cell atlas of healthy, AD non-lesional and AD lesional adult human skin, we analyzed a cohort of 19 adult individuals, including 11 AD patients, 6 gender-matched healthy controls and 2 scleroderma patients, to a total of 43 specimens, spanning Caucasian, Black, Asian and Hispanic descent (**Figure 1a** and **Figure S1**). Unlike some prior work[10–12], we profiled the entire tissue without flow enrichment, by establishing a tissue dissociation workflow of whole 3mm skin punch biopsies into single cell suspensions. We analyzed cells using droplet based 3’-scRNA-seq, TCR-seq/5’-scRNA-seq and a single-cell assay for transposase-accessible chromatin by sequencing (scATAC-seq) (**Figure 1a, Methods**). We retained 310,691 high-quality 3’-scRNA-seq profiles from 43 samples (7 healthy, 32 AD, 4 scleroderma), 1,138 TCR clonotypes from two samples and 3,220 scATAC-seq profiles from two samples (**Figure 1a** and **Figure S2a-c**; **Methods**). There was good sample mixing across individuals (**Figure S2d**), indicating the absence of strong batch effects and allowing for downstream analysis without batch correction.

**Figure 1.**
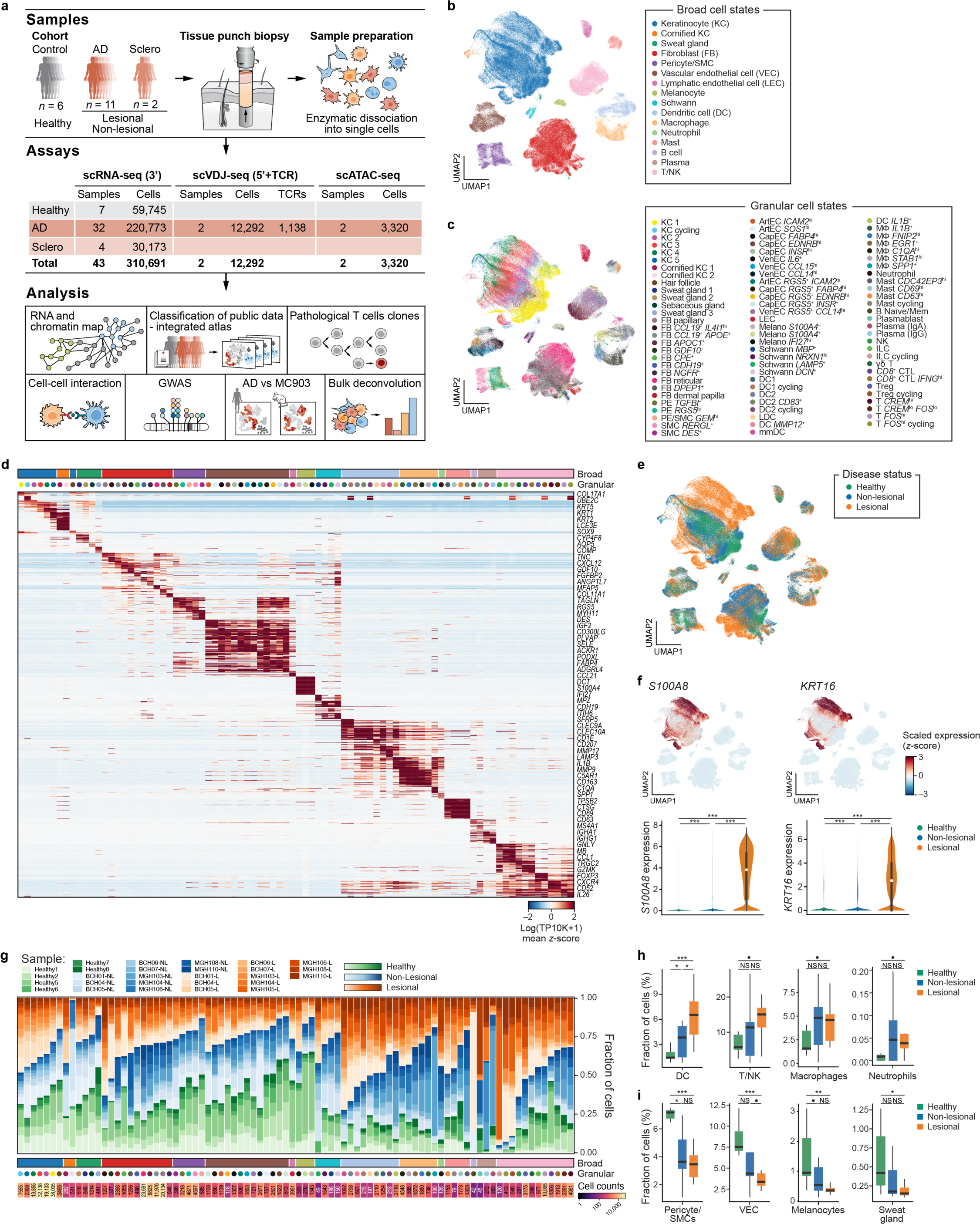
Multi-modal single-cell atlas of healthy, non-lesional and lesional skin of atopic dermatitis patients. **a**, Study overview. **b-f**, Skin cell atlas. Two-dimensional (2D) embedding of scRNA-seq profiles (dots) from all donors, colored by annotated coarse cell types (b), granular cell subsets (c), disease state (e), or z-scored expression values of epidermal inflammatory marker genes (f, along with distribution of their expression values across disease conditions (x-axis), white dot: median expression). **d**, Mean z-scored expression values (color) of marker genes (rows) across 86 granular cell subsets (columns; color code as in **c**). Top bar: coarse cell types (as in **b**). **g-i**, Increased immune cell proportions in lesional skin. **g**, Proportion (y-axis) of granular cell subsets (columns, ordered within broad categories (color bar) by proportion of cells from lesional samples) in healthy (green), non-lesional (blue) and lesional (orange) samples. Color tones: individual patients. Bottom: total number of cells for each subset in the dataset. **h-i**, Distribution of proportions (y-axis) of each coarse cell type that is significantly increased (h) or decreased (i) in lesions (*vs*. healthy controls) in healthy, non-lesional and lesional conditions (x-axis). Center line: median; box limits: first and third quartiles; whiskers: 1.5× interquartile range. •FDR<0.1, *<0.05, **<0.01,***<0.001, ns: non-significant; Dirichlet regression.

Iterative clustering and *post hoc* annotation (**Methods**) resolved 16 broad cell categories (**Figure 1b** and **Figure S2e**) and 86 granular cell subsets (**Figure 1c,d** and **Figure S3a; Supplementary Table 1**), including subsets reflecting previously-unidentified states and rare subsets. These included, for example, *MMP12*^+^ DCs, *CPE*^+^ fibroblasts, *NGFR*^+^ and *CDH19*^+^ Schwann-like fibroblasts that are *CD90/THY1*-negative, multiple rare subsets of skin appendageal cell subsets (containing hair follicles, sweat gland and sebaceous gland subsets), and cycling innate lymphoid cells (ILCs) (**Figure S3a-e**). As expected, KC expression of key epidermal inflammatory markers *S100A8* (FDR < 10^-16^, Wald test; **Methods**) and *KRT16* (FDR <10^-16^, Wald test) was much higher in lesional (inflamed) *vs*. non-lesional or healthy samples (**Figure 1e,f** and **Figure S3f,g**).

To corroborate the cell subsets, and generate a cross-study skin cell atlas, we built a linear classifier based on the 86 granular cell subsets. We then classified profiles from four previous scRNA-seq studies of healthy and AD skin[8,10–12], and co-embedded these re-annotated cell profiles with our data for an integrated skin cell atlas of 707,012 cells (**Figure S4a-g**; **Methods; Supplementary Table 5**). There was overall coherent cell type annotation and composition between prior work and our study (**Figure S4a-g; Supplementary Table 6**), suggesting the robustness and comprehensive nature of the atlas, although some of the fine cell subsets we identified were not captured in previous data (**Figure S4h-j**). Most strikingly, our atlas included cornified keratinocytes and select dermal stromal subsets that may be absent from prior studies as they are either difficult to capture (cornified keratinocytes) or underrepresented in suction blistering, which mainly captures the epidermis (dermal subsets)[11,12] (**Figure S4h-j**).

### Increased DCs, T cells and NK cells in AD skin

Comparison of the overall cellular composition between healthy, non-lesional and lesional AD skin in our dataset revealed an increased abundance of DCs in both non-lesional (FDR = 0.033, Dirichlet regression, **Methods**) and lesional (FDR = 2.34*10^-8^) AD skin compared to healthy skin, as well as of T and natural killer (NK) cells (FDR = 0.078), macrophages (FDR = 0.063) and neutrophils (FDR = 0.018) in lesions (**Figure 1g,h** and **Figure S5; Supplementary Table 2**), consistent with infiltration of immune cells into lesional skin[1]. Conversely, there were lower fractions of pericytes/smooth muscle cells (SMCs) in non-lesional (FDR = 0.027) and lesional (FDR = 1.09*10^-9^) AD skin, and of vascular endothelial cells (FDR = 4.63*10^-8^), melanocytes (FDR = 0.004) and sweat gland cells (FDR = 0.0114) in lesions (**Figure 1g,h** and **Figure S5; Supplementary Table 2**).

### Comprehensive reconstruction of keratinocyte differentiation

Skin barrier formation is accomplished via terminal differentiation of KCs through basal, suprabasal, spinous, granular and cornified layers[17,18]. Graph-based clustering and partition-based approximate graph abstraction (PAGA)[19] of KCs (**Methods**) revealed 8 interconnected KC states that recapitulated known features of KC differentiation (**Figure 2a**): from basal marker-expressing KC1s (*KRT15*, *COL17A1*) towards cycling KC (*UBE2C*, *TOP2A*) and KC2s (*MT1X*, *MT1G*), spinous marker-positive KC3s (*KRT1*, *KRT10*), granular marker-expressing KC4s (*KRT2*, *DSC1*) and KC5s (*CNFN, KLK7*), and finally cornified KC1s (*SPRR2A*, *SPRR2B*) and cornified KC2s (*IL37*, *PSORS1C2, LCE1B*) (**Figure 2a-c** and **Figure S6a-c; Supplementary Table 1**), with gene sets enriched in the different KC subsets consistent with known biology (**Figure S6f; Methods; Supplementary Table 3**). Several lines of evidence support the ordering from KC5 to cornified KC1 and KC2s, including differential expression of late cornified envelope (*LCE) vs*. basal and spinous KC marker genes (**Figure S6d**), higher expression of autophagy and mitophagy genes in cornified KCs (**Figure S6f**)[20], and RNA velocity analysis[21,22] (**Figure S6e**).

**Figure 2.**
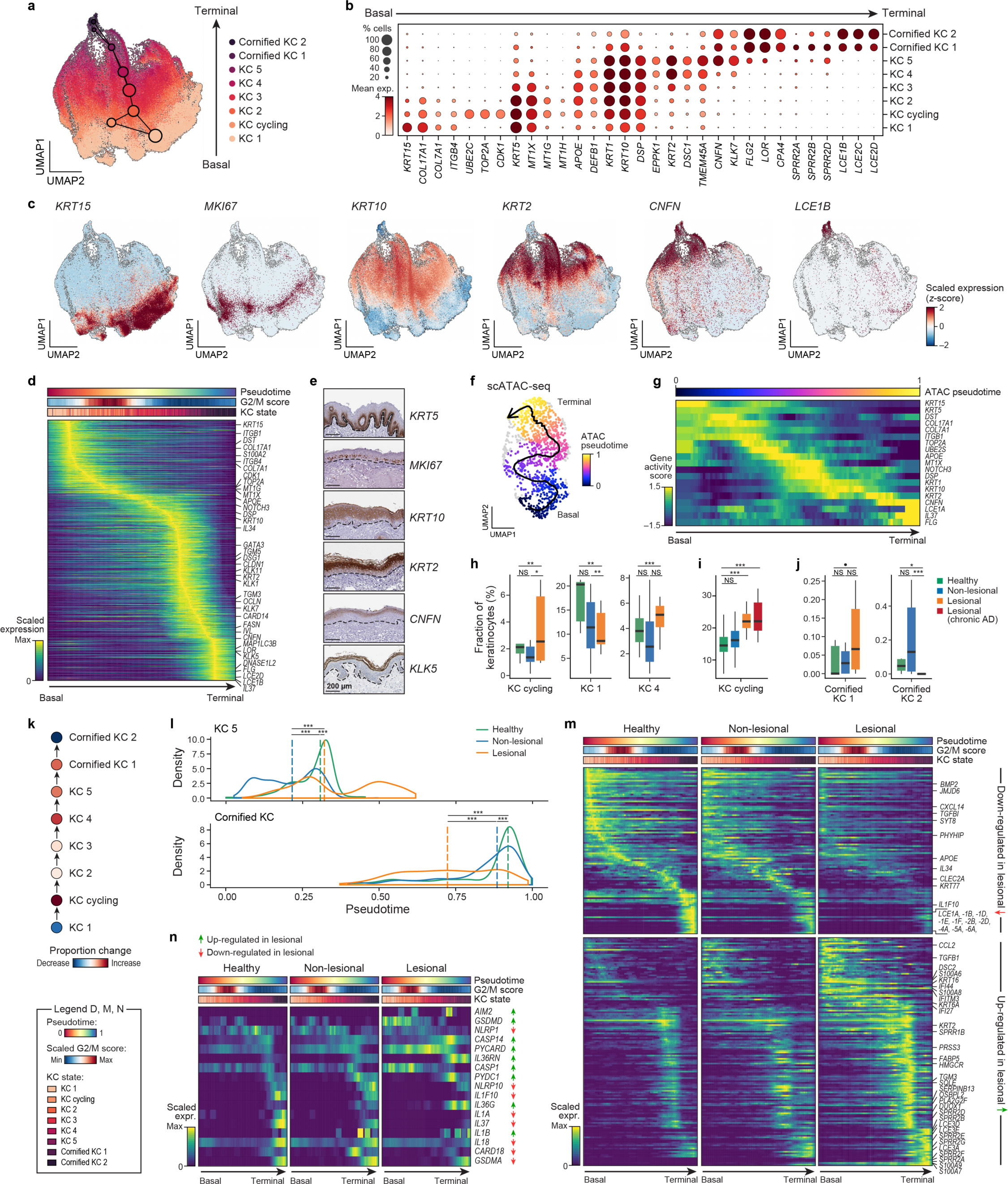
Shift in keratinocyte differentiation in atopic dermatitis. **a-g**, Full keratinocyte differentiation trajectory from basal to terminal cornified layer. **a,c**, 2D embedding and PAGA connectivity map (nodes and edges) of keratinocyte scRNA-seq profiles (dots) from all samples, colored by annotated keratinocyte cell subset (a) or by z-scored expression of keratinocyte subset marker genes (c). Arrow in **a**: differentiation axis. **b**, Mean expression (log(TP10K+1), node color) and proportion of expressing cells (node size) of marker genes (columns) of keratinocyte cell subset (rows). Arrow: keratinocyte differentiation axis. **d**, Scaled expression values (color) of marker genes (rows) in cells (columns) from healthy controls ordered along diffusion pseudotime (DPT) of keratinocyte differentiation from basal (left) to terminal cornified (right) cells. Colored bars: DPT (top), G2/M score (middle) and keratinocyte subsets (bottom), data were downsampled (**Methods**) to show equal numbers of cells for each subset. **e**, Immuno-histochemistry (IHC) derived from the Human Protein Atlas[74] of healthy skin tissue sections with subset markers. Dotted line: border between epidermis (top) and dermis (bottom). Scale bar: 200µm. **f,g**, Keratinocyte cell differentiation by scATAC-Seq. **f,** 2D embedding of keratinocyte scATAC-seq profiles from a lesional biopsy, colored by diffusion pseudotime (DPT). Arrow: inferred trajectory (**Methods**). **g**, ScATAC-seq gene activity scores (color) of marker genes (rows) in cells (columns) from **f** ordered by DPT (top bar). Arrow: differentiation axis. **h-m**, Shift towards cornified KC1s in differentiation in lesional skin. **h-j**, Proportions (y-axis) of keratinocyte cell subsets (x-axis) in scRNA-seq (h,j) or deconvolved (i, **Methods**) bulk RNA-seq data[16]. Center line: median; box limits: first and third quartiles; whiskers: 1.5× interquartile range. **k**, Changes in keratinocyte proportions along differentiation. Changes in cell proportions (color) in each cell type (node) (red, increase; blue, decrease; scale bar, bottom). **l**, Aberrant KC differentiation trajectories in AD skin. Distribution of DPT (**Methods**) for KC5 (left) and cornified KC (right) cells, colored by disease state.*** BH FDR<0.05 (Mann-Whitney U test). Vertical dashed lines: median. **m,n**, Scaled expression (color) of differentially expressed genes (rows) that are decreased (m, top), increased (m, bottom) or are *IL-1* family genes (n) across DPT-ordered cells (columns) in lesional (right), non-lesional (middle), or healthy (left) skin. Arrows: keratinocyte differentiation axis. Data is downsampled (**Methods**). Color bars as in **d**. Colored arrows in **m,n**: increased (green) or decreased (red) expression in lesional skin.

Next, reconstructing healthy human KC differentiation as a continuous process (by calculating diffusion pseudotime (DPT)[23] for KCs from all healthy samples; **Methods**), we observed a consistent ordering, and captured dynamic gene expression changes along differentiation (**Figure 2d; Methods**), aligned with the spatial localization of corresponding marker proteins in Human Protein Atlas (**Figure 2e**). This highlighted the temporal patterns of key known differentiation transcription factors (TFs) (*TP63, GATA3, GRHL3, GRHL1, PRDM1*)[24–26] and novel ones (*e.g.*, *NFE2, NPAS1, ETV7*) (**Figure S6g,h**), as well as monogenic skin disease genes. For example, expression of epidermolysis bullosa-causing genes peaked at early DPT points in basal KCs, whereas ichthyosis-causing genes peaked at late DPT points in upper granular and cornified layers (**Figure S6i**), in line with roles in KC attachment to the basement membrane and lipid cornified envelope formation, respectively.

We further validated our reconstructed differentiation by showing congruent dynamic remodeling of chromatin accessibility along keratinocyte differentiation from 3,220 high quality scATAC-seq profiles in two samples (**Figure S7a,b**; **Methods**). ScATAC-Seq recovered major cell types at proportions consistent with scRNA-seq (**Figure S7c**), with high chromatin accessibility (**Methods**) around cell type marker genes (**Figure S7d**). Ordering the scATAC-seq profiles and deriving gene activity scores along the DPT axis (**Methods**) yielded patterns consistent with those from scRNA-seq, with a temporal progression from high basal gene activity scores towards spinous, granular and cornified KCs (**Figure 2f,g** and **Figure S7e,f**), and similar patterns for TFs (**Figure S7g**).

### Altered keratinocyte differentiation and cornified layer in lesional AD skin

Skin inflammation in AD altered the KC trajectory and states of KCs, with remodeling of the cornified KC layer and a shift from cornified KC2 towards cornified KC1. First, there was an increase in the fraction of cycling KCs (FDR = 0.002, Dirichlet regression) and granular KC4s (FDR = 1.45*10^-6^) in lesional AD skin, and a decrease in basal KC1s (FDR = 0.004) (**Figure 2h,i** and **Figure S8a; Supplementary Table 2**), and a shift from cornified KC2s dominant in healthy samples (FDR = 0.018) to near-exclusive cornified KC1s in lesional AD (FDR = 0.078) (**Figure 2j,k** and **Figure S8b; Supplementary Table 1**). This was related to an altered KC differentiation trajectory in lesional AD compared to healthy or non-lesional skin based on diffusion pseudotime (DPT, **Figure 2l**; **Methods**), where lesional KC5s had a significantly later DPT with one particularly delayed mode and lesional cKCs are primarily cKC1s (with earlier DPT) (**Figure 2l**; FDR<0.05, Mann–Whitney U test). This suggests a model where lesional AD is characterized by a shift from basal to cycling KCs, followed by decreased differentiation from KC5s to cKCs, leading to defective cornification. This shift was accompanied by decreased cell intrinsic expression of late cornified envelope genes in lesional AD KCs (*e.g.*, *LCE5A, LOR*) and an increase in small proline rich proteins (*SPRR*) (**Figure 2m; Supplementary Table 4**), also captured by bulk RNA-Seq[16] (**Figure S8c**). Remodeling of the cornified KC layer and the shift from cornified KC2 towards cornified KC1 were not observed in non-lesional and lesional samples from two scleroderma patients (**Figure S8f,g**), emphasizing unique lesional AD-driven alterations to cornification.

There was also increased cell intrinsic expression in lesional KCs of epidermal inflammatory markers, interferon pathway genes, antimicrobial response genes, lipid and rate-limiting cholesterol metabolism genes (**Figure 2m** and **Figure S8d; Supplementary Table 3**)[8,10,13], and IL-20-related cytokines *IL19* and *IL24* (**Figure S8e**), implicated in epidermal hyperplasia and barrier dysfunction in AD[27]. Across all KCs, cornified KCs preferentially expressed IL-1 family cytokines and regulators (*IL1A, IL1B, IL18, IL36G, IL37, IL1F10/IL38, IL36RN*) and inflammasome-related pathway genes (*GSDMA, NLRP10, PYDC1*) (**Figure 2n** and **Figure S8d**), consistent with prior *in vitro* observations[28]. While pro-inflammatory *IL36G* was up-regulated in lesional cornified KCs (FDR < 10^-16^, Wald test), anti-inflammatory cytokines *IL37* and *IL1F10/IL38* were down-regulated (FDR < 10^-16^ and FDR = 3.39*10^-6^).

ScRNA-seq of dissociated dorsal skin from control and MC903-treated mice, a model of AD-like skin inflammation[29] mimicked many aspects of human epidermal inflammation (**Figure S9a-d, Methods**), consistent with previous work[30]. Analogous to human AD, MC903-treated inflamed skin displayed a loss of cornified KCs (**Figure S9f,g**), increased antimicrobial peptides and *Sprrs*, decreased levels of *Lce* genes and increased cycling KCs (**Figure S9h,i**). However, unlike in human AD, *Il4*^+^/*Il1*3^+^ basophils arose in MC903-treated skin (**Figure S9g,j**), as another source of type 2 cytokines (**Figure S9j**), consistent with reports of Il4^+^ basophils as an essential driver of pathology in the MC903 model[31]. Increased numbers of *Il4*^+^/*Il13*^+^ basophils as well as decreased expression of cornification genes have been noted in the murine OVA sensitization model of AD-like skin inflammation, indicating that observed changes are congruent across various mouse AD disease models[32].

### Two DC maturation trajectories in healthy, non-lesional and lesional skin

Myeloid cells from healthy and AD skin spanned a continuum of cell states, which we broadly partitioned into multiple *C1QA*-high macrophage (MΦ) subsets (*C1QA, MS4A7, MAF, CCL18, CCL13*), inflammatory *IL1B*^+^ MΦ (*IL1B, MMP9*), lipid-associated *SPP1*^+^ MΦ (*SPP1, LPL*)[33], tissue-resident Langerhans dendritic cells (LDCs) (*CD207/Langerin, FCGBP*), DC1s (*CLEC9A, XCR1*) and DC2s (*CD1C, FCER1A*)[34], *CD83*^+^ DC2s, inflammatory *IL1B*^+^ DCs (previously coined DC3s[35]; *IL1B, IL23A, CXCL8*), newly identified *MMP12*^+^ DCs (*MMP12, CD1B, RAB7B*), mature migratory DCs (mmDCs) (*LAMP3, CCR7*; also referred to as mature DCs enriched in immunoregulatory molecules (mregDCs)[36]), cycling DC1s and DC2s (*UBE2C, MKI67*), as well as neutrophils (*CXCL8, FCGR3B, S100P*) (**Figure 3a,b** and **Figure S10a; Supplementary Table 1**).

**Figure 3.**
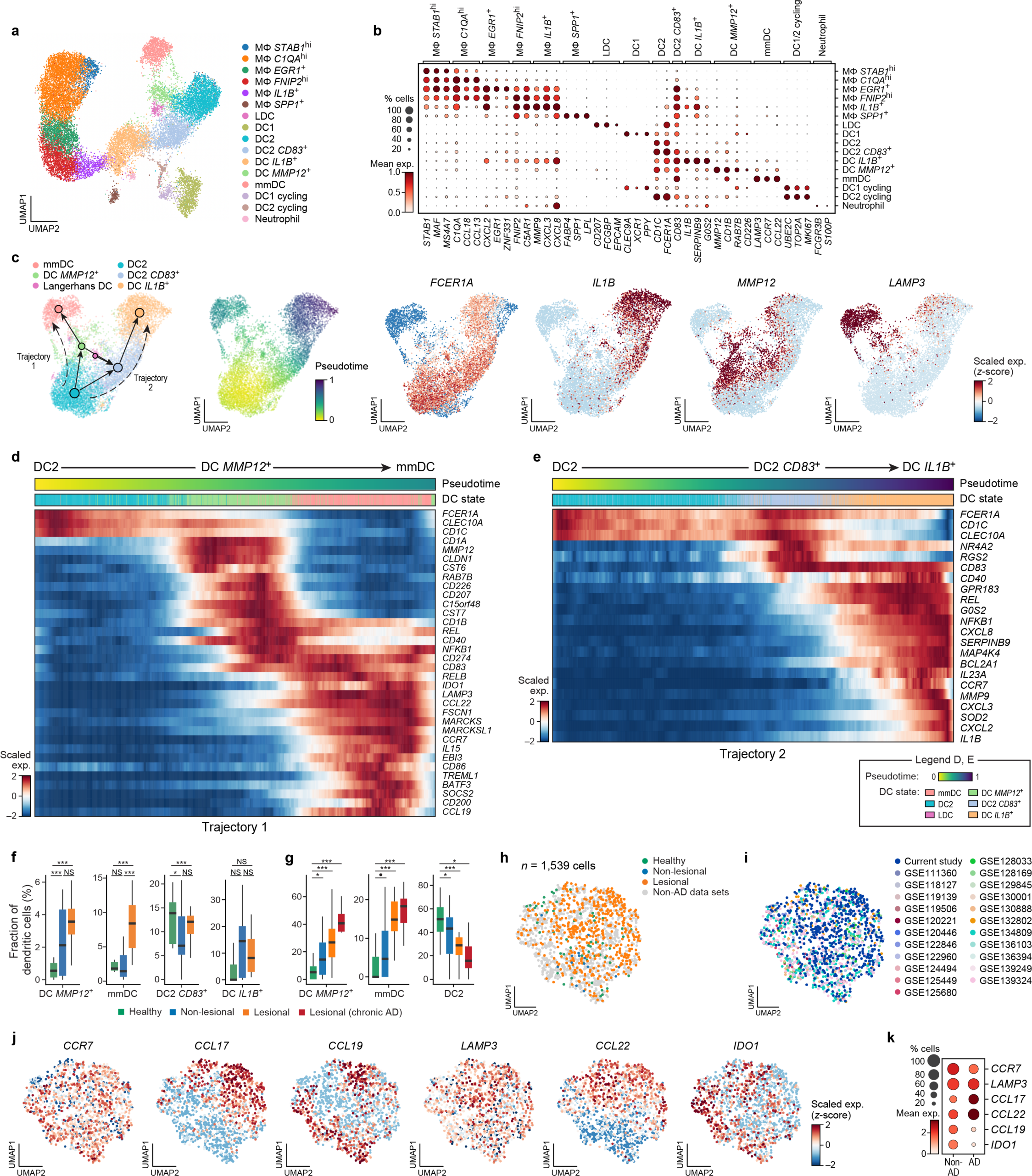
Increase in *MMP12*^+^ DCs and mmDCs in AD through an activation trajectory. **a,b**, Skin myeloid cells. **a**, 2D embedding of myeloid scRNA-seq profiles (dots), colored by cell subset. **b,** Mean (dot color) expression and fraction of expression cells (dotsize) of myeloid cell markers (columns) across cell subsets (rows). **c-e**, Two trajectories of dendritic cells activation. **c**, 2D embedding of selected DC scRNA-seq profiles (dots) and directed PAGA connectivity map (nodes and edges) from all samples, colored (from left) by cell subset, DPT, or z-scored expression of marker genes. Dashed arrows: activation trajectories. **d,e**, Expression (colorbar) of marker genes (rows) in cells (columns) from all samples ordered by DPT of DC activation trajectory 1 (d) or 2 (e). Colored bars: DPT (top) and cell annotation (bottom, as in **c**). Data were down sampled (**Methods**) to show equal numbers of cells for each subset. **f,g**, Increased proportion of DC *MMP12*^+^ and mmDC cells in lesional skin. Proportion (y axis) of different DC cell subsets in scRNA-seq (f) or deconvolved bulk RNA-seq[16] (g). Center line: median; box limits: first and third quartiles; whiskers: 1.5× interquartile range. **h-k**, A *CCL22*^hi^ *CCL17*^hi^ *CCL19*^lo^ *IDO1*^lo^ state enriched in mmDC from AD skin. **h-j**, 2D embedding of mmDCs profiles from our data and 22 additional scRNA-seq studies (**Methods**) colored by condition (h), study (i), or marker gene expression (j). **k**, Mean expression (dot color) and proportion of expressing cells (dot size) of mmDC marker genes (rows) in mmDCs from AD or non-AD samples (columns).

In DCs we inferred two distinct maturation trajectories, both originating from DC2s and transitioning either through *MMP12*^+^ DCs towards mmDCs (Trajectory 1) or via *CD83*^+^ DC2s towards inflammatory *IL1B*^+^ DCs (Trajectory 2) (**Figure 3c-e** and **Figure S10b-d**). Additional paths involving Langerhans cells are also suggested in the model, but were not fully resolved (**Figure 3c**).

Within trajectory 1, *MMP12*^+^ DCs constituted a *CD40*^hi^*CD83*^hi^*CD86*^lo^*LAMP3*^lo^*CCR7*^lo^ transition state, suggesting an activated, but non-fully mature DC state (**Figure 3d** and **Figure S10d**) preceding mmDCs. Consistent with DC activation, *MMP12*^+^ DCs highly expressed antigen presentation genes (*HLA-DRB1, HLA-DQB1, CD1B, CD1A*), maturation-controlling TFs (*REL, NFKB1, SPI1, IRF4*) and co-stimulatory membrane proteins (*CD226, CD40, CD83, TNFRSF4/OX40*) (**Figure 3d** and **Figure S10d**). Apart from expressing the key AD disease marker gene *MMP12*[37], these *MMP12*^+^ DCs also expressed genes encoding select protease inhibitors (*CST6, CST7*) and Langerhans cell markers (*CLDN1, CD207/Langerin*) (**Figure 3d** and **Figure S10b**), indicating the presence of activated LDCs among *MMP12*^+^ DCs. Along trajectory 1, *MMP12*^+^ DCs progressed towards mmDCs, which expressed reported mregDC signatures[36,38,39], including the key TF, *IRF4*[40], actin cytoskeleton and migration genes, DC maturation markers, an immunomodulatory program (*CD274/PD-L1, PDCD1LG2/PD-L2, CD200, IDO1*)[8,10–12,36], *CCL22* and *CCL17*, AD disease markers involved in recruitment of *CCR4*^+^ type 2 inflammatory T cells[41], TSLP alarmin receptor (*CRLF2*, *IL7R*)[12], and the cytokines *IL15* and IL27 subunit *EBI3* (**Figure 3d** and **Figure S10d**), as noted previously[8,10–12,36]. MmDCs profiles had striking similarities to *IRF4*^+^ PD-L2^+^ mouse mature DCs (**Figure S10e**), which are essential regulators of type 2 immunity to protease allergen and worm infection[40,42].

In trajectory 2, DC2s transitioned via a *CLEC10A*^hi^ *FCER1A*^hi^ *CD83*^+^ transition state (*CD83*^+^ DC2s) towards *IL1B*^+^ DCs, which were characterized by high expression of distinct TFs (*CEBPB, HIF1A, STAT3*), select chemokines (*CXCL2, CXCL3, CXCL8, CCL20*), DC maturation markers (*CCR7, LAMP3*) and abundant expression of *IL1B* and psoriasis-associated *IL23A* (**Figure 3e**), an expression signature consistent with reported skin DC3s[35]. The two sets of TFs highlighted in the two trajectories are part of a mutually exclusive gene circuit previously described in bone marrow derived mouse DCs (**Figure S10e**)[43].

### Enhanced differentiation leads to enriched mmDCs in AD

Across DCs, both *MMP12*^+^ DCs and mmDCs were highly increased in lesional skin (FDR = 3.57*10^-6^, Dirichlet regression and FDR = 1.52*10^-7^, respectively), with *MMP12*^+^ DCs also enriched in non-lesional samples compared to healthy controls (FDR = 3.33*10^-5^) (**Figure 3f; Supplementary Table 2**), while the proportion of DC *IL1B^+^* and *CD83^+^* DC2s was unchanged (**Figure 3f** and **Figure S10f,g**). The fractions of *MMP12^+^* DCs and mmDCs were also increased in non-lesional and lesional skin in scRNA-seq from two other AD patient cohorts annotated by our classifier[8,11] (**Figure S10h,i**; **Supplementary Table 5; Methods**) and when deconvolving bulk RNA-seq from AD patients[16] (**Figure 3g** and **Figure S10j**; **Methods**). *MMP12*^+^ DCs and mmDCs in lesional skin had increased predicted *STAT6* TF activity (**Figure S10k; Supplementary Table 7; Methods**), which could impact mmDC activation or differentiation, as seen for IL4-driven PD-L2^+^ mature DCs in mice[40].

Comparing differentiation pseudotimes for Trajectory 1, shows a shift in differentiation between healthy, non-lesional and lesional DCs. Lesional mmDCs had a later DPT compared to healthy and non-lesional counterparts (Kolmogorov–Smirnov (KS) test; FDR<10^-3^ and FDR<0.1, respectively), while non-lesional *MMP12*^+^ DCs had a later DPT compared to healthy and lesional cells (FDR<0.05 and FDR<10^-14^) (**Figure S10l**). Along with the increased fraction of *MMP12*^+^ DCs and mmDCs in lesional skin, this is consistent with a model where a partially activated state arises in non-lesional *MMP12*^+^ DCs that becomes a fully activated mmDC state in lesional AD.

As mmDCs/mregDCs were reported in various inflammatory diseases[36,38,39,44], we compared the distinguishing features of mmDCs in AD to the human mmDC/mregDC landscape. We assembled a human mmDC atlas of 1,092 AD mmDCs from our study (**Figure 3h,i; Methods**) and 1,539 mmDCs expressing our core mmDC signature from 22 scRNA-seq datasets across tissues and diseases (**Methods; Supplementary Table 5** and **8**). Compared to other mmDCs, those from lesional AD skin constitute a *CCL22*^hi^ *CCL17*^hi^ *CCL19*^lo^ *IDO1*^lo^ state (**Figure 3j,k**), consistent with a function in polarizing towards a type 2 immune response.

### A T cell inflammatory program including *IL13*, *IL22* and *IL26* is active in CD4 and CD8 T cells in skin

The 16 lymphocyte subsets included *CD3D-*negative NK cells (*KLRD1, KLRF1, GNLY*^hi^) and ILCs (*KIT, MB, SPINK2*), γδT cells *(TRGC2, FXYD2, ZNF683/HOBIT)*, *CD8A*^+^ cytotoxic T cells (*CD8A, CCL5, GZMK*), *CD4^+^*regulatory T cells (Tregs) (*CTLA4, TIGIT, FOXP3, CCR8*), multiple *CD40LG*^+^ T cells with T *CREM*^hi^ (CREM, *CXCR4*) and T *FOS*^hi^ (*FOS, IL32*) states in both *CD4*^+^ and *CD8*^+^ T cells, naive/memory B cells (*MS4A1/CD20, CD79A, SELL/CD62L*), and plasmablasts (*FKBP4, IRF8*) and IgA or IgG-expressing plasma cells (*JCHAIN, MZB1, TNFRSF17*) (**Figure 4a,b** and **Figure S11a-e; Supplementary Table 1**).

**Figure 4.**
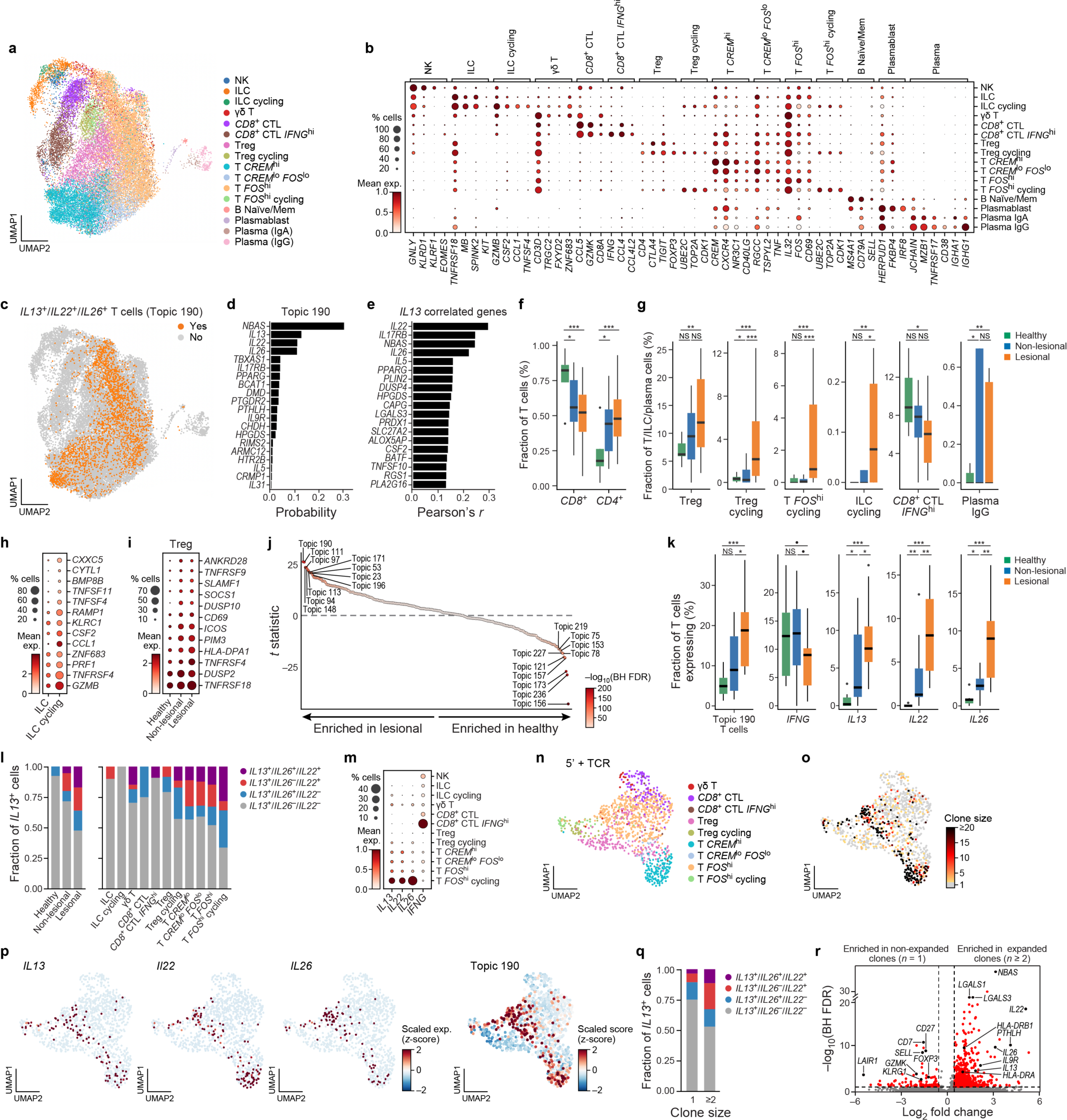
An *IL13/IL22/IL26* program in T cell clones expanded in AD. **a,b**, Skin lymphocytes. **a**, 2D embedding of lymphocyte scRNA-seq profiles (dots) from all samples, colored by annotation. **b**, Mean expression (dot color) and percent of expressing cells (dot size) of marker genes (columns) across cell subsets (rows). **c-e**, An *IL13^+^/IL22^+^/IL26^+^*effector-cytokine program expressed in T cells. **c**, 2D embedding of lymphocyte scRNA-seq profiles (as in **a**), colored by binarized expression of *IL13/IL22/IL26* topic 190 (**Methods**). **d**, Probability (x axis) of top scoring genes (y axis) from the *IL13/IL22/IL26* program (topic 190). **e**, Correlation coefficient (Pearson’s r) of genes (y axis) with highest correlation to *IL13*-expression across single lymphocyte profiles. **f,g**, Cell proportion remodeling in AD. Proportions (y axis) of specific cell subsets across healthy, non-lesional and lesional samples (x axis). **h,i**, Cell intrinsic changes in AD. Mean expression (dot color) and percent of expressing cells (dot size) of selected lesion-enriched genes (rows) in ILC subsets (h, columns) or Tregs from different conditions (i, columns). **j-m**, *IL13/IL22/IL26* co-expressing T cells emerge in lesional skin and are enriched in cycling T cells. **j**, Differential topic expression (*y* axis, t statistic) between healthy and lesional samples for each topic (dot, colored by FDR, colorbar) ranked ordeted (x axis) by t statistic. **k**, Fraction of lymphocytes (y axis) expressing (from left) topic 190 or effector cytokines in each condition (x axis). **l**, Fraction (y axis) of single, double or triple positive *IL13/IL22/IL26* co-expressing cells (colors) in different conditions (left) or lymphoid cell subset (right). **m**, Mean expression (min-max scaled log(TP10K+1), dot color) and percent of expressing cells (dot size) of *IFNG*, *IL13*, *IL22*, *IL26* (columns) across lymphocyte cell subsets (rows). **n-r**, Expanded T cell clones in AD skin are enriched for *IL13/IL22/IL26* program expression. **n-p**, 2D embedding of lymphocyte 5’ scRNA-seq profiles from lesional skin of two patients (**Methods**), colored by cell subset (n), clone size (o), z-scored expression of cytokines (p, left) or z-scored scaled topic 190 score (p, right). **q**, Fraction (y axis) of single, double or triple positive co-expressing cells (bar color) in non-expanded and expanded T cell clones (x axis). **r**, Effect size (x axis, log2 fold change) and significance (y axis, -log10(FDR)) (**Methods**) of expression between expanded and non-expanded T cells. Boxplots: Center line: median; box limits: 1^st^ and 3^rd^ quartiles; whiskers: 1.5× interquartile range.

Because many skin lymphocytes vary continuously — rather than in discrete subsets — and can share cellular processes[45], we characterized lymphocyte gene programs using stochastic block modeling (SBM)[46], a hierarchical topic modeling approach (**Methods**). While some programs reflected distinct cell types, such as regulatory (topic #148) and cytotoxic (#227) T cells (**Figure S11f; Supplementary Table 9**), others reflected cell biological processes (cell cycle, #97), or key immune processes (# 196).

A prominent inflammatory immune program (#190) was expressed across multiple T cell subsets (**Figure 4c,d**), and included key type 2 or 17 genes, such as cytokines *IL13*, *IL5*, pruritogenic *IL31*, *IL22* and *IL26*, *IL25* alarmin receptor *IL17RB* and arachidonic acid, prostaglandin, and eicosanoid pathway genes (*TBXAS1, HPGDS, PTGDR2* (*CRTH2*), *SLC27A2, ALOX5AP*, *PLA2G16*) (**Figure 4e** and **Figure S11g,h**; **Methods**), including *PPARG*, which has been implicated in Th2 cell differentiation and activation[47,48]. *IL31* expression may allow interaction with dorsal root ganglia (DRG) NP3 neurons expressing *IL31* receptor subunits (*IL31RA, OSMR*), based on scRNA-seq of 2,518 non-human primate DRG neurons[49] (**Figure S14e-g**; **Methods**). Both *CD4^+^* and *CD8^+^*T cells expressed this program, most prominently *CREM* and *FOS*^hi^ T cells (log OR=1.269, p-value (of positive association) 1.28×10^-167^, Fisher’s exact test), whereas Tregs and *CD8*^+^*GZMK*^+^ cytotoxic lymphocytes typically did not (log OR=-1.883, p-value (of negative association) 9.8×10^-85^, Fisher’s exact test, **Figure S11i,j**), as expected for an inflammatory effector program.

### A shift from cytotoxic to pro-inflammatory *IL13*/*IL22*/*IL26* expressing T cell clones in lesional AD skin

In contrast to a largely *CD8*^+^ cytotoxic T cell compartment in healthy skin, both non-lesional and lesional AD had a higher proportion of *CD4*^+^ T cells (p-values = 0.017 and 0.001, Dirichlet regression) and an increased fraction of non-cycling and cycling Tregs (FDR = 0.002, Dirichlet regression and FDR = 1.23*10^-8^), cycling *FOS*^hi^ T cells (FDR = 4.2*10^-9^), cycling ILCs (FDR = 0.002) and *IgG*^+^ plasma cells (FDR = 0.003) (**Figure 4f,g** and **Figure S11k; Supplementary Table 2**). Cycling ILCs in lesional skin expressed cytotoxic genes (*GZMB, KLRC1, PRF1*), *CCL1*, and key inflammatory mediators *CSF2/GM-CSF*, the OX40 ligand *TNFSF4*[50], and the cytotoxicity and tissue-residency TF *ZNF683/HOBIT*[51,52] (**Figure 4h**). Tregs in lesional skin expressed higher levels of *TNFRSF9* (4-1BB), *TNFRSF4* (OX40), *TNFRSF18* (GITR) and T cell activation markers (*CD69, HLA-DPA1*) (**Figure 4i; Supplementary Table 4**).

In line with an AD-specific shift from cytotoxic to pro-inflammatory T cell states, there was a prominent increase in T cells expressing the inflammatory immune program (#190, FDR<5%), along with Treg (#148), cell cycle (#111, #97, #94, #23) and T cell activation/differentiation programs (#53, #113, #196) (**Figure 4j,k** and **Figure S11f**; **Supplementary Table 9**), and a decrease in cytotoxic programs (#227, #78) (**Figure 4j,k** and **Figure S11f**; **Supplementary Table 9**). Consistently, an increased fraction of cells expressed *IL13*, *IL22*, and *IL26* in both non-lesional and lesional AD skin (FDR<5%, Mann-Whitney U test) and a smaller fraction expressed *IFNG* in lesional AD (FDR<10%) *vs*. healthy samples (**Figure 4k** and **Figure S11l**)[8,12]. In lesional skin, T cells expressing two or three of *IL13*, *IL22* and *IL26* emerged, especially among cycling *FOS*^hi^ T cells, such that most (>66%) *IL13*^+^ cells also co-expressed at least one of *IL22* and *IL26* (**Figure 4l,m**). Combined TCR- and 5’RNA-seq of lesional biopsies from two patients (**Figure 4n,o; Supplementary Table 10; Methods**) detected 440 expanded (n ≥ 2) and 698 non-expanded (n = 1) T cell clones (**Figure 4o** and **Figure S11m,n**), with strong enrichment of *IL13/IL22/IL26* double- and triple-positive T cells and topic 190 expression in expanded *vs*. non-expanded T cells (**Figure 4p-r** and **Figure S11o-q; Supplementary Table 11**). This suggests the presence of a disease-associated, clonally expanded, pro-inflammatory T cell population with overlapping type 2 and type 17 characteristics.

### Remodeling of the AD stroma with increase in immunoregulatory fibroblasts

Stromal cells in skin spanned eleven subsets of fibroblasts, eight subsets of vascular and lymphatic endothelial cells (**Figure 5a-c**,), as well as 5 pericyte subsets (**Figure S12a,b**), smooth muscle cells, and myelinating and non-myelinating Schwann cells (**Table S1**). The eleven fibroblasts subsets (**Figure 5d,e** and **Figure S12c,d**) included two subsets of *CCL19*^+^ fibroblasts enriched for immune regulating functions: *CCL19*^+^*IL4I1*^+^ fibroblasts (*IL32, TNC, VCAM1*) expressing high levels of MHC-I and II presentation genes (**Figure 5f** and **Figure S12e**) and *CCL19*^+^*APOE*^+^ fibroblasts (*RBP5, TNFRSF13B, CXCL1*) (**Figure S12e**), both expressing markers previously reported in *CCL19*^+^ and *ADAMDEC1*^+^ fibroblasts in other human pathologies, including ulcerative colitis[53,54]. Fibroblasts also included two subsets expressing markers of recently described nerve-associated fibroblasts[55]: *CDH19*^+^ cells (*SCN7A, ANGPTL7, EBF2*) and *NGFR*^+^ cells (*ITGA6, TNNC1, WNT6*) (**Figure 5d,e** and **Figure S12f,g**). In addition to recently-reported *THY1*^+^ skin stromal cells[15], our sorting-free workflow revealed multiple *THY1*^-^ stromal cell populations, including *CDH19*^+^ and *NGFR*^+^ cells and *RERGL*^+^ and *DES*^+^ smooth muscle cells.

**Figure 5.**
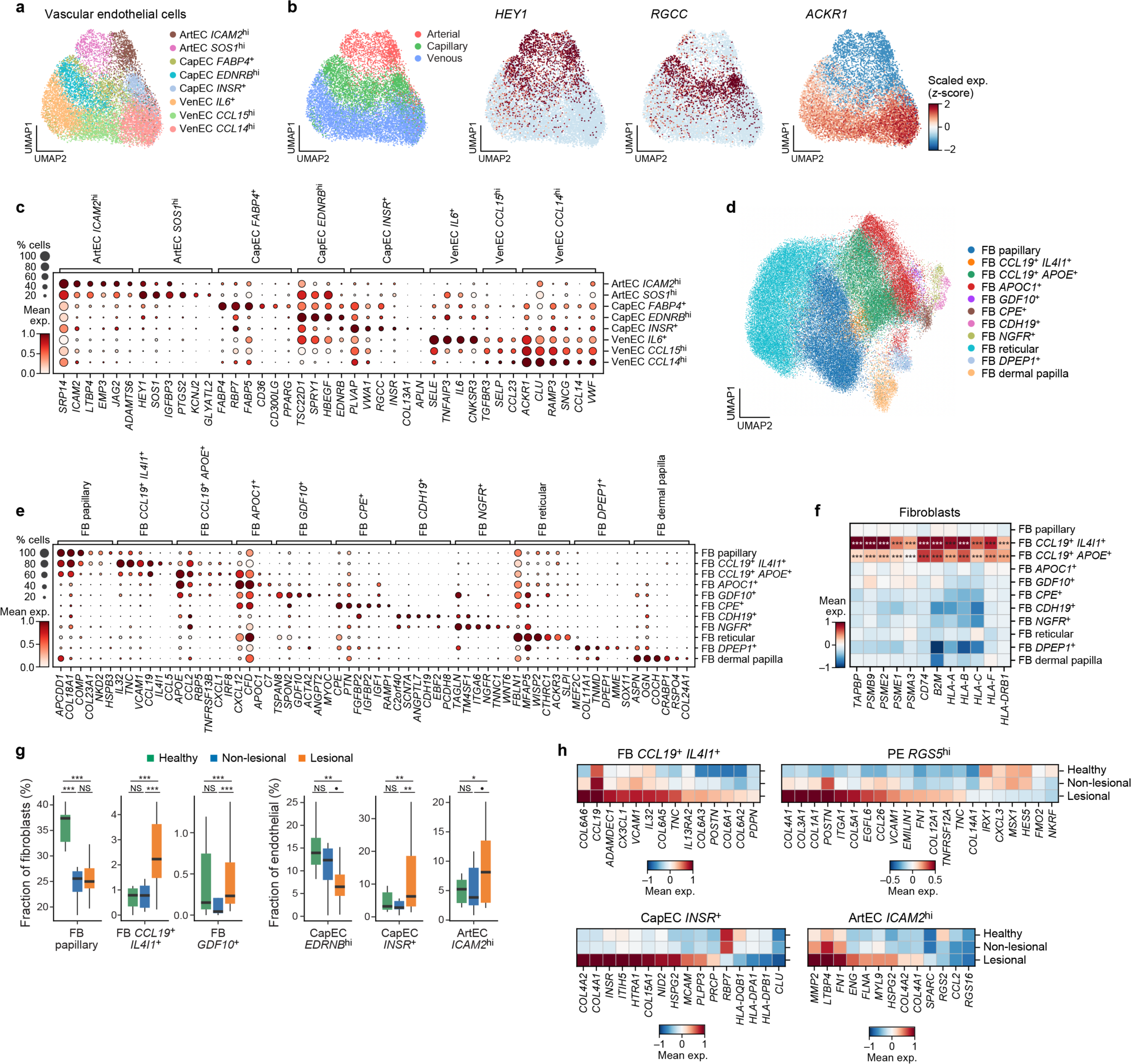
Increase in immunoregulatory fibroblasts in AD skin stroma. **a-c**, Skin vascular endothelial cells. **a,b**, 2D embedding of vascular endothelial cell scRNA-seq profiles from all samples, colored by annotation (a), blood vessel type (b, left), or z-scored expression of vessel marker genes (b, right). **c**, Mean expression (dot color) and percent of expressing cells (dot size) of marker genes (columns) across vascular endothelial cell subsets (rows). **d-f**, Skin fibroblasts. **d**, 2D embedding of fibroblasts scRNA-seq profiles from all samples, colored by granular annotations. **e**, Mean expression (dot color) and percent of expressing cells (dot size) of marker genes (columns) across subsets (rows). **f**, *CCL19*^+^ *IL4I1*^+^ fibroblasts preferential express antigen presentation genes. Z-scored mean expression (color) of HLA class I and II genes (columns) across fibroblast subsets (rows). P values: ***<0.01, negative binomial regression. **g,h**, *CCL19*^+^ *IL4I1*^+^ fibroblasts increase in AD and activate an inflammatory expression program. **g**, Proportions (y axis) of stromal cell subsets (x axis) across conditions (bar color). Center line: median; box limits: 1^st^ and 3^rd^ quartiles; whiskers: 1.5× interquartile range. **h**, Z-scored mean expression of genes (columns) differentially expressed between disease conditions (rows) for stromal cell subsets (top).

Stromal subset composition changed substantially in AD lesions *vs*. healthy skin, with significantly higher fractions of *CCL19*^+^*IL4I1*^+^ fibroblasts (FDR = 2*10^-6^), *GDF10*^+^ FB (FDR = 7.5*10^-4^), *INSR*^+^ CapEC (FDR = 0.005) and *ICAM2*^hi^ ArtEC (FDR = 0.011) and lower proportion of papillary fibroblasts (FDR = 4*10^-4^) and *EDNRB*^hi^ CapECs (FDR = 0.007) (**Figure 5g** and **Figure S12h-j; Supplementary Table 2**). Consistent with a recent report[8], *CCL19*^+^*IL4I1*^+^ fibroblasts showed the most pronounced increase in AD lesions, which we further confirmed in re-annotated published AD single cell[8,11] and deconvoluted bulk RNA-seq[16] (**Figure 5g** and **Figure S12k**). Compared to counterparts in healthy skin, *CCL19*^+^*IL4I1*^+^ fibroblasts in lesional skin had increased expression of type VI collagen genes, *PDPN*, the *IL13* decoy receptor *IL13RA2*, T cell and neutrophil recruiting chemokines (*CCL19*, *CXCL1*) and regulators of immune cell adhesion and tissue repair (*VCAM1, POSTN*) (**Figure 5h; Supplementary Table 4**). These are similar to features of *CCL19*^+^ fibroblasts in the colon in inflammatory bowel disease[54] and in tertiary lymphoid structures of salivary glands[56].

### A multicellular community of mmDCs, *IL13*^+^/*IL22*^+^/*IL26*^+^ T cells, NK cells, cycling ILCs and *CCL19*^+^*IL4I1*^+^ fibroblasts in AD may impact KC differentiation

Healthy, non-lesional, and lesional samples separated by their cellular composition (**Figure 6a** and **Figure S13a-g**) in a Principal Component Analysis (PCA), with the second principal component (PC2) capturing variation between lesional and non-lesional/healthy samples, reflecting the joint increased presence of *IL13^+^/IL22^+^/IL26^+^* T cells, mmDCs, *MMP12*^+^ DCs, neutrophils and *CCL19*^+^*IL4I1*^+^ fibroblasts and the strong depletion of cornified KC2 in lesions (**Figure 6b**). PC2 captured further heterogeneity between patient samples, distinguishing those with higher fractions of neutrophils and *IL1B*^+^ DCs (**Figure S13d**), consistent with increased neutrophil numbers observed in AD patients[57] (**Figure 1h**).

**Figure 6.**
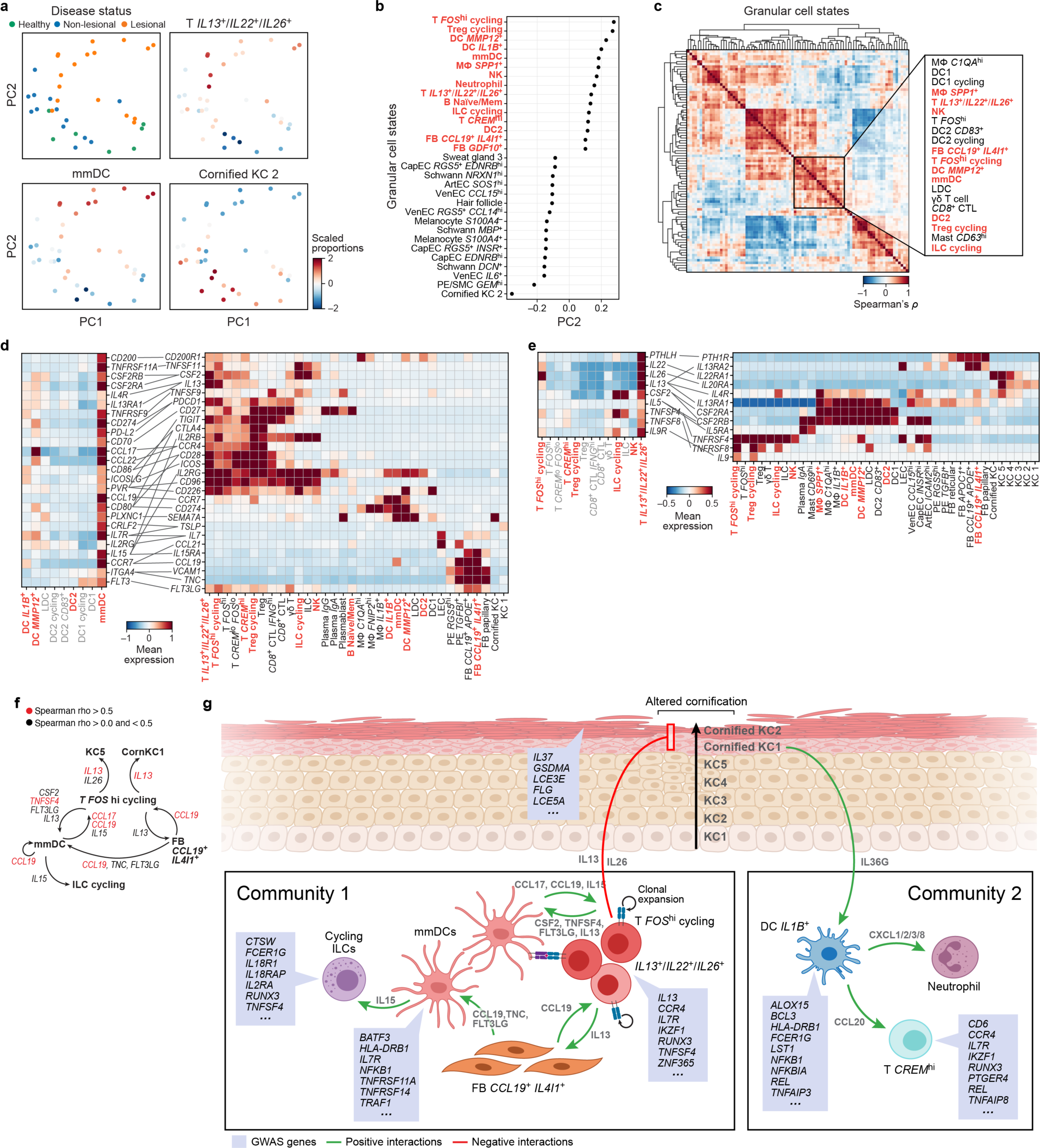
Two multicellular communities associated with disease. **a-c**, A multicellular community associated with lesional AD skin. **a**, 2D embedding of healthy, non-lesional and lesional samples by the first two PCs derived from a PCA of cell composition, colored by disease status (top left) or scaled proportions (z-scores of centered log-ratio (CLR) transformed proportions) of indicated cell subsets. **b**, PC2 loading (x axis) of cell subsets (y axis) with top (red) and bottom (black) 15 loadings. **c**, Correlation coefficient (Spearman’s ρ, colorbar) between proportions of each pair of cell subsets (rows, columns) across samples. Inset: a cluster of *IL13/IL22/IL26* T cells, mmDCs and *CCL19*^+^ *IL4I1*^+^ fibroblasts; red: cell subsets with high PC2 loading. **d,e**, Potential cell-cell interactions between mmDCs and *IL13/IL22/IL26* co-expressing T cells in lesional skin. Mean z-scored expression values (colorbar) of ligand or receptor genes (rows) specifically expressed in mmDCs (d, left, columns) or *IL13/IL22/IL26* T cells (e, left, columns) and their cognate interacting partners (right, rows) across all subsets (right, columns, **Methods**) with a significant interacting partner expression. Edges: ligand-receptor pairs. Red: cell subsets with high PC2 loadings. **f**, Multiple positive feedback loops may initiate or stabilize multicellular community and impact KCs. Cell subsets (nodes) connected from ligand-expressing to receptor expressing subset (edges) where proportion of the receptor-expressing subsets is correlated with ligand expression across samples (see also **Figure S14d**). Red: highly correlated pairs (Spearman’s ρ> 0.5). **g**, A multicellular pathological circuit in AD lesional skin. Schematic showing a model of the interplay between aberrant KC differentiation from KC1 (bottom, epidermis) to cornified KC2 subsets (top, epidermis) and disease-associated multicellular communities with the core of community 1 including clonally expanded *IL13*^+^/*IL22*^+^/*IL26*^+^ T cells, mmDCs, *CCL19*^+^ *IL4I1*^+^ fibroblasts and cycling ILCs as well as the community 2 core of *IL1B*^+^ DCs, T *CREM*^hi^ and neutrophils. The intercellular feedback loops predicted to contribute to community formation are shown (green arrows) and negative impact of community 1 on terminal KC differentiation is indicated (red arrow). GWAS genes specifically enriched in each cell subsets are displayed (blue boxes).

Cell fractions of lesion-enriched immune and stromal cell subsets were highly correlated with each other across samples, identifying several multicellular communities in lesional skin (**Figure 6c** and **Figure S13h,i**), including one community comprising *IL13*^+^/*IL22*^+^/*IL26*^+^ T cells, cycling *FOS*^hi^ T cells, mmDCs, *MMP12*^+^ DCs and *CCL19*^+^*IL4I1*^+^ fibroblasts (Community 1) (**Figure 6c** and **Figure S13h,i**), and another (partly overlapping) with neutrophils, *IL1B*^+^ DCs and *CREM*^hi^ T cell subsets (Community 2) (**Figure S13h,i**).

Community 1 was supported by multiple intra-community receptor-ligand (R-L) interactions that are significantly enriched in lesional AD (**Figure 6d-f** and **Figure S14a**; **Methods; Supplementary Table 12**), with multiple self-reinforcing positive feedback loops, especially between *FOS*^hi^ T cells, mmDCs, ILCs, and *CCL19*^+^*IL41*^+^ fibroblasts, which may contribute to community assembly and stability (**Figure 6f**). For example, mmDCs expressed *CCL19*, *CCL17*, and *IL15*, which recruit and expand T cells and ILC/NK cells, respectively[58], and, in turn, cycling *FOS*^hi^ T cells expressed *TNFSF4*, *FLT3LG*, *CSF2* and *TNFSF11/RANKL*, which are mediators of DC activation, expansion, survival and maturation[59–61] (**Figure 6d-f** and **Figure S14a**). Consistently with this model, there was a positive correlation between the level of *CCL19* and *CCL17* cell intrinsic expression in mmDCs and the *proportion* of cycling *FOS*^hi^ T cells (**Figure 6f** and **Figure S14d**; Spearman’s ρ = 0.84 and 0.76; **Supplementary Table 13**), as well as between *TNFSF4* expression in cycling *FOS*^hi^ T cells and mmDC proportions (**Figure 6f** and **Figure S14d**; ρ = 0.53) (**Methods**,[62]).

Moreover, through extra-community interactions, Community 1 likely impacts other components of the lesional AD skin, especially aberrant KC cornification. T effector cytokines *IL13*, *IL22* and *IL26* displayed putative interactions with KCs via *IL13RA1/IL4R*, *IL22RA1* and *IL20RA*, the latter highly expressed in cornified KCs and KC5, respectively (**Figure 6e**). Moreover, *IL13* expression in cycling *FOS*^hi^ T cells was positively correlated to the proportions of KC5 and cornified KC1 cells (**Figure 6f** and **Figure S14d**; ρ = 0.57 and 0.63), consistent with the known impact of type 2 cytokines on the expression of filaggrin and other cornified envelope components[63]. Together with the observed proportional shifts in those apical KC subsets and an altered KC trajectory in lesional skin (**Figure 2j** and **Figure S8c**), this suggests an impact of KC-community 1 interactions on KC differentiation.

### A cornified KC - *IL1B*^+^ DC - neutrophil - T cell communication axis in AD

Within Community 2, there were multiple putative interactions related to type 17 immunity and neutrophil recruitment (**Figure S14c**), two hallmarks of psoriasis[64]. *IL1B*^+^ DCs had multiple putative interactions with neutrophils via *CXCL1/2/3/8*-*CXCR1/2* and with multiple T cell subsets including *IL13*^+^/*IL22*^+^/*IL26*^+^ T cells and *CREM*^hi^ T and DCs (*e.g.*, mmDCs, *MMP12*^+^ DC) via *CCL20-CCR6* (**Figure S14c**). These putative R-L interactions suggest possible cross-talk between community 2 and community 1 members, also reflected by partial overlap in the co-emergence of both communities (**Figure S13h** and **Figure S14d**), with *SPP1*^+^ macrophages, *IL13*^+^/*IL22*^+^/*IL26*^+^ T cells, and NK cells as shared components (**Figure S13h**). Both CXCL1/2/3/8-CXCR1/2 and CCL20-CCR6 have been implicated in mediating Th17 cell recruitment and polarization in psoriasis[64]. Consistently, *CCL20* expression in *IL1B*^+^ DCs correlated positively with *CREM*^hi^ T cell proportions in lesional skin (**Supplementary Table 13;** ρ = 0.56).

Community 2 might be impacted by cornified KCs, which expressed high levels of *IL1* family cytokines and inhibitors and up-regulated the psoriasis disease genes *IL36G*, *IL36B* and *IL36RN*[65] (**Figure S14b**) in cornified KC1 in lesional skin (**Figure 2k**). The IL36 receptor (*IL1RL2*) was mainly detected in KCs, LDCs and *IL1B*^+^ DC, indicating intraepidermal signaling and cornified KC-DC crosstalk via IL36 (**Figure S14b**)[64]. Thus, a cornified KC - *IL1B*^+^ DCs - neutrophils - T cell communication axis may operate in inflammatory skin disease. Community 2 is detectable only in a subset of patients (**Figure S13b-d**), possibly reflecting proposed patient heterogeneity[57,66].

### GWAS-nominated AD genes are enriched within the lesional multicellular community including *IL13*^+^/*IL22*^+^/*IL26*^+^ T cells and in cornified keratinocytes

To relate our findings to AD genetic etiology, we examined genes associated with AD risk by GWAS to the cell subsets in our skin cell atlas. We obtained disease-associated genes from a GWAS compendium of genome-wide significant genetic loci (2,678 studies, including 8 AD studies) using the OpenTargets Genetics Locus2Gene mapping[67] and tested for their enrichment in each of the 86 granular cell subset signatures as well as all lymphocyte gene programs (topics) (**Methods**).

Among cell subsets, AD GWAS gene expression was significantly enriched in *IL13*^+^/*IL22*^+^/*IL26*^+^ T cells (Log OR: 3.43, FDR<10^-9^, Fisher’s exact test), cornified keratinocytes (Log OR: 3.25, FDR<0.01), ILCs (Log OR: 2.96, FDR<10^-6^) and overall members of multicellular Community 1 (P value < 0.01, Fisher’s exact test; **Figure S15a, red**; **Supplementary Table 14**). Consistently, among lymphocyte gene programs, topic #190-positive cells were the most enriched for AD GWAS genes (**Figure S15c**), as were topic #3 (with key TF regulators of Th2 fate *BHLHE40*[48] and *NFIL3*[68]) and #171 (with GWAS genes *IKZF3* and *TSHZ2*, implicated in T cell differentiation) (**Figure S15c,d**)[69]).

AD risk genes driving the cornified keratinocyte enrichment included *FLG*2 located in the epidermal differentiation complex locus, *IL37*, an anti-inflammatory cytokine, and Gasdermin A (*GSDMA*), a member of a protein family involved in cell death and inflammation (**Figure S15e**). The *IL13*^+^/*IL22*^+^/*IL26*^+^ T cell enrichment was driven by *IL13, CCR4, IL7R, IKZF1, RUNX3, CD274, CD6* and *THEMIS* (**Figure S15f**).

Finally, to highlight possible distinct and shared disease mechanisms, we expanded the scope of our enrichment analysis across GWAS of multiple skin and type 2 diseases (**Figure S15b; Supplementary Table 14**). Cornified KCs were associated with GWAS genes for atopic dermatitis, psoriasis and atopic asthma, the latter ranking highest across tested diseases together with AD (**Figure S15b** and **Figure S15g**). Asthma, AD and allergic rhinitis were also among the top diseases displaying *IL13*^+^/*IL22*^+^/*IL26*^+^ T cell-associated GWAS signal (**Figure S15h**), further supporting common genetic susceptibility and underlying mechanism of these type 2 diseases[70].

## Discussion

In this study, we mapped the cellular ecosystems of healthy and inflamed skin in AD by combined scRNA-seq with TCR-seq and scATAC-seq. By establishing a rapid and efficient whole skin tissue processing workflow without epidermal-dermal tissue layer separation, lengthy dissociation, or marker-based flow cytometry enrichment, we constructed a skin atlas that more comprehensively captures the range of cells in skin tissue, especially rare or difficult-to-profile subsets, such as cornified keratinocytes. Through computational analysis of the atlas in the context of disease we tie together pathologies at the genetic (risk genes), cellular and gene program, cell differentiation, and cell community levels, thus providing a tissue biology perspective to skin disease.

We found strong support for a unified multi cellular model of disease, where lesional AD skin has an expanded immune compartment with the co-emergence of two multicellular immune/stromal communities – one present in all patients and another present only in a subset of patients. These are connected to a pathological KC differentiation trajectory with increase in KC5 and early cornified cells (cKC1s) at the expense of terminally cornified cells (cKC2s), such that one community may help drive pathological KC differentiation and the other may be driven by it (**Figure 6g**). GWAS genes are particularly enriched in many of those components, closely tying the multicellular skin pathological state to the genetics driver of the disease (**Figure 6g**).

Disease-associated Community 1 included clonally expanded *CD4*^+^ and *CD8*^+^ *IL13*^+^/*IL22*^+^/*IL26*^+^ and cycling *FOS*^hi^ T cells with overlapping type 2, 22 and 17 characteristics, *MMP12*^+^ DCs, *CCL17*^hi^*CCL22*^hi^ mregDC/mmDCs, *CCL19*^+^ *IL4I1*^+^ fibroblasts, and cycling ILCs, with multiple putative intercellular positive feedback loops that might contribute to its formation, stability and maintenance (**Figure 6g**). At the core of Community 1, cycling *FOS*^hi^ and *IL13*^+^/*IL22*^+^/*IL26*^+^ T cells may be both recruited via *CCR4*-*CCL17/CCL22* and activated via *IL15* by mmDCs/mregDCs. *IL13*^+^/*IL22*^+^/*IL26*^+^ T cells profiles resembled those of Th2A cells and of disease-associated T cells in eosinophilic esophagitis[71] and asthma[72], suggesting a common pathogenic state across type 2 diseases. Antigen-loaded mmDCs/mregDCs may further impact these T cells in a TCR-dependent manner and we hypothesize that antigen stimulation may contribute to the observed clonal expansion and increased cycling of *IL13*^+^/*IL22*^+^/*IL26*^+^ T cells (**Figure 4l,m** and **Figure 4q-r**). Indeed, in the context of non-small cell lung cancer, antigen loading and presentation by mregDCs has been demonstrated and antigen uptake has been suggested to be a signal for mregDC generation[36]. Further studies to test if ongoing antigen stimulation in lesional skin tissue is a contributor to the chronic inflammatory response can leverage TCR sequences from clonally-expanded, pathology-associated *IL13*^+^/*IL22*^+^/*IL26*^+^ T cells (**Figure 4o**) to characterize recognized antigens and elucidate mechanisms of antigen sensitization in the course of disease and therapy. Upon activation, *IL13*^+^/*IL22*^+^/*IL26*^+^ T cells, are in turn predicted to impact DCs, including mmDCs/mregDCs and *MMP12*^+^ DCs via myeloid cell activation and expansion factors *GM-CSF, FLT3LG* and *TNFSF4/OX40L* as well as *IL13* (**Figure 6f,g**). This is consistent with the increase in STAT6 inferred activity in lesional *MMP12*^+^ DCs and mmDCs/mregDCs, induction of IL13-STAT6 target genes (*e.g.*, *CCL17, IL4I1*) (**Figure 3d** and **Figure S10k**), and shift in DC differentiation/activation trajectories to a partially activated *MMP12*^+^ DC state in non-lesional skin and a fully activated mmDC/mregDC state in AD lesions (**Figure S10l**). This fully activated mmDC/mregDCs state expresses both *CCL19* and its receptor *CCR7*, which further may result in the attraction and clustering of mmDCs/mregDCs in DC-T rich areas[64,73].

Community 1 may play a key role in the pathological KC differentiation and cornification trajectory observed in AD lesional skin, with a shift towards a less mature cornified KC1 subset, with lower expression of late cornified envelope constituents, and accumulation of non-fully differentiated KC5s (preceding cKC1s). This connection is supported by a significant correlation between the cell intrinsic expression of key cytokines *IL13* and *IL26* in T cells and the proportion of KC5s and cornified KC1s (**Figure 6f**) expressing their cognate receptors. This supports a model where IL13 directly affects terminal KC differentiation, consistent with reports of disrupted cornified gene expression in IL13-treated KCs *in vitro*[63]. Restoring the transition from cornified KC1s to cornified KC2s in AD might present an avenue for therapeutic barrier restoration.

In a subset of AD patients, we observed a second AD-associated Community 2, with co-emergence of *IL1B*^+^ DCs, *CREM*^hi^ T cells and neutrophils (**Figure 6g**). *IL1B*^+^ DCs described here displayed striking similarity to the recently described *IL23A*^+^*IL1B*^+^ DC3 subset, which was shown to be prevalent in psoriasis[35]. *IL1B*^+^ DCs formed multiple putative interactions with neutrophils and expressed the Th17 regulators *CCL20* and *IL23A*. Varying amounts of skin-infiltrating neutrophils and type 17 expression signatures have been reported in previous AD patient cohorts, suggesting the presence of multiple AD endotypes[57,66]. Future studies in larger patient cohorts will further help stratify patient groups and will be able to corroborate patient heterogeneity in respect to Community 2. One intriguing possibility is that Community 2’s formation may be impacted by aberrant KC differentiation, because cornified KCs, especially in lesional skin, express high levels of *IL1* family cytokines and inhibitors, including up-regulation of the psoriasis disease genes *IL36G*, *IL36B* and *IL36RN*[65], and the IL36 receptor (*IL1RL2*) was expressed in detected in *IL1B*^+^ DC (**Figure S14b**).

The cell types and programs in the pathological circuit in lesional AD skin may be formed or driven by genes associated with AD through GWAS. There is a significant enrichment of GWAS genes in cornified KCs, *IL13*^+^/*IL22*^+^/*IL26*^+^ T cells, and multiple other Community 1 and 2 cells, as well as in Community 1 cells as a whole (but not Community 2 as a whole). This agrees with a model of multiple routes to disease, where either epithelial or immune dysfunction and their interactions can initiate and then converge towards AD, and where genetic variants in multiple loci may affect the pathological formation of Community 1 and its interaction with skin cornification through multiple feedback loops (**Figure 6g**). Importantly, multiple key GWAS genes, such as *TNFSF4/OX40L* and *IL13* are part of the predicted intercellular feedback loops that could contribute to Community 1 formation and maintenance and its interactions with KCs and their differentiation (**Figure 6g**).

Together, our multi-modal fine-grained skin census provides insight into the remodeling of human skin under chronic inflammation, nominates targetable cellular states and transitions and serves as a fundamental resource towards a deeper understanding of skin biology.

## Acknowledgements

We thank members of the Regev, Xavier and Geha laboratories for helpful discussions. We thank L. Gaffney and A. Hupalowska for help with figure preparation and E. Heppenheimer for critical reading of the manuscript. This research was supported by NHGRI grants 5RM1 HG006193 (Center for Cell Circuits, to A.R.), Klarman Cell Observatory (A.R., R.J.X), DK043351 and DK114784 (to R.J.X), NIAID/NIH grant 1UM1AI151958 (to R.S.G.), and a gift from the Food Allergy Science Initiative (to A.R. and R.J.X), a gift from the Manton Foundation (to A.R. and R.J.X), and HHMI (to A.R.). E.F. is supported by an EMBO Long-Term fellowship, J.M.L-C. by NIAID T32 grant (5T32AI007512-32), NIH award (UL1 TR002541) and a Dermatology Foundation Career Development Award. C.A.L. is supported by a Stanford Science Fellowship. A.R. was a Howard Hughes Medical Institute Investigator (until July 31, 2020). This paper is part of the Human Cell Atlas.

## Author Contributions

E.F., G.E., O.R.-R., R.S.G., R.J.X. and A.R. conceived and designed the project. E.F. designed and performed experiments E.F. and G.E. analyzed and interpreted the data, with guidance from A.R. G.E. designed and led data analysis, with guidance from A.R. M.B.A-P. performed clinical work. C.A.L. contributed ATAC-seq data analysis. E.F., M.B.A-P, J.M.L-C. collected biopsies, E.P.F. and I.T.-S. performed additional tissue processing and scRNA-seq experiments. E.F., S.I. performed mouse MC903 experiments. M.S., J.W. and T.M.D. provided experimental assistance. H.C., S.K., H.L., B.L., A.C., B.W., W.P., L.S., C.P., and J.C. assisted in patient recruitment and biopsy collection. G.P.S., T.M.D., J.D., O.R.-R., R.S.G., R.J.X. and A.R. provided project oversight and acquired funding. E.F., G.E., R.J.X. and A.R. wrote the manuscript with input from all authors.

## Supplementary Tables

Supplementary Table 1: Cell subset markers

Supplementary Table 2: Coarse and granular proportion analysis

Supplementary Table 3: GO enrichment results

Supplementary Table 4: Differential gene expression analysis

Supplementary Table 5: Classifier markers

Supplementary Table 6: Classification concordance

Supplementary Table 7: Differential TF activity in dendritic cells

Supplementary Table 8: mmDC atlas meta-data

Supplementary Table 9: Lymphocyte topic model gene loadings and differential topic expression results

Supplementary Table 10: VDJ analysis

Supplementary Table 11: Differential gene expression of expanded vs non-expanded T cells

Supplementary Table 12: Receptor-ligand analysis

Supplementary Table 13: Ligand expression and receptor cell subset correlations

Supplementary Table 14: GWAS enrichment results

## Supplementary Figure legends

**Figure S1. Overview of current patient cohort**

Assessed disease severity, biopsy locations and co-morbidities are noted.

**Figure S2. Quality control of scRNA-seq skin cell atlas**

**a-c**, Quality metrics. Number of recovered cell barcodes passing quality control and filtering steps (a, y axis), distribution of the number of genes per cell barcode (b, y axis), and distribution of fraction of detected mitochondrial genes per cell barcode (c, y axis) in each processed sample (x axis) from healthy (green), non-lesional (blue) and lesional (orange) skin. White dots: median. **d,e**, Individual mixing and marker expression. 2D embedding of scRNA-seq profiles from the entire cohort colored by profiled individuals (d) or expression of coarse cell type markers (e, log(TP10K+1), color bar).

**Figure S3. Coarse and fine cell subsets in skin and skin appendages**

**a**, Coarse and fine cell subsets in skin. 2D embedding of epithelial (left), immune (middle) or stromal (right) scRNA-seq profiles from the entire cohort, colored by annotated coarse (top) or granular (middle) cell subsets or disease status (bottom). **b-e**, Rare hair follicle, sweat and sebaceous gland cell subsets. **b,d**, 2D embedding of scRNA-seq profiles from the entire cohort, colored by annotated granular cell subsets (b) or by z-score expression values of cell subset marker genes (d). **c**, Mean expression (dot color) and fraction of expressing cells (dot size) of hair follicle and gland subset marker genes (columns) across cell subsets (rows). **e**, Immuno-histochemistry image of sweat gland marker Dermicidin (DCD) in a healthy skin tissue section derived from the Human Protein Atlas[74]. Scale bar: 200µm. **f,g**, All lesional samples display higher expression of inflammatory epidermal markers. Distribution of the expression of *S100A8* (f, y axis) or *KRT16* (g, y axis) across each processed 10x Chromium channel (x axis). White dots: median.

**Figure S4. Integrated skin cell atlas**

**a,b**, Consistent cell subset annotations across datasets. Agreement (color, row-normalized) between coarse (a, columns) or granular (b, columns) cell subsets in the current study and annotations from each of four published skin scRNA-seq dataset[8,10–12] (rows) based on a linear classifier trained on the current study. **c-f**, Integrated skin atlas of current and published skin scRNA-seq datasets. 2D UMAP embedding of integrated scRNA-seq profiles (**Methods**) from the current cohort and four published datasets[8,10–12], colored by study (c), coarse cell types (d), granular cell subsets (e), or disease status (f). **g**, Cell compartment representation across studies. Proportion (x axis) of cells from each major compartment (color of bars) in each dataset (y axis). **h-j**, Cornified keratinocytes and other cell subsets absent from published data sets. **h,j**, Proportion (%) (color) of annotated coarse (h) or granular (j) subsets (columns) in the current and published studies (rows). **i**, Number (x axis) of profiled cornified keratinocytes (left) or cells overall (right) in the current and published studies (y axis). Color: condition.

**Figure S5. Changes in broad cell type proportions between healthy, non lesional and lesional skin**

**a,b**, Variation in cell proportions across individuals and conditions. **a**, Proportion (x axis) of each coarse cell types (color) in each processed 10x Chromium channel (rows, ordered from highest to lowest immune cell proportions within each condition) in each condition (right). **b**, Proportion (x axis) of coarse cell types (rows, ordered by compartment) in healthy, non-lesional and lesional conditions in each individual (color). Left bar: total number of cells profiled of each coarse type. **c**, Differences in coarse cell types between conditions. Fraction of cells (y axis) in each condition (x axis) for each coarse category (panels). Center line: median; box limits: first and third quartiles; whiskers: 1.5× interquartile range.

**Figure S6. Keratinocyte differentiation continuum in adult skin**

**a-c**, Identification of cycling keratinocytes. **a**, 2D embedding of keratinocyte scRNA-seq profiles colored by cell cycle phase (**Methods**). Arrow: differentiation axis from basal to terminal KCs. **b,c**, Distribution of G2/M (b) and S (c) phase scores (y axis) in KC subsets (x axis). **d**, Higher expression of late cornified envelope genes in cornified KC2s. Mean z-scored expression values (color) of genes (columns) differentially expressed between cornified KC1s and KC2s (rows). BH FDR: **<0.01, ***<0.001 (Welch’s t test). **e**, Cornification trajectories in healthy, non-lesional and lesional skin. RNA velocities (arrows) projected onto 2D UMAP embedding of scRNA-seq profiles of KC5s, cornified KC1s and cornified KC2s in healthy, non-lesional and lesional skin colored by cell subset (left) or disease status (right). **f**, Enriched pathways and processes in different KC subsets recapitulate human skin biology. Reactome[75] terms (rows) that are significantly enriched (x axis, -log10(BH FDR), Fisher’s exact test, **Methods**) in each KC subset. **g-i**, TF and monogenic skin disease temporal gene expression along KC differentiation follows function and phenotype. **g,i**, Scaled expression values (color) of TF encoding genes (g, rows) or monogenic skin disease genes from OMIM (i, rows) in cells (g,i, columns) from healthy controls skin ordered along DPT representing keratinocyte differentiation from basal (left) to terminal cornified (right) cells. Colored bars: DPT (top), G2/M score (middle) and annotated KC subset (bottom). Data were downsampled (**Methods**) and equal numbers of cells for each KC subset are shown. **h**, Predicted mean TF activity (color) (**Methods**) for each TF gene in (g) across KC subsets (columns) from basal (left) to terminal (right).

**Figure S7. Single-cell chromatin accessibility in human skin and KC differentiation**

**a,b**, QC metrics of skin scATAC-seq. **a**, TSS enrichment score (y axis) and number of unique chromatin fragments (x axis, log10 scale) for each cell barcode (dots) passing filtering (dashed lines) (**Methods**), colored by density. Top right: median number of fragments and median TSS scores. **b**, Frequency distribution of size (x axis, base pairs) of detected chromatin fragments. **c-f**, KC differentiation in scATAC-Seq profiles. 2D embedding of scATAC-seq profiles from patient skin biopsy, colored by annotated coarse cell types (c), gene activity scores of cell type (d) or KC subset (f), or by DPT (e). **g**, Change in inferred TF activity along KC differentiation. ScATAC-seq gene activity scores (color) of TF encoding genes (rows) in cells (columns) ordered by KC DPT (top bar). Arrow: KC differentiation from basal (left) to terminal (right).

**Figure S8. Remodeling of KC expression and cornification in lesional AD skin**

**a**, Changes in KC subset composition. Proportions (y axis) of KC subsets (x axis) in each disease condition (colored bars). Center line: median; box limits: first and third quartiles; whiskers: 1.5× interquartile range. **b**, Near absence of KC2s in lesional AD skin. 2D diffusion map of KC scRNA-seq profiles from healthy (left), non-lesional (middle) and lesional (right) skin colored by KC subset. Arrows: differentiation trajectory from basal to terminal cornified KCs. **c**, Expression changes in cornified KCs and in bulk RNA-seq. Mean z-scored expression values (color) of genes (columns) differentially expressed (by scRNA-seq) in cornified keratinocytes between conditions (rows) in scRNA-seq profiles (top) and in bulk RNA-seq[16] (bottom). BH FDR: #<0.1, *<0.05, **<0.01, ***<0.001. **d**, Enriched pathways and processes in KC subsets in AD lesional skin. Top 10 most significant Reactome[75] terms (rows) (x axis, -log10(BH FDR), Fisher’s exact test, DEGs with FDR < 0.1, |log2FC|>0.5 were used) in each KC subset (colors) (**STAR Methods**). **e**, Fraction of keratinocytes (y axis) expressing *IL19* (left) and *IL24* (right) in healthy, non-lesional and lesional conditions (x-axis). **f,g**, Decreased abundance of cornified KC2s and increased abundance of cycling KC2s are features of lesional atopic dermatitis skin but not observed in scleroderma scRNA-seq. **f**, 2D embedding of scRNA-seq profiles from 7 healthy donors and two scleroderma patients, colored by annotated coarse cell types. **g**, Proportion of cornified KC1s and KC2s (y axis) in healthy, scleroderma and AD skin (columns).

**Figure S9. The MC903 mouse model of skin inflammation recapitulates aspects of human atopic dermatitis within the KC compartment**

**a-c**, Mouse skin scRNA-seq in uninflamed and MC903-induced inflammation. **a**, Experiment overview (**Methods**). **b,c**, 2D embedding of scRNA-seq profiles from control and MC903-treated mice, colored by annotated coarse cell types (b) or by treatment (c). **d**, Mean expression (dot color) and proportion of expressing cells (dot size) of coarse cell type marker genes (columns) in each coarse cell type (rows). **e**, Conserved KC differentiation trajectory across species. 2D embedding of KC scRNA-seq profiles from control mice, colored by z-scored expression values of human KC subset markers. **f-i**, MC903-induced inflammation mimics multiple cell composition and expression changes of human AD skin. **f**, Proportion (y axis) of cells of each coarse type (color) across conditions (columns). **g**, Infiltration of neutrophils and basophils and near absence of cornified KCs and in MC903-treated skin. Proportion of cells (y axis) from untreated and treated mice (color) in each coarse cell type (x axis). **h,i**, Increased expression of epidermal inflammation markers and KC cycling, and decrease of late cornified envelope transcripts. Proportion (y axis) of KCs expressing specific marker genes in control and MC903-treated mice (color). **j**, Mean expression (dot color; min-max scaled log(TP10K+1)) and fraction of expressing cells (dot size) of *Il13* and *Il4* (columns) in broad cell types in control and MC903-treated skin (rows).

**Figure S10. A shift in DC activation trajectories towards a mature migratory state in lesional AD skin**

**a-c**, mmDCs express a migratory expression signature. **a**, Mean expression (dot color) and fraction of expressing cells (dot size) of actin cytoskeleton or cell migration genes (rows) in myeloid cell subsets (columns). **b,c**, 2D embedding of DC scRNA-seq profiles from healthy and AD skin, colored by expression of marker genes (b, z score) or by disease condition (c). **d**, DC activation trajectory. Expression of DC subset marker genes (rows) in cells (columns) from healthy, non-lesional or lesional skin ordered by DPT (top color bar) along DC activation trajectory and labeled by subset (bottom color bar, as in **Figure 3d**). Data were downsampled (**Methods**) and equal numbers of cells for each cell subset are shown. **e**, Key TF regulators of DCs. Mean expression (dot color) and fraction of expressing cells (dot size) of key DC TF genes (columns) in each DC subset (rows). **f,g**, Changes in myeloid cell composition across conditions. Proportion of cells (y axis) in different DC (f) and macrophage (g) subsets in each condition (x axis). **h-j**, Increased DC *MMP12*^+^ cells in AD. **h,i**, Proportion of *MMP12*^+^ DC within DCs (y axis) in each condition (x axis) in published scRNA-Seq (h[8], and i[11]). **j**, Mean expression (color, z-score) of mmDC and *MMP12*^+^ DC marker genes (columns) across conditions (rows) in bulk RNA-Seq[16]. **k**, High inferred STAT6 activity in *MMP12*^+^ DCs and mmDCs. Differentially TF activity scores (y axis; lesional - healthy) for each TF in each myeloid cell subset (dot) sorted by TF rank (x axis) (**Methods**). **l**, Aberrant DC differentiation trajectories in AD skin. Distribution of diffusion pseudotimes (DPT, x axis, **Methods**) for DC2s (top), *MMP12*^+^ DCs (middle) and mmDCs (bottom) in each condition (color). *BH FDR<0.1 ***BH FDR<0.001 (Mann-Whitney U test). Vertical dashed lines: median. Boxplots for **f-i**: center line, median; box limits, first and third quartiles; whiskers, 1.5× interquartile range.

**Figure S11. Characterization of skin lymphocytes and expanded effector IL13/IL22/IL26 co-expressing T cells**

**a-e**, Lymphocyte subsets in healthy and inflamed human skin tissue. **a-d**, 2D embedding of scRNA-seq profiles of lymphocytes from healthy and AD skin, colored by expression of indicated marker genes (a, z score), cell cycle phase (b) or *CD4*/*CD8* status (d). **c**, Cycling lymphocyte subsets. Proportion (x axis) of lymphocytes in each cell cycle phase (color, as in **b**) from each subset (rows). **e**, *CD4*/*CD8* T cell subsets. Proportion (x axis) of *CD4*^+^ or *CD8*^+^ cells (color, as in D) in each T cell subset (rows). **f**, Lymphocyte gene programs. Probabilities (x axis) of top scoring genes (y axis) in each topic. **g-h**, Co-expression of *IL13, IL22* and *IL26* across lymphocytes in lesional skin. **g**, Pearson’s correlation coefficient (r, x axis) for the top genes (y axis) correlated with *IL22, IL26, IL5* or *IL31* across lymphocytes, rank ordered by correlation. **h**, 2D embedding of lymphocyte scRNA-seq profiles from healthy and AD skin, colored expression of *IL13, IL22* or *IL26* (z score). **i,j**, Distribution of *IL13^+^/IL22^+^/IL26^+^*co-expressing T cells across lymphocyte subsets. Proportion (x axis) of *IL13^+^/IL22^+^/IL26^+^*co-expressing T cells (“yes”) and all other T cells (“no”) from each lymphocyte subset (i, color) or from *CD4*^+^, *CD8*^+^, and *CD4^+^*/*CD8^+^*cells (j, color). **k**, Lymphocyte subsets that do not change in proportion between conditions. Proportions (y axis) of lymphocyte subsets (x axis) in each condition (*x* axis, color). Center line: median; box limits: first and third quartiles; whiskers: 1.5× interquartile range. **l**, *IL13^+^/IL22^+^/IL26^+^*expressing T cells are detected in lesional AD but not scleroderma skin. Fraction of lymphocytes (y axis) expressing *IFNG*, *IL13*, *IL22*, or *IL26* in each condition (*x* axis, color). **m-p**, T cell clonality analysis. **m**, Clone size (y-axis) for each T cell clone detected in two patients (color). **n**, 2D embedding of lymphocyte 5’ scRNA-seq profiles from lesional skin colored by expansion status. **o,p**, Proportion (x axis) of lymphocytes of each subset (o, as in **i**) or of CD4 or CD8 (p) in expanded (“>=2”) or non-expanded (“1”) T cells (rows). **q**, Expanded *Il22/IL13* co-expressing T cells. *IL22* (y axis) and *IL13* (x axis) expression in T cells from expanded (red) and non-expanded (grey) clones.

**Figure S12. Stromal cell diversity in healthy and AD skin**

**a,b**, Pericyte and vascular smooth muscle cell (SMC) subsets. **a**, 2D embedding of scRNA-seq profiles of pericytes and SMCs from healthy and AD skin, colored by subset. **b**, Mean expression (dot color) and fraction of expressing cells (dot size) of pericyte and SMC subset marker genes (columns) across cell subsets (rows). **c,d**, Fibroblast subsets. 2D embedding of scRNA-seq profiles of fibroblasts from healthy and AD skin, colored by their proposed dermal location[76,77] (c) or by marker gene expression (d, z-score). **e**, Distinct functional processes in fibroblast subsets. Reactome[75] terms (rows) enriched (x axis, -log10(BH FDR), Fisher’s exact test) in each fibroblast subset (**STAR Methods**). **f,g**, Schwann-like fibroblast subsets. **f**, 2D embedding of scRNA-seq profiles of Schwann-like fibroblasts and Schwann cells from healthy and AD skin, colored by coarse (left) or granular (right) subset. **g**, Mean expression (dot color) and proportion of expressing cells (dot size) of Schwann-like fibroblast and Schwann cell subset marker genes (columns) in each cell subset (rows). **h-j**, Cell subsets that are largely unchanged across conditions. Cell proportions (y axis) of different subsets of fibroblasts (h), endothelial cells (i) or pericyte/SMCs (j) across conditions (color). **k**, Increased abundance of classified *CCL19*^+^ *IL4I1*^+^ fibroblasts in lesional AD skin across studies. Proportion (y axis) of *CCL19*^+^ *IL4I1*^+^ fibroblasts annotated by our classifier (**Methods**) in published scRNA-seq studies (k, left[8] and middle[11]) and deconvoluted bulk RNA-seq (k, right[16]).

**Figure S13. Multicellular communities in lesional AD skin**

, Sample characterization by cell composition. **a**, Scaled proportions (z-scores of centered log ratio (CLR) transformed proportions, color) of granular cell subsets (rows) across samples (columns). Red: cell subsets with high PC2 loading (**Figure 6b**). **b-g**, 2D embedding of healthy, non-lesional and lesional samples along the first two PCs from PCA of cell composition, colored by disease status (b, left), individual (b, right), mean expression of effector T cell cytokines (c) or epidermal inflammation markers (e and f) or scaled cell subset proportions (d and g). **h**, Correlation coefficient (Spearman’s ρ, colorbar) between proportions of each pair of 86 granular cell subsets (rows, columns) across samples (larger version of **Figure 6c**). Red: cell subsets with high PC2 loading. **i**, Selected correlated cell subsets from Community 1. Proportion of cells (x and y axis) of selected pairs of subsets in each sample from healthy (green), non-lesional (blue) or lesional (orange) skin. Top left: Spearman’s ρ.

**Figure S14. Cell-cell interactions in AD**

**a-c**, Cell-cell interactions formed by cycling ILCs, cornified KCs and *IL1B*^+^ DCs. Mean z-scored expression values (color) of ligand and receptor genes of cycling ILCs (a), cornified KCs (b) or *IL1B*^+^ DCs (c) across related cell subsets (top, rows) and their cognate interacting partners (bottom, columns) across all cell subsets with significant interacting partner expression (bottom, rows) (**Methods**). Lines: interacting ligand-receptor pairs. Red: cell subsets with high PC2 loading (**Figure 6b**). **d**, Relation between cellular communities. 2D UMAP embedding of granular cell subset proportion profiles (dots) across all samples colored by proportion log odds (top) and community membership (bottom). Positive log odds: enriched in lesional. **e**, Correlation between ligand expression and proportion of cells expressing a cognate receptor in Community 1. Mean expression level of a ligand in one cell subset (x axis) and proportion of a cell subset expressing the cognate receptor (y axis) in each lesional sample. Dashed line: best linear fit. Top left: Spearman’s ρ. **f-h**, Putative interactions with peripheral neurons. **f**, 2D embeddings of scRNA-seq profiles of all cells from the current study and from a published non-human primate dorsal root ganglion (DRG) scRNA-seq dataset[49] (top) (**Methods**), and from the DRG neurons only (bottom, inset), colored by cell types. **g**, Mean expression (dot color) and fraction of expressing cells (dot color) of peripheral neuron marker genes (columns) across cell subsets (rows). **h**, Mean expression (color, z-score) of neuron-specific ligand and receptor genes (left, rows) across DRG neuron subsets (left, columns) and of their cognate interacting genes (right, rows) in each skin cell subset with significant expression (right, columns). Lines: ligand-receptor pairs. Red: cell subsets with high PC2 loading (**Figure 6b**).

**Figure S15. GWAS gene enrichment in skin cell subsets**

**a**, *IL13^+^/IL22^+^/IL26^+^* T cells and cornified KCs are enriched in expression of AD GWAS genes. Skin cell subsets (rows) enriched (-log10(FDR), Fisher’s exact test, dot color) for expressing AD GWAS nominated genes, sorted by odds ratio (x axis, log2 scale). Red: Cells from Community 1. Blue: Cells from Community 2. **b**, Relation of skin cell subsets to skin and inflammatory diseases. Enrichment (-log10(FDR), Fisher’s exact test, dot color) and odds ratio (log2, dot size) of GWAS nominated gene sets for inflammatory and skin-related traits (rows) in genes expressed in each skin cell subset (columns) (**Methods**). **c,d**, Lymphocyte gene program enriched in AD GWAS genes. **c**, Topics (rows) enriched (-log10(BH FDR), Fisher’s exact test, x axis) for expressing AD GWAS nominated genes, sorted by -log10(BH FDR) and colored by log odds ratio. **d**, Probabilities (y axis) of each of the top scoring genes (y axis) in each of the top five GWAS enriched topics (after topic 190) (**Methods**). **e,f**, Selected AD GWAS-nominated genes expressed in skin cell subsets. Mean expression (color, z-score) of cornified KC-specific (e) or lymphocyte-specific (f) AD GWAS genes (rows) in coarse cell types (rows). **g,h**, Enrichment of GWAS genes from various atopic and type 2 inflammatory diseases in cornified KCs and *IL13^+^/IL22^+^/IL26^+^*co-expressing T cells. Significance (-log10(BH FDR), x axis) and effect size (log2(odds ratio), dot color) of enrichment of GWAS nominated gene sets for different traits (rows) enriched in cornified KCs (g) or *IL13^+^/IL22^+^/IL26^+^* co-expressing T cells (h) (**Methods**).

## Methods

### Code availability

Code for all analyses is available on GitHub: https://github.com/klarman-cell-observatory/human_skin_inflammation.

### Patient samples

Whole skin punch biopsy samples of 3-4mm diameter were obtained from gender-matched atopic dermatitis patients (AD) patients, scleroderma patients, and healthy individuals after informed consent and approval to the 2018P002325 and 2017P000025/PHS studies at Massachusetts General Hospital and Boston Children’s Hospital. Clinical information and metadata for the samples are provided in **Figure S1**. Healthy controls were individuals without a history of AD. Patients were included based on a clinical diagnosis of AD and were observed to have active disease via macroscopic assessment from a physician. A non-lesional and lesional biopsy were collected. Skin punch biopsies were immediately placed into tubes with cold DMEM containing 10% FBS, 2 mM L-glutamine and 100 U/ml penicillin/streptomycin (Thermo Scientific) and kept on ice until enzymatic dissociation.

### Enzymatic dissociation of human whole skin punch biopsies

Whole skin punch biopsies (3-4mm) were cut into two pieces, such that each piece contains all skin layers and storage medium was removed. Enzymatic dissociation was performed using Whole Skin Dissociation kit (Miltenyi Biotec, #130-101-540) with a modified protocol. Cut biopsies were transferred into a gentleMACS C-tube containing 435µl Buffer L, 12.5µl Enzyme P, 50µl Enzyme D and 2.5µl Enzyme A. Enzymatic dissociation was performed for 3.5h at 37°C. After incubation, 500µl of cold DMEM containing 10% FBS and 2 mM L-glutamine were added. The tissue was mechanically dissociated using a gentleMACS dissociator run on program h_skin_01. The resulting cellular suspension was filtered through a 70µm cell strainer (Falcon, #352350) and 4.5ml cold DMEM containing 10% FBS and 2 mM L-glutamine were added. Cells were spun down at 300g for 10min at 4°C. The supernatant was then aspirated and cells were resuspended in 1.2ml ACK buffer (A10492-01). Suspensions were transferred to 1.5ml tubes and incubated for 2min for red blood cell lysis and spun at 350g for 4min at 4°C. Supernatants were aspirated and cells were resuspended in 50-100µl 0.4% BSA in PBS. Cell viability and cell counts were determined and cells were processed for scRNA-seq. For scATAC-seq, cells were fixed in PBS containing 1% formaldehyde (ThermoFisher Scientific) for 10min at room temperature. Fixation was quenched by addition of 2.5M glycine to a final concentration of 0.125M and cells were washed in PBS and processed for lysis and tagmentation.

### Mouse model of AD-like inflammation using topical MC903 application

9 week old female C57BL/6J mice were obtained from Jackson Laboratories (Bar Harbor, ME) and maintained in specific pathogen–free conditions in the animal facility of the Massachusetts General Hospital. All experiments were approved by the Massachusetts General Hospital Subcommittee on Research Animal Care (Protocol number 2016N000203). A 2 cm x 2 cm patch of dorsal skin was carefully shaved 2 days before the initiation of treatment. 2 nM MC903 (R&D, Tocris, MN, USA) dissolved in ethanol or ethanol as control were topically applied daily for a period of 14 days. Mice were euthanized with CO2 24 hours after the last treatment.

### Enzymatic dissociation of mouse dorsal skin and flow cytometry

On the day of harvest, dorsal skin was shaved using hair clippers and the treated skin patch was harvested using scissors. Subcutaneous adipose tissue was removed, and skin tissue was transferred to cold PBS. Mouse skin was finely chopped using sterile razor blades, and the tissue was transferred into a gentleMACS C-tube containing 1.74ml Buffer L, 50µl Enzyme P, 200µl Enzyme D and 10µl Enzyme A. Enzymatic dissociation was performed for 2.5h at 37°C and after incubation 2ml of DMEM containing 2% FBS and 10 mM HEPES at room temperature were added. Tissue was mechanically dissociated using a gentleMACS dissociator run on program h_skin_01 and for enzyme quenching 12µl 0.5M EDTA and 400µl FBS were added. The resulting cellular suspension was filtered through a 70µm cell strainer (Falcon, #352350) and cells were spun down at 400g for 10min at 4°C. Supernatant was aspirated and cells were resuspended in 1ml ACK buffer (A10492-01). Suspensions were transferred to 1.5ml tubes and incubated for 2min for red blood cell lysis and spun at 400g for 4min at 4°C. Supernatants were aspirated and cells were washed once in ice-cold fresh FACS buffer (PBS + 2% FCS + 2mM EDTA). Cells were resuspended in 500µl cold FACS buffer containing 1µl 7-AAD (Thermo Scientific) and filtered through a 40µm cap filter into FACS tubes. Viable cells (100.000 events) were sorted into 0.4% BSA in PBS on a Sony SH800 flow sorter. Cells were centrifuged at 400g for 10min at 4°C and supernatant was aspirated. Cells were resuspended in 0.4% BSA in PBS, counted and processed for scRNA-seq.

### ScRNA-Seq and combined sc(5’-RNA and TCR)-seq

Cells were counted and ∼8,500 cells were loaded per channel of the 10x Genomics Chromium chips using the 3’-v3 chemistry for scRNA-seq or the 5’-v1.1 chemistry for single-cell TCR-enriched V(D)J profiling. Single cells were processed through the 10x Genomics Chromium Platform according to manufacturer’s instructions (10x Genomics). Briefly, cells were encapsulated into Gel Beads in Emulsion (GEMs) droplets using the Chromium instrument. After encapsulation and cell lysis, cDNA expression libraries were generated following the manufacturer’s protocol, including barcoded reverse transcription, cDNA amplification, enzymatic fragmentation, adapter ligation and sample indexing steps. Libraries were quantified using Qubit dsDNA High Sensitivity assay kit (ThermoFisher Scientific). Fragment sizes of final libraries were assessed and quantified using DNA High Sensitivity Bioanalyzer Chip (Agilent), multiplexed and sequenced on Illumina Nextseq500, using a high output 150 cycle kit and the following read structure: Read 1: 28 cycles, Read 2: 96 cycles, Index Read 1: 8 cycles.

### ScATAC-Seq

Fixed and washed cell suspensions were lysed using 10x Genomics scATAC lysis buffer (10mM Tris-HCl pH 7.4, 10mM NaCl, 3mM MgCl2, 0.1% NP-40, 0.1% Tween-20, 0.01% Digitonin, 1% BSA) for 3min on ice. After lysis, nuclei were washed by addition of 1ml cold scATAC wash buffer (10x Genomics; lysis buffer without NP-40 and Digitonin), inverted and centrifuged (5min, 500g, 4°C). Supernatant was aspirated, and pelleted nuclei were resuspended in 1x Nuclei Dilution Buffer (10x Genomics). Nucleus aliquots were mixed with Trypan Blue to assess complete lysis and nuclei were counted using a Countess II FL Automated Cell Counter. Processing of nuclei for tagmentation, loading of 10x Genomics chips and droplet encapsulation via the 10x Genomics Chromium controller microfluidics instrument was performed according to Chromium Single Cell ATAC Solution protocol. ∼8,500 nuclei were loaded per channel of the 10x Genomics Chromium chips using the v1 kit. Libraries were quantified using Qubit dsDNA High Sensitivity assay kit (ThermoFisher Scientific). Fragment sizes of final libraries were assessed and quantified using DNA High Sensitivity Bioanalyzer Chip (Agilent), multiplexed and sequenced on Illumina Nextseq500 using a high output 75 cycle kit and the following read structure: Read 1: 34 cycles, Read 2: 34 cycles, Index Read 1: 8 cycles, Index Read 2: 16 cycles.

### scRNA-seq data preprocessing

10X Cell Ranger v3 pipeline was used to align reads and generate count matrices, with the “cellranger mkfastq” and “cellranger count” commands, respectively. EmptyDrops function from the DropletUtils v1.6.1 R package was run on the unfiltered count matrices with ignore=10, retain=800 and lower=200 parameters with an FDR 1% cutoff. Barcodes with a total number of UMIs or genes < 200 were filtered out. Percent mitochondrial UMI cutoffs of 10% and 20% were used for the samples profiled with 10X v2 and v3 chemistry, respectively. Counts were normalized with a log(TP10K+1) transformation via the sc.pp.normalize_total function of Scanpy v1.8. Preprocessing and normalization steps were performed similarly for human scleroderma and mouse (control and MC903) scRNA-seq datasets. QC steps resulted in 29,689 and 9,133 high quality cell profiles in scleroderma and mouse datasets, respectively.

### Annotation of coarse and granular cell subsets

PCA was performed with the top 2,000 highly variable genes and a *k*-nearest neighbor (*k-*NN) graph was built with *k*=15 on 50 PCs using sc.pp.pca and sc.pp.neighbors functions of Scanpy, respectively. The *k*-NN graph was clustered with the Leiden community detection method[78,79] to partition cells into coarse clusters using the sc.tl.leiden in Scanpy. Marker genes were found using one-*vs*-rest two-tailed Welch’s t-test using sc.tl.rank_genes_groups in Scanpy. For coarse cell type annotations, markers for each cluster were compared to literature-derived known markers. For the granular subset annotations, first, PCA was performed with the top 1,000 highly variable genes for each coarse cell type separately and a *k*-NN graph was built with *k*=15 on 50 PCs for each coarse cell type. Cells of the same coarse cell type were then clustered into more granular subsets using the Leiden algorithm with varying levels of resolution (ranging from 0.5 to 3.0) and were iteratively visualized with and without batch correction (PyTorch implementation of Harmony[80]). The harmonize() function from the PyTorch implementation of Harmony (https://github.com/lilab-bcb/harmony-pytorch[81]) was used with channel IDs as a batch covariate based on the PCA representation of cells for removing unwanted inter-sample and inter-individual variation. The cell subset markers, which were found similarly to the coarse cell type markers, were then compared with known literature markers in both batch-corrected and uncorrected views of the data. UMAP (Uniform Manifold Approximation and Projection)[82] was used for visualizations. Normalization, PCA, *k*-NN, data harmonization, clustering and data-driven marker identification steps were performed similarly to those of human healthy and AD samples. Only coarse cell types were annotated in mouse and scleroderma scRNA-seq datasets.

### Differential expression

Negative binomial regression on raw counts using the diffxpy (https://github.com/theislab/diffxpy/releases, v0.1.7) Python package. ‘∼ 1 + sex + mt_frac + log10_n_umis + disease_status’ formula was used to compare healthy *vs*. lesional and healthy *vs*. non-lesional groups within each coarse and granular subset, while correcting for the fraction of mitochondrial UMIs, total number of UMIs and sex of the donor. Wald test was performed to estimate the effects of disease status.

### Proportion analysis of single-cell and bulk RNA-seq data

The dirichreg function from the DirichletReg R package (v0.7) was used for testing the significance of differences in cell subset proportions across conditions for both scRNA-seq and deconvolved bulk RNA-seq. ‘’y ∼ chemistry_10x + disease_status | disease_status’ formula was used to fit the proportion models with the alternative parametrization for each cell subset separately in a one-*vs*-rest manner. The effect of the diseae_status variable was tested for significance using a likelihood ratio test. The ggplot R package with the geom_boxplot2 function from the Ipaper R package was used for Box plot visualizations. geom_boxplot2 function was run with the “width.errorbar=0” argument. Lesional samples of patients BCH04 and BCH07 were excluded from the proportion analysis due to their lack of robust levels of epidermal inflammatory marker gene expression (**Figure S3f,g**).

### Deconvolution of bulk RNA-seq

Bulk RNA-seq data[16] were downloaded from Gene Expression Omnibus (GEO, accession GSE121212). Psoriasis samples were discarded. The ag.optimize function from the autogenes Python package[83] was run on log(TPM+1)-transformed scRNA-seq expression values with the arguments “ngen=5000, seed=0, nfeatures=400, mode=’fixed’, offspring_size=100”. Granular subset annotations were used for the marker optimization. Next, the ad.deconvole function was run with the NuSVR method on log(TPM+1) transformed bulk RNA-seq expression values. Negative weights were set to zero.

### Preprocessing of published skin scRNA-seq data

Raw counts of published scRNA-seq were downloaded either from GEO (GSE158432[12], GSE153760[11], GSE147424[8]) or the HCA skin portal[10] at https://developmentcellatlas.ncl.ac.uk/datasets/hca_skin_portal/.

### Cell type classification

A multinomial logistic ridge regression classifier was used to classify cell profiles from the published scRNA-seq skin datasets, and the 5’ scRNA-seq and scleroderma scRNA-seq datasets in this study. log(TP10k+1)-transformed expression values were concatenated across the published scRNA-seq skin datasets and the present study for a total of 921,832 cells. To maintain comparable regression coefficients across genes, z-score-transformed expression values across the five datasets were used for the fit. Protein coding genes were filtered for expression in at least 30 cells for the classification, which resulted in 17,494 genes.

LogisticRegressionCV class from scikit-learn Python package was instantiated with parameters “class_weight=’balanced’, scoring=’balanced_accuracy’, n_jobs=12, Cs=numpy.logspace(-6, -4, 20))” to determine the *L*2 regularization coefficient via stratified 5-fold cross-validation with a balanced accuracy metric.

To classify the (5’RNA + TCR)-seq dataset, mean expression was subtracted and multiplied with standard deviation values calculated across the cells in five datasets to bring the expression levels to the same level as the rest.

### Graph abstraction analysis

For the undirected and directed PAGA graphs, sc.tl.paga and scv.tl.paga functions were used from Scanpy and scVelo packages, respectively. In the directed PAGA analysis of DC subsets (**Figure 3c)**, RNA velocity values (below) were used with velocity pseudotime as a prior to determine the directionality.

### RNA velocity

Velocyto.py command line interface (v0.17.15)[21] was used to count spliced and unspliced reads using the “velocyto run possorted_genome_bam.bam cellranger-GRCh38-3.0.0-genes.gtf -o sample -m hg38_rmsk.gtf -b barcodes_sample.csv -@ 30 -vv --samtools-memory 100000 -e sample” command. Repetitive elements were masked using the RepeatMasker track downloaded from UCSC Genome Browser in GTF format. Downstream analysis was performed using the scv.pp.moments, scv.tl.recover_dynamics, scv.tl.velocity, scv.tl.paga, and scv.tl.velocity_pseudotime functions of scVelo (v0.2.1)[22]. scv.tl.velocity was run in stochastic and dynamical modes for DCs and KCs, respectively. Patients BCH01, BCH04, BCH07 were excluded in the RNA velocity analysis of keratinocytes due to high inter-individual variation introduced by these samples.

### Pseudotime trajectories of keratinocytes and dendritic cells

For the keratinocyte pseudotime trajectories, KC 5, cKC1 and cKC2 subsets were subsetted from all keratinocytes. PCA was fitted using top 500 highly variable genes. Inter-individual effects were removed using the harmonize() function in the PyTorch version of Harmony[81] based on PCA representation of cells (50 PCs) with individual IDs as batch covariate. *k*-NN graph was built using batch corrected Harmony representations with 150 neighbors (*k*=150). sc.tl.diffmap and sc.tl.dpt in Scanpy[23,84] to calculate the trajectories. cKC populations were collapsed to gain more statistical power.

For the dendritic cell pseudotime trajectories, we used the latent time inference based on the RNA velocities of six DC subsets (mmDC, *MMP12*^+^ DC, Langerhans cells, DC2s, *CD83*^+^ DCs, *IL1B*^+^ DCs), as inferred by scVelo (described above). scv.tl.latent_time function was used to unify the gene-specific latent timepoints into a universal latent time.

Pseudotime distributions were plotted using the Python implementation of grammar of graphics, plotnine (https://github.com/has2k1/plotnine, version 0.8). Comparison of pseudotime distributions between conditions were performed using the scipy.stats.mannwhitneyu function.

### Gene set enrichment analysis

The sc.queries.enrich function from Scanpy was used for gene set enrichment analysis with REACTOME gene sets and the arguments “gprofiler_kwargs={’no_evidences’: False, ‘sources’:[’REAC’], ‘all_results’: True}”. FDR < 0.1 and log2(fold change) > 0.5 cutoffs were used for fibroblast and KC marker analyses. For differentially expressed genes between healthy vs lesional KCs, FDR < 0.1 and log2 fold change > 1 cutoffs were used.

### scATAC-seq data preprocessing

Raw sequencing data were demultiplexed using CellRanger-ATAC mkfastq and aligned to a mtDNA blacklist-modified[85] hg19 human reference genome using CellRanger-ATAC count v1.1. Fragment files from this pre-processing were used as input to ArchR[86]. Cell barcodes displayed high median transcription start site enrichment score (9.9), high number of median ATAC fragments per cell (25,177) and expected ATAC fragment size distribution. A total of 3,220 high-quality cells with a minimum TSS score of 5 and at least 2,500 nuclear chromatin accessibility fragments were identified. Dimensionality reduction was performed using latent semantic indexing for the top 25,000 variable genomic loci bins, and the top 30 components were retained. Cell clustering was performed using shared nearest neighbor modularity optimization on a *k*=10 *k*-NN graph. Chromatin accessibility peaks were called for each cluster and then aggregated based on the peaksummit accessibility--all default settings in the ArchR workflow. Gene activity scores for established marker genes from the literature facilitated the identification of keratinocytes, fibroblasts, endothelial cells, and immune cell subsets. Single-cell TF activities were computed using chromVAR[87] using the default settings, including a bias-correction of 50 background peaks per accessibility locus. The keratinocyte trajectory was inferred from the addTrajectory function with a user-specified ordering of the cluster medoids. Heatmaps of TF and gene activities were visualized on this trajectory using the plotTrajectoryHeatmap function using the default hyperparameters.

### Topic modeling of lymphocytes

A stochastic block model-based hierarchical topic modeling approach[46] (https://github.com/martingerlach/hSBM_Topicmodel) was used. The number of topics (253) was determined automatically during the SBM inference using the minimum description length-based Bayesian model selection approach. For visualization of differential topic weights between conditions, an empirical cumulative distribution function (ECDF) of topic weights was calculated by the distributions.empirical_distribution ECDF function from the statsmodels Python package (v0.11.1).

T cells with topic 190 (pathological T cell topic) weights greater than the 90^th^ percentile of the weight distribution of this topic (0.0049) were assigned the label of “T *IL13^+^/IL22^+^/IL26^+^*”.

### Correlation analysis of cytokine programs

The corrcoef function in the NumPy Python package was used to find genes correlated with *IL5, IL13, IL22, IL26, IL31* within lymphocytes. Barplot visualizations were produced using the plotnine (v0.7) Python package.

### TF activity analysis

The Python implementation of Dorothea (https://github.com/saezlab/dorothea-py, commit=5e3ee0e)[88,89] was used to calculate the activity scores of 118 transcription factors, which are in A and B confidence categories of Dorothea, for each cell. dorothea.run function was used with “dorothea.load_regulons([’A’, ‘B’]), center=True, scale=True, use_raw=False” arguments.

For the healthy vs. lesional and non-lesional differential activity analysis, a linear regression model for each cell subset was fit with the formula “score ∼ 1 + sex + mt_frac + log10_n_umis + disease_status” where disease_status is a binary variable indicating whether a cell has healthy or lesional/non-lesional status. statsmodels.formula.api.ols function from the statsmodels Python package (v0.11.1) was used for the regression analysis.

### Preprocessing and analysis of joint single cell (5’RNA + TCR)-seq

Fastq, h5 and JSON files representing the reads, 5’ gene expression and VDJ information were o obtained with Cell Ranger (v2.0.2) V(D)J pipeline commands “cellranger mkfastq” and “cellranger vdj”.

Scirpy[90] (v0.5.0) was used for the TCR analysis. First, for each patient, TCR data were loaded in json format using the ir.io.read_10x_vdj function with “patient_tcr.json’, filtered=True” arguments. After merging 5’ scRNA-seq expression data with TCR information, PCA was fitted with top 2,000 highly variable genes on the log(TP10K+1)-normalized expression matrix. Graph-based Leiden clustering was run with resolution=1.0 parameter on the *k*-NN graph built with *k*=15 on 50 PCs to annotate the T cell cluster which showed high CD3D and CD3E expression. Only cells in this cluster that also have TCR information were retained for downstream analysis, yielding 1,138 cells from two patients. Granular cell subset annotations were predicted using the cell subset classifier trained on the 3’ scRNA-seq data.

Data from each patient were merged and integrated using the harmonize() function of the PyTorch implementation of Harmony[81] similarly to the procedure described in the “Annotation of coarse and granular cell subsets” subsection. Clonotypes were determined jointly using the amino acid sequences of the CDR3 region on the VDJ receptor arms (TCR-β, TCR-δ, and IG-heavy chains) by the ir.pp.ir_neighbors(adata, sequence=’aa’, receptor_arms=’VDJ’) and ir.tl.define_clonotype_clusters(adata) functions of Scirpy, where a clone is defined as all cells with an identical VDJ CDR3 amino acid sequence. Clones of two or more were labeled as expanded.

Two-tailed Welch’s t-test via sc.tl.rank_genes_groups function from the Scanpy Python package was used to identify genes differentially expressed between cells from expanded (n>=2) *vs*. non-expanded (n=1) clones. sc.tl.score_genes function in Scanpy was used with genes that have non-zero probabilities in Topic 190 (**Supplementary Table 9**) to calculate the average expression of Topic 190 in the T cells with VDJ information. scipy.stats.zscore function was used to z-score the calculated Topic 190 scores.

### Sample PCA by granular cell subset proportions

Granular cell type proportions (including the *IL13*^+^/*IL22*^+^/*IL26*^+^ T cell subset) of each channel were ‘centered log-ratio’ (CLR)-transformed using the ‘clr’ function from the ‘composition_stats’ Python package. For PCA embeddings of channels, the sc.pp.pca() function from Scanpy package was used with the CLR-transformed proportions. BCH01 non-lesional channels (BCH01-NL1 and BCH01-NL2) were not used in the analysis since these samples have absolute z-score proportion values beyond 2.0 in 29 (BCH01-NL1) and 32 (BCH01-NL2) out of 87 cell subsets. Proportions shown in the heatmaps and PCA scatterplots (**Figure 6a** and **Figure S14a,d,g**) are z-score-transformed.

### Granular cell type embeddings using proportions

Granular cell subset proportions (including *IL13*^+^/*IL22*^+^/*IL26*^+^ T cells) were calculated for each channel. After CLR and z-score transformations, cell subsets in the transposed matrix (cell subset × channel) were embedded by UMAP with a *k*-NN graph representation (*k*=20) of cell subsets. sc.pp.neighbors and sc.tl.umap functions were used for the embedding. clr() function from the composition-stats Python package (v2.0.0) was used for the centered log-ratio transform. BCH01 non-lesional channels (BCH01-NL1 and BCH01-NL2) were excluded from the analysis as explained above.

### Correlation of cell subsets

Spearman correlation coefficients of the granular cell subset proportion profiles for each pair of samples were calculated using the DataFrame.corr() function of Pandas Python package with the method=’spearman’ argument. Correlations were visualized as a heat map using the ‘clustermap’ function from the Seaborn Python package.

### Receptor-ligand analysis

Putative cell-cell interactions were identified between cell subsets that are significant in at least one lesional channel using the command line interface of CellPhoneDB v2.1[91] (“cellphonedb method statistical_analysis meta.tsv counts.tsv --counts-data hgnc_symbol --project-name ad -- threads 90 --subsampling --subsampling-num-cells 3000 --subsampling-log false”). Next, z-score-transformed expression level of the receptor and ligand was calculated. For a cell subset of interest, z-score within the coarse cell type was used, whereas for the other cell subsets potentially interacting with the cell subset of interest, global z-score was used. Finally, significant receptor or ligand genes that are highly expressed in either cell subset of interest or its partners were manually selected using the mean z-score of receptor and ligand expression. For the *IL13^+^*/*IL22^+^*/*IL26^+^* T cell subset, gene selections were made based on expression cutoffs (z-score > 1.0) instead of the CellPhoneDB significance.

For receptor cell frequency and ligand mean expression correlations[62], granular cell subset proportions across lesional sample channels that had at least six cells expressing the ligand gene (log(TP10K+1) > 0) were used. Z-score-transformed sample-level mean expression values were correlated with cell type proportions using the DataFrame.corr() function from the Pandas Python package (version 1.3.5) with “spearman” parameter.

### Non-human primate data preprocessing and integration

A *Macaca mulatta* DRG sensory neuron scRNA-seq dataset[49] was downloaded from GEO (GSE165566). The top 2,000 highly variable genes were identified using the sc.pp.highly_variable_genes function of Scanpy. PCA was fitted using the highly variable genes. Next, data across three profiled individuals was integrated using the PyTorch implementation of Harmony in the PC space with 50 PCs and a *k*-nearest neighbors (*k*-NN) graph (*k*=15) was constructed using batch-corrected Harmony representation of cells using harmony.harmonize(adata.obsm[’X_pca’], adata.obs, ‘sample_name’) and sc.pp.neighbors(adata, use_rep=’X_harmony’) functions. The cell type annotations provided by the authors were used.

### GWAS enrichment analysis

GWAS-nominated genes for 2,963 diseases and traits were downloaded from the Open Targets Genetics (OTG)[67] GraphQL API (https://github.com/opentargets/genetics-api, v20.02). Fisher’s exact test implemented in Fisher python package was used to test of gene sets nominated by the Locus2Gene (L2G) framework of OTG for a particular phenotype were enriched in markers of each of the 86 cell subsets and the *IL13*^+^/*IL22*^+^/*IL26*^+^ T cell subset, to a total of 87 subsets. Cell subset marker genes were defined in six different ways and each tested for enrichment. First, three types of markers were defined as (**1**) genes with higher than z-score of 1.0; (**2**) genes with top 200 highest z-score values; (**3**) genes with top 50 highest z-score values. In each of these three cases, z-scores were calculated either (**1**) across all cells (*i.e.*, global marker signatures); or (**2**) across cells within each coarse cell type annotation (*i.e.*, local marker signatures), overall yielding six gene sets for each of the 86 granular subsets. GWAS enrichment analyses were performed with 87 subsets.

For the AD GWAS gene set, the union of genes nominated in GWAS were used, where the trait name is one of eczema, atopic dermatitis, “inflammatory skin disease [Atopic dermatitis”, “atopic dermatitis [random effects]”, “atopic dermatitis [EA, fixed effects]”, “Eczema/dermatitis | non-cancer illness code, self-reported” in the OTG trait list. This led to 9 GWAS, including 8 published studies[6,92–98], and the UK Biobank GWAS round 2 results of Neale lab (http://www.nealelab.is/uk-biobank).

### Differential expression of lymphocyte topics between conditions

Differential expression analysis of topic expression was performed using Welch’s two-sided *t* test (sc.tl.rank_genes_groups function in Scanpy). Benjamini-Hochberg FDR values are used in the figures. ggplot function from the plotnine plotting package was used for visualizations of the DE results.

## References

1. Langan SM, Irvine AD, Weidinger S. Atopic dermatitis. Lancet. 2020;396: 345–360.

2. Jungersted JM, Scheer H, Mempel M, Baurecht H, Cifuentes L, Høgh JK, et al. Stratum corneum lipids, skin barrier function and filaggrin mutations in patients with atopic eczema. Allergy. 2010;65: 911–918.

3. Sanyal RD, Pavel AB, Glickman J, Chan TC, Zheng X, Zhang N, et al. Atopic dermatitis in African American patients is T2/T22-skewed with T1/T17 attenuation. Ann Allergy Asthma Immunol. 2019;122: 99–110.e6.

4. Nettis E, Ferrucci SM, Ortoncelli M, Pellacani G, Foti C, Di Leo E, et al. Use of Dupilumab in 543 Adult Patients With Moderate-to-Severe Atopic Dermatitis: A Multicenter, Retrospective Study. J Investig Allergol Clin Immunol. 2022;32: 124–132.

5. Renert-Yuval Y, Guttman-Yassky E. New treatments for atopic dermatitis targeting beyond IL-4/IL-13 cytokines. Ann Allergy Asthma Immunol. 2020;124: 28–35.

6. Paternoster L, Standl M, Chen C-M, Ramasamy A, Bønnelykke K, Duijts L, et al. Meta-analysis of genome-wide association studies identifies three new risk loci for atopic dermatitis. Nat Genet. 2011;44: 187–192.

7. Stubbington MJT, Rozenblatt-Rosen O, Regev A, Teichmann SA. Single-cell transcriptomics to explore the immune system in health and disease. Science. 2017;358: 58–63.

8. He H, Suryawanshi H, Morozov P, Gay-Mimbrera J, Del Duca E, Kim HJ, et al. Single-cell transcriptome analysis of human skin identifies novel fibroblast subpopulation and enrichment of immune subsets in atopic dermatitis. J Allergy Clin Immunol. 2020;145: 1615–1628.

9. Hughes TK, Wadsworth MH 2nd, Gierahn TM, Do T, Weiss D, Andrade PR, et al. Second-Strand Synthesis-Based Massively Parallel scRNA-Seq Reveals Cellular States and Molecular Features of Human Inflammatory Skin Pathologies. Immunity. 2020;53: 878–894.e7.

10. Reynolds G, Vegh P, Fletcher J, Poyner EFM, Stephenson E, Goh I, et al. Developmental cell programs are co-opted in inflammatory skin disease. Science. 2021;371. doi:10.1126/science.aba6500

11. Rojahn TB, Vorstandlechner V, Krausgruber T, Bauer WM, Alkon N, Bangert C, et al. Single-cell transcriptomics combined with interstitial fluid proteomics defines cell type-specific immune regulation in atopic dermatitis. J Allergy Clin Immunol. 2020;146: 1056–1069.

12. Bangert C, Rindler K, Krausgruber T, Alkon N, Thaler FM, Kurz H, et al. Persistence of mature dendritic cells, TH2A, and Tc2 cells characterize clinically resolved atopic dermatitis under IL-4Rα blockade. Science immunology. 2021;6. doi:10.1126/sciimmunol.abe2749

13. Cheng JB, Sedgewick AJ, Finnegan AI, Harirchian P, Lee J, Kwon S, et al. Transcriptional Programming of Normal and Inflamed Human Epidermis at Single-Cell Resolution. Cell Rep. 2018;25: 871–883.

14. Wang S, Drummond ML, Guerrero-Juarez CF, Tarapore E, MacLean AL, Stabell AR, et al. Single cell transcriptomics of human epidermis identifies basal stem cell transition states. Nat Commun. 2020;11: 4239.

15. Gur C, Wang S-Y, Sheban F, Zada M, Li B, Kharouf F, et al. LGR5 expressing skin fibroblasts define a major cellular hub perturbed in scleroderma. Cell. 2022;185: 1373–1388.e20.

16. Tsoi LC, Rodriguez E, Degenhardt F, Baurecht H, Wehkamp U, Volks N, et al. Atopic Dermatitis Is an IL-13-Dominant Disease with Greater Molecular Heterogeneity Compared to Psoriasis. J Invest Dermatol. 2019;139: 1480–1489.

17. Candi E, Schmidt R, Melino G. The cornified envelope: a model of cell death in the skin. Nat Rev Mol Cell Biol. 2005;6: 328–340.

18. Fuchs E, Raghavan S. Getting under the skin of epidermal morphogenesis. Nat Rev Genet. 2002;3: 199–209.

19. Wolf FA, Hamey FK, Plass M, Solana J, Dahlin JS, Göttgens B, et al. PAGA: graph abstraction reconciles clustering with trajectory inference through a topology preserving map of single cells. Genome Biol. 2019;20: 59.

20. Akinduro O, Sully K, Patel A, Robinson DJ, Chikh A, McPhail G, et al. Constitutive Autophagy and Nucleophagy during Epidermal Differentiation. J Invest Dermatol. 2016;136: 1460–1470.

21. La Manno G, Soldatov R, Zeisel A, Braun E, Hochgerner H, Petukhov V, et al. RNA velocity of single cells. Nature. 2018;560: 494–498.

22. Bergen V, Lange M, Peidli S, Wolf FA, Theis FJ. Generalizing RNA velocity to transient cell states through dynamical modeling. Nat Biotechnol. 2020;38: 1408–1414.

23. Haghverdi L, Büttner M, Wolf FA, Buettner F, Theis FJ. Diffusion pseudotime robustly reconstructs lineage branching. Nat Methods. 2016;13: 845–848.

24. de Guzman Strong C, Wertz PW, Wang C, Yang F, Meltzer PS, Andl T, et al. Lipid defect underlies selective skin barrier impairment of an epidermal-specific deletion of Gata-3. J Cell Biol. 2006;175: 661–670.

25. Ting SB, Caddy J, Hislop N, Wilanowski T, Auden A, Zhao L-L, et al. A homolog of Drosophila grainy head is essential for epidermal integrity in mice. Science. 2005;308: 411–413.

26. Mills AA, Zheng B, Wang XJ, Vogel H, Roop DR, Bradley A. p63 is a p53 homologue required for limb and epidermal morphogenesis. Nature. 1999;398: 708–713.

27. Mitamura Y, Nunomura S, Nanri Y, Ogawa M, Yoshihara T, Masuoka M, et al. The IL-13/periostin/IL-24 pathway causes epidermal barrier dysfunction in allergic skin inflammation. Allergy. 2018;73: 1881–1891.

28. Lachner J, Mlitz V, Tschachler E, Eckhart L. Epidermal cornification is preceded by the expression of a keratinocyte-specific set of pyroptosis-related genes. Sci Rep. 2017;7: 17446.

29. Li M, Hener P, Zhang Z, Kato S, Metzger D, Chambon P. Topical vitamin D3 and low-calcemic analogs induce thymic stromal lymphopoietin in mouse keratinocytes and trigger an atopic dermatitis. Proc Natl Acad Sci U S A. 2006;103: 11736–11741.

30. Joost S, Zeisel A, Jacob T, Sun X, La Manno G, Lönnerberg P, et al. Single-Cell Transcriptomics Reveals that Differentiation and Spatial Signatures Shape Epidermal and Hair Follicle Heterogeneity. Cell Syst. 2016;3: 221–237.e9.

31. Kim BS, Wang K, Siracusa MC, Saenz SA, Brestoff JR, Monticelli LA, et al. Basophils promote innate lymphoid cell responses in inflamed skin. J Immunol. 2014;193: 3717–3725.

32. Leyva-Castillo JM, Sun L, Wu S-Y, Rockowitz S, Sliz P, Geha RS. Single-cell transcriptome profile of mouse skin undergoing antigen-driven allergic inflammation recapitulates findings in atopic dermatitis skin lesions. J Allergy Clin Immunol. 2022;150: 373–384.

33. Jaitin DA, Adlung L, Thaiss CA, Weiner A, Li B, Descamps H, et al. Lipid-Associated Macrophages Control Metabolic Homeostasis in a Trem2-Dependent Manner. Cell. 2019;178: 686–698.e14.

34. Villani A-C, Satija R, Reynolds G, Sarkizova S, Shekhar K, Fletcher J, et al. Single-cell RNA-seq reveals new types of human blood dendritic cells, monocytes, and progenitors. Science. 2017;356. doi:10.1126/science.aah4573

35. Nakamizo S, Dutertre C-A, Khalilnezhad A, Zhang XM, Lim S, Lum J, et al. Single-cell analysis of human skin identifies CD14+ type 3 dendritic cells co-producing IL1B and IL23A in psoriasis. J Exp Med. 2021;218. doi:10.1084/jem.20202345

36. Maier B, Leader AM, Chen ST, Tung N, Chang C, LeBerichel J, et al. A conserved dendritic-cell regulatory program limits antitumour immunity. Nature. 2020;580: 257–262.

37. Hamilton JD, Suárez-Fariñas M, Dhingra N, Cardinale I, Li X, Kostic A, et al. Dupilumab improves the molecular signature in skin of patients with moderate-to-severe atopic dermatitis. J Allergy Clin Immunol. 2014;134: 1293–1300.

38. Zhang Q, He Y, Luo N, Patel SJ, Han Y, Gao R, et al. Landscape and Dynamics of Single Immune Cells in Hepatocellular Carcinoma. Cell. 2019;179: 829–845.e20.

39. Zilionis R, Engblom C, Pfirschke C, Savova V, Zemmour D, Saatcioglu HD, et al. Single-Cell Transcriptomics of Human and Mouse Lung Cancers Reveals Conserved Myeloid Populations across Individuals and Species. Immunity. 2019;50: 1317–1334.e10.

40. Gao Y, Nish SA, Jiang R, Hou L, Licona-Limón P, Weinstein JS, et al. Control of T helper 2 responses by transcription factor IRF4-dependent dendritic cells. Immunity. 2013;39: 722–732.

41. Tubo NJ, McLachlan JB, Campbell JJ. Chemokine receptor requirements for epidermal T-cell trafficking. Am J Pathol. 2011;178: 2496–2503.

42. Kumamoto Y, Linehan M, Weinstein JS, Laidlaw BJ, Craft JE, Iwasaki A. CD301b^+^ dermal dendritic cells drive T helper 2 cell-mediated immunity. Immunity. 2013;39: 733–743.

43. Dixit A, Parnas O, Li B, Chen J, Fulco CP, Jerby-Arnon L, et al. Perturb-Seq: Dissecting Molecular Circuits with Scalable Single-Cell RNA Profiling of Pooled Genetic Screens. Cell. 2016;167: 1853–1866.e17.

44. Martin JC, Chang C, Boschetti G, Ungaro R, Giri M, Grout JA, et al. Single-Cell Analysis of Crohn’s Disease Lesions Identifies a Pathogenic Cellular Module Associated with Resistance to Anti-TNF Therapy. Cell. 2019;178: 1493–1508.e20.

45. Bielecki P, Riesenfeld SJ, Hütter J-C, Torlai Triglia E, Kowalczyk MS, Ricardo-Gonzalez RR, et al. Skin-resident innate lymphoid cells converge on a pathogenic effector state. Nature. 2021;592: 128–132.

46. Gerlach M, Peixoto TP, Altmann EG. A network approach to topic models. Sci Adv. 2018;4: eaaq1360.

47. Bapat SP, Whitty C, Mowery CT, Liang Y, Yoo A, Jiang Z, et al. Obesity alters pathology and treatment response in inflammatory disease. Nature. 2022;604: 337–342.

48. Henriksson J, Chen X, Gomes T, Ullah U, Meyer KB, Miragaia R, et al. Genome-wide CRISPR Screens in T Helper Cells Reveal Pervasive Crosstalk between Activation and Differentiation. Cell. 2019;176. doi:10.1016/j.cell.2018.11.044

49. Kupari J, Usoskin D, Parisien M, Lou D, Hu Y, Fatt M, et al. Single cell transcriptomics of primate sensory neurons identifies cell types associated with chronic pain. Nat Commun. 2021;12: 1510.

50. Halim TYF, Rana BMJ, Walker JA, Kerscher B, Knolle MD, Jolin HE, et al. Tissue-Restricted Adaptive Type 2 Immunity Is Orchestrated by Expression of the Costimulatory Molecule OX40L on Group 2 Innate Lymphoid Cells. Immunity. 2018;48: 1195–1207.e6.

51. Yomogida K, Bigley TM, Trsan T, Gilfillan S, Cella M, Yokoyama WM, et al. Hobit confers tissue-dependent programs to type 1 innate lymphoid cells. Proc Natl Acad Sci U S A. 2021;118. doi:10.1073/pnas.2117965118

52. Mackay LK, Minnich M, Kragten NAM, Liao Y, Nota B, Seillet C, et al. Hobit and Blimp1 instruct a universal transcriptional program of tissue residency in lymphocytes. Science. 2016;352: 459–463.

53. Buechler MB, Pradhan RN, Krishnamurty AT, Cox C, Calviello AK, Wang AW, et al. Cross-tissue organization of the fibroblast lineage. Nature. 2021;593: 575–579.

54. Kinchen J, Chen HH, Parikh K, Antanaviciute A, Jagielowicz M, Fawkner-Corbett D, et al. Structural Remodeling of the Human Colonic Mesenchyme in Inflammatory Bowel Disease. Cell. 2018;175: 372–386.e17.

55. Madissoon E, Oliver AJ, Kleshchevnikov V, Wilbrey-Clark A, Polanski K, Richoz N, et al. A spatially resolved atlas of the human lung characterizes a gland-associated immune niche. Nat Genet. 2022;55: 66–77.

56. Nayar S, Campos J, Smith CG, Iannizzotto V, Gardner DH, Mourcin F, et al. Immunofibroblasts are pivotal drivers of tertiary lymphoid structure formation and local pathology. Proc Natl Acad Sci U S A. 2019;116: 13490–13497.

57. Dhingra N, Suárez-Fariñas M, Fuentes-Duculan J, Gittler JK, Shemer A, Raz A, et al. Attenuated neutrophil axis in atopic dermatitis compared to psoriasis reflects TH17 pathway differences between these diseases. J Allergy Clin Immunol. 2013;132: 498–501.e3.

58. Waldmann TA. The biology of interleukin-2 and interleukin-15: implications for cancer therapy and vaccine design. Nat Rev Immunol. 2006;6: 595–601.

59. Anderson DM, Maraskovsky E, Billingsley WL, Dougall WC, Tometsko ME, Roux ER, et al. A homologue of the TNF receptor and its ligand enhance T-cell growth and dendritic-cell function. Nature. 1997;390: 175–179.

60. Ingelfinger F, De Feo D, Becher B. GM-CSF: Master regulator of the T cell-phagocyte interface during inflammation. Semin Immunol. 2021;54: 101518.

61. Waskow C, Liu K, Darrasse-Jèze G, Guermonprez P, Ginhoux F, Merad M, et al. The receptor tyrosine kinase Flt3 is required for dendritic cell development in peripheral lymphoid tissues. Nat Immunol. 2008;9: 676–683.

62. Smillie CS, Biton M, Ordovas-Montanes J, Sullivan KM, Burgin G, Graham DB, et al. Intra- and Inter-cellular Rewiring of the Human Colon during Ulcerative Colitis. Cell. 2019. pp. 714–730.e22. doi:10.1016/j.cell.2019.06.029

63. Howell MD, Kim BE, Gao P, Grant AV, Boguniewicz M, DeBenedetto A, et al. Cytokine modulation of atopic dermatitis filaggrin skin expression. J Allergy Clin Immunol. 2009;124: R7–R12.

64. Kim T-G, Jee H, Fuentes-Duculan J, Wu WH, Byamba D, Kim D-S, et al. Dermal clusters of mature dendritic cells and T cells are associated with the CCL20/CCR6 chemokine system in chronic psoriasis. J Invest Dermatol. 2014;134: 1462–1465.

65. Marrakchi S, Guigue P, Renshaw BR, Puel A, Pei X-Y, Fraitag S, et al. Interleukin-36-receptor antagonist deficiency and generalized pustular psoriasis. N Engl J Med. 2011;365: 620–628.

66. Guttman-Yassky E, Krueger JG. Atopic dermatitis and psoriasis: two different immune diseases or one spectrum? Curr Opin Immunol. 2017;48: 68–73.

67. Mountjoy E, Schmidt EM, Carmona M, Schwartzentruber J, Peat G, Miranda A, et al. An open approach to systematically prioritize causal variants and genes at all published human GWAS trait-associated loci. Nat Genet. 2021;53: 1527–1533.

68. Kashiwada M, Cassel SL, Colgan JD, Rothman PB. NFIL3/E4BP4 controls type 2 T helper cell cytokine expression. EMBO J. 2011;30: 2071–2082.

69. Powell MD, Read KA, Sreekumar BK, Oestreich KJ. Ikaros Zinc Finger Transcription Factors: Regulators of Cytokine Signaling Pathways and CD4 T Helper Cell Differentiation. Front Immunol. 2019;10: 1299.

70. Ferreira MA, Vonk JM, Baurecht H, Marenholz I, Tian C, Hoffman JD, et al. Shared genetic origin of asthma, hay fever and eczema elucidates allergic disease biology. Nat Genet. 2017;49: 1752–1757.

71. Morgan DM, Ruiter B, Smith NP, Tu AA, Monian B, Stone BE, et al. Clonally expanded, GPR15-expressing pathogenic effector TH2 cells are associated with eosinophilic esophagitis. Science immunology. 2021;6. doi:10.1126/sciimmunol.abi5586

72. Vieira Braga FA, Kar G, Berg M, Carpaij OA, Polanski K, Simon LM, et al. A cellular census of human lungs identifies novel cell states in health and in asthma. Nat Med. 2019;25: 1153–1163.

73. Natsuaki Y, Egawa G, Nakamizo S, Ono S, Hanakawa S, Okada T, et al. Perivascular leukocyte clusters are essential for efficient activation of effector T cells in the skin. Nat Immunol. 2014;15: 1064–1069.

74. Uhlén M, Fagerberg L, Hallström BM, Lindskog C, Oksvold P, Mardinoglu A, et al. Proteomics. Tissue-based map of the human proteome. Science. 2015;347: 1260419.

75. Jassal B, Matthews L, Viteri G, Gong C, Lorente P, Fabregat A, et al. The reactome pathway knowledgebase. Nucleic Acids Res. 2020;48: D498–D503.

76. Solé-Boldo L, Raddatz G, Schütz S, Mallm J-P, Rippe K, Lonsdorf AS, et al. Single-cell transcriptomes of the human skin reveal age-related loss of fibroblast priming. Commun Biol. 2020;3: 188.

77. Philippeos C, Telerman SB, Oulès B, Pisco AO, Shaw TJ, Elgueta R, et al. Spatial and Single-Cell Transcriptional Profiling Identifies Functionally Distinct Human Dermal Fibroblast Subpopulations. J Invest Dermatol. 2018;138: 811–825.

78. Traag VA, Waltman L, van Eck NJ. From Louvain to Leiden: guaranteeing well-connected communities. Sci Rep. 2019;9: 5233.

79. Blondel VD, Guillaume J-L, Lambiotte R, Lefebvre E. Fast unfolding of communities in large networks. J Stat Mech. 2008;2008: P10008.

80. Korsunsky I, Millard N, Fan J, Slowikowski K, Zhang F, Wei K, et al. Fast, sensitive and accurate integration of single-cell data with Harmony. Nat Methods. 2019;16: 1289–1296.

81. Li B, Gould J, Yang Y, Sarkizova S, Tabaka M, Ashenberg O, et al. Cumulus provides cloud-based data analysis for large-scale single-cell and single-nucleus RNA-seq. Nat Methods. 2020;17: 793–798.

82. McInnes L, Healy J, Saul N, Großberger L. UMAP: Uniform Manifold Approximation and Projection. Journal of Open Source Software. 2018;3: 861.

83. Aliee H, Theis FJ. AutoGeneS: Automatic gene selection using multi-objective optimization for RNA-seq deconvolution. Cell Syst. 2021;12: 706–715.e4.

84. Wolf FA, Angerer P, Theis FJ. SCANPY: large-scale single-cell gene expression data analysis. Genome Biol. 2018;19: 15.

85. Lareau CA, Ludwig LS, Muus C, Gohil SH, Zhao T, Chiang Z, et al. Massively parallel single-cell mitochondrial DNA genotyping and chromatin profiling. Nat Biotechnol. 2021;39: 451–461.

86. Granja JM, Corces MR, Pierce SE, Bagdatli ST, Choudhry H, Chang HY, et al. ArchR is a scalable software package for integrative single-cell chromatin accessibility analysis. Nat Genet. 2021;53: 403–411.

87. Schep AN, Wu B, Buenrostro JD, Greenleaf WJ. chromVAR: inferring transcription-factor-associated accessibility from single-cell epigenomic data. Nat Methods. 2017;14: 975–978.

88. Garcia-Alonso L, Holland CH, Ibrahim MM, Turei D, Saez-Rodriguez J. Benchmark and integration of resources for the estimation of human transcription factor activities. Genome Res. 2019;29: 1363–1375.

89. Holland CH, Tanevski J, Perales-Patón J, Gleixner J, Kumar MP, Mereu E, et al. Robustness and applicability of transcription factor and pathway analysis tools on single-cell RNA-seq data. Genome Biol. 2020;21: 36.

90. Sturm G, Szabo T, Fotakis G, Haider M, Rieder D, Trajanoski Z, et al. Scirpy: a Scanpy extension for analyzing single-cell T-cell receptor-sequencing data. Bioinformatics. 2020;36: 4817–4818.

91. Efremova M, Vento-Tormo M, Teichmann SA, Vento-Tormo R. CellPhoneDB: inferring cell–cell communication from combined expression of multi-subunit ligand–receptor complexes. Nat Protoc. 2020;15: 1484–1506.

92. Hirota T, Takahashi A, Kubo M, Tsunoda T, Tomita K, Sakashita M, et al. Genome-wide association study identifies eight new susceptibility loci for atopic dermatitis in the Japanese population. Nat Genet. 2012;44: 1222–1226.

93. Weidinger S, Willis-Owen SAG, Kamatani Y, Baurecht H, Morar N, Liang L, et al. A genome-wide association study of atopic dermatitis identifies loci with overlapping effects on asthma and psoriasis. Hum Mol Genet. 2013;22: 4841–4856.

94. Baurecht H, Hotze M, Brand S, Büning C, Cormican P, Corvin A, et al. Genome-wide comparative analysis of atopic dermatitis and psoriasis gives insight into opposing genetic mechanisms. Am J Hum Genet. 2015;96: 104–120.

95. Schaarschmidt H, Ellinghaus D, Rodríguez E, Kretschmer A, Baurecht H, Lipinski S, et al. A genome-wide association study reveals 2 new susceptibility loci for atopic dermatitis. J Allergy Clin Immunol. 2015;136: 802–806.

96. Paternoster L, Standl M, Waage J, Baurecht H, Hotze M, Strachan DP, et al. Multi-ancestry genome-wide association study of 21,000 cases and 95,000 controls identifies new risk loci for atopic dermatitis. Nat Genet. 2015;47: 1449–1456.

97. Kichaev G, Bhatia G, Loh P-R, Gazal S, Burch K, Freund MK, et al. Leveraging Polygenic Functional Enrichment to Improve GWAS Power. Am J Hum Genet. 2019;104: 65–75.

98. Johansson Å, Rask-Andersen M, Karlsson T, Ek WE. Genome-wide association analysis of 350 000 Caucasians from the UK Biobank identifies novel loci for asthma, hay fever and eczema. Hum Mol Genet. 2019;28: 4022–4041.

