## Supplementary Figures for "Multi-modal skin atlas identifies a multicellular immune-stromal community associated with altered cornification and specific T cell expansion in atopic dermatitis"

### Figure S1

| | ID | Disease status | Sex | Age | Ethnicity | Race | Global disease assessment | EASI | Biopsy site lesional | Biopsy site non-lesional | Biopsy site healthy | Asthma |
|---|---|---|---|---|---|---|---|---|---|---|---|---|
| Hospital site A | BCH01 | AD | M | 27 | Non-Hispanic | Caucasian | 2 - Mild | 5.9 | Antecubital fossa (L) | Inner arm (L) | — | ? |
| | BCH04 | AD | F | 21 | Non-Hispanic | Black | 4 - Severe | 23.6 | Upper back (L) | Lower back (R) | — | ? |
| | BCH05 | AD | F | 21 | Hispanic | Black | 1 - Almost clear | 0.7 | Antecubital fossa (L) | ? | — | ? |
| | BCH06 | AD | F | 61 | Non-Hispanic | Caucasian | 4 - Severe | 36.6 | Lower back | ? | — | ? |
| | BCH07 | AD | M | 29 | Non-Hispanic | Caucasian | 2 - Mild | 3.5 | Arm (L) | ? | — | ? |
| Hospital site B | MGH103 | AD | M | ? | Hispanic | Other | 3 - Moderate | ? | Lower leg (L) | Lower leg (L) | — | No |
| | MGH104 | AD | M | ? | Non-Hispanic | Black | 3 - Moderate | ? | Back (R) | Back (R) | — | No |
| | MGH105 | AD | F | ? | Non-Hispanic | Caucasian | 3 - Moderate | ? | Thigh (R) | Thigh (R) | — | Yes |
| | MGH106 | AD | F | ? | Non-Hispanic | Caucasian | 2 - Mild | ? | Antecubital fossa (R) | Antecubital fossa (R) | — | No |
| | MGH108 | AD | M | ? | Non-Hispanic | Asian/Caucasian | 3 - Moderate | ? | Back (R) | Back (R) | — | No |
| | MGH110 | AD | F | ? | Non-Hispanic | Black | 4 - Severe | ? | Forearm (L) | Forearm (L) | — | Yes |
| | MGH101 | Sclero | F | ? | Non-Hispanic | Caucasian | 3 - Moderate | — | Chest (R) | Chest (R) | — | No |
| | MGH109 | Sclero | F | ? | Non-Hispanic | Caucasian | 4 - Severe | — | Forearm (L) | Forearm (L) | — | No |
| | Healthy1 | Healthy | F | ? | ? | ? | 0 - Clear | — | — | — | ? | No |
| | Healthy2 | Healthy | F | ? | ? | ? | 0 - Clear | — | — | — | ? | No |
| | Healthy5 | Healthy | F | ? | ? | ? | 0 - Clear | — | — | — | ? | No |
| | Healthy6 | Healthy | M | ? | ? | ? | 0 - Clear | — | — | — | ? | No |
| | Healthy7 | Healthy | M | ? | ? | ? | 0 - Clear | — | — | — | ? | No |
| | Healthy8 | Healthy | M | ? | ? | ? | 0 - Clear | — | — | — | ? | No |

### Figure S2

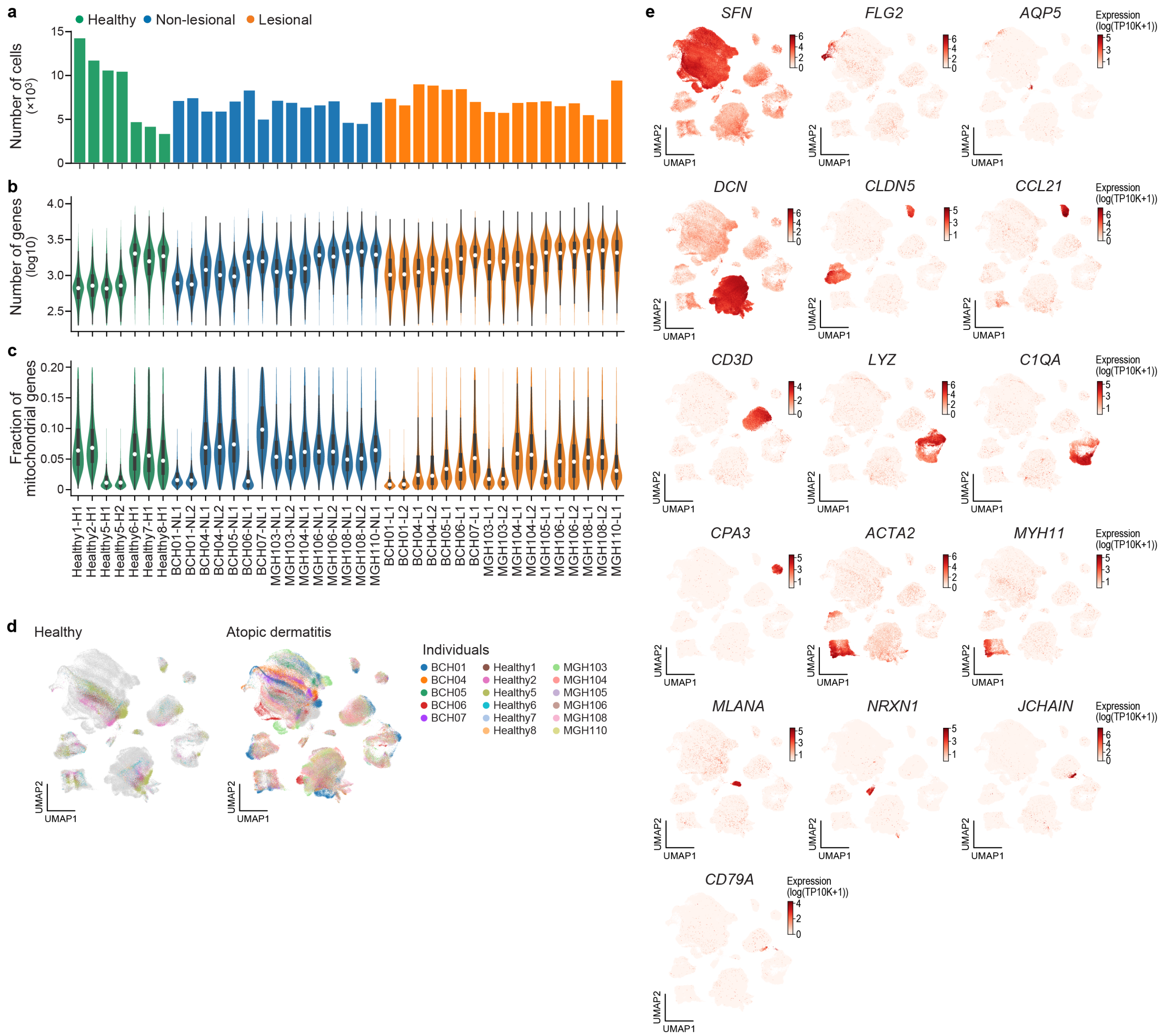
a
Healthy
Non-lesional
Lesional
Number of cells (×10³)
b
Number of genes (log10)
c
Fraction of mitochondrial genes
Healthy1-H1
Healthy2-H1
Healthy5-H1
Healthy5-H2
Healthy6-H1
Healthy7-H1
Healthy8-H1
BCH01-NL1
BCH01-NL2
BCH04-NL1
BCH04-NL2
BCH05-NL1
BCH06-NL1
BCH07-NL1
MGH103-NL1
MGH103-NL2
MGH104-NL1
MGH106-NL1
MGH106-NL2
MGH108-NL1
MGH108-NL2
MGH110-NL1
BCH01-L1
BCH01-L2
BCH04-L1
BCH04-L2
BCH05-L1
BCH06-L1
BCH07-L1
MGH103-L1
MGH103-L2
MGH104-L1
MGH104-L2
MGH105-L1
MGH106-L1
MGH106-L2
MGH108-L1
MGH108-L2
MGH110-L1
d
Healthy
Atopic dermatitis
Individuals
BCH01
BCH04
BCH05
BCH06
BCH07
Healthy1
Healthy2
Healthy5
Healthy6
Healthy7
Healthy8
MGH103
MGH104
MGH105
MGH106
MGH108
MGH110
UMAP1
UMAP2
e
SFN
FLG2
AQP5
DCN
CLDN5
CCL21
CD3D
LYZ
C1QA
CPA3
ACTA2
MYH11
MLANA
NRXN1
JCHAIN
CD79A
Expression (log(TP10K+1))

### Figure S3

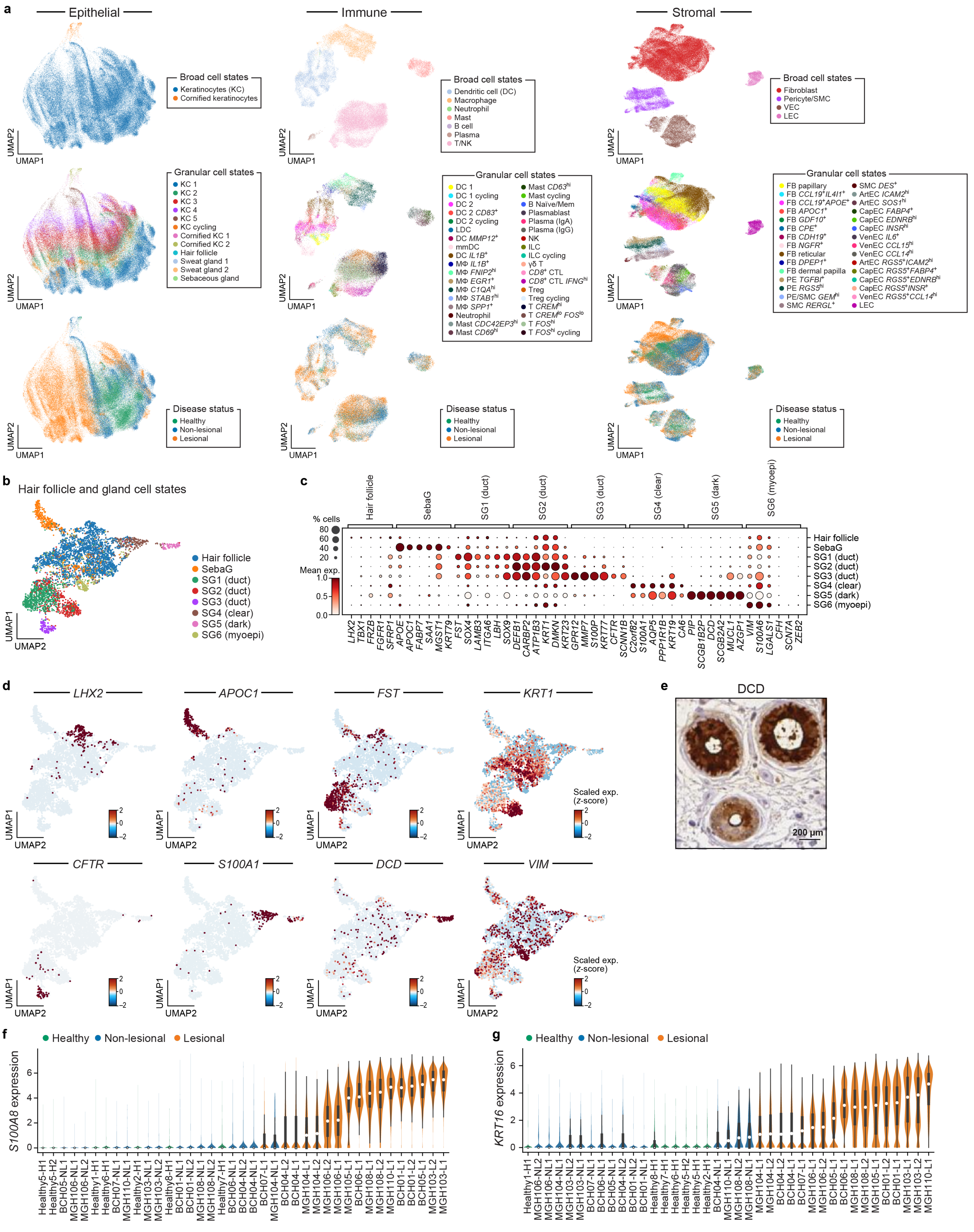

a
Epithelial
Immune
Stromal
Broad cell states
Keratinocytes (KC)
Cornified keratinocytes
Dendritic cell (DC)
Macrophage
Neutrophil
Mast
B cell
Plasma
T/NK
Fibroblast
Pericyte/SMC
VEC
LEC
Granular cell states
KC 1
KC 2
KC 3
KC 4
KC 5
KC cycling
Cornified KC 1
Cornified KC 2
Hair follicle
Sweat gland 1
Sweat gland 2
Sebaceous gland
Disease status
Healthy
Non-lesional
Lesional
UMAP1
UMAP2
b
Hair follicle and gland cell states
Hair follicle
SebaG
SG1 (duct)
SG2 (duct)
SG3 (duct)
SG4 (clear)
SG5 (dark)
SG6 (myoepi)
c
% cells
Mean exp.
d
LHX2
APOC1
FST
KRT1
CFTR
S100A1
DCD
VIM
Scaled exp. (z-score)
e
DCD
200 µm
f
S100A8 expression
g
KRT16 expression

### Figure S4

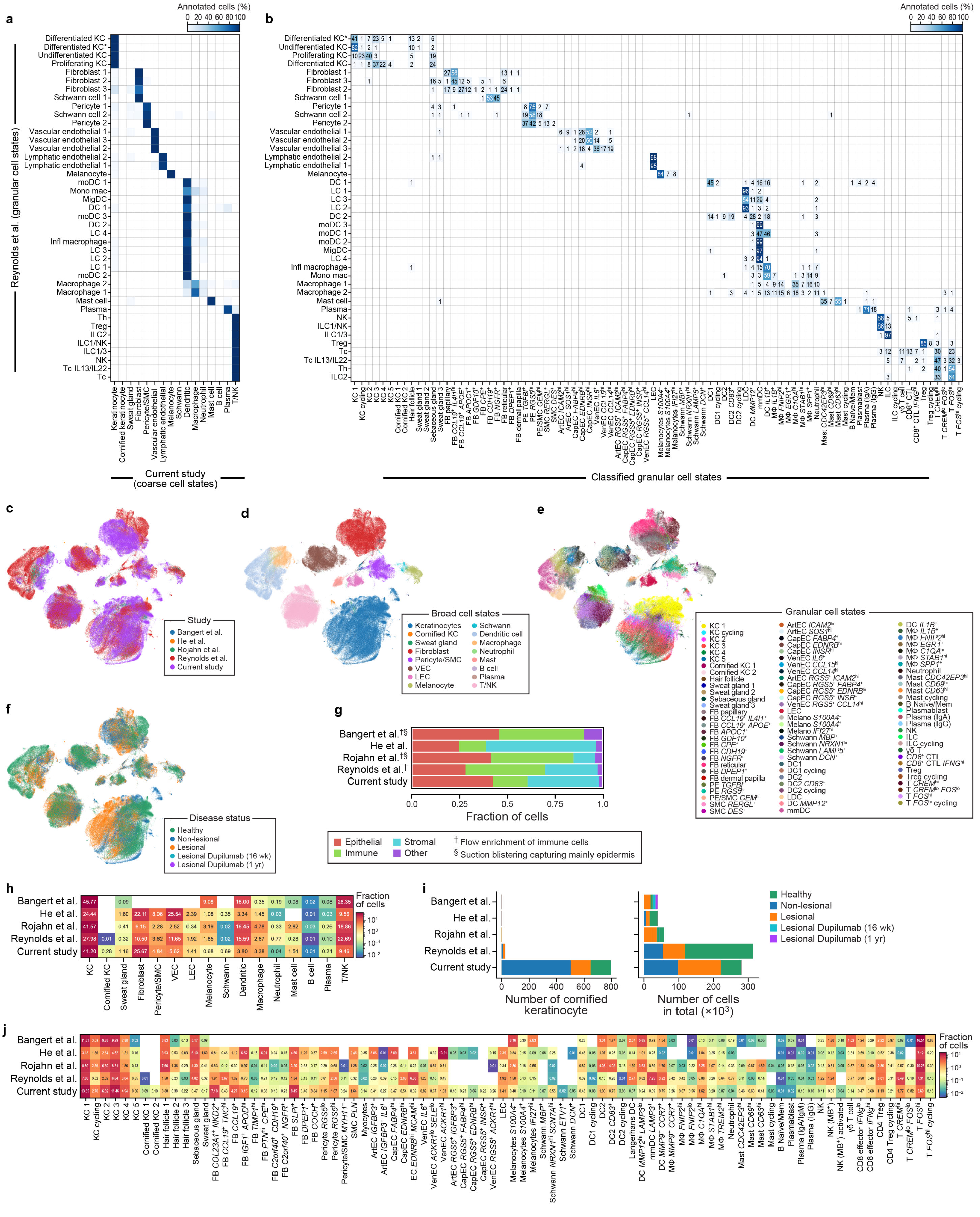

a
b
Annotated cells (%)
0 20 40 60 80 100
Reynolds et al. (granular cell states)
Current study
(coarse cell states)
Classified granular cell states
Keratinocyte
Cornified keratinocyte
Sweat gland
Fibroblast
Pericyte/SMC
Vascular endothelial
Lymphatic endothelial
Melanocyte
Schwann
Dendritic
Macrophage
Neutrophil
Mast cell
B cell
Plasma
T/NK
c
Study
Bangert et al.
He et al.
Rojahn et al.
Reynolds et al.
Current study
d
Broad cell states
Keratinocytes
Cornified KC
Sweat gland
Fibroblast
Pericyte/SMC
VEC
LEC
Melanocyte
Schwann
Dendritic cell
Macrophage
Neutrophil
Mast
B cell
Plasma
T/NK
e
Granular cell states
f
Disease status
Healthy
Non-lesional
Lesional
Lesional Dupilumab (16 wk)
Lesional Dupilumab (1 yr)
g
Bangert et al.†§
He et al.
Rojahn et al.†§
Reynolds et al.†
Current study
0.0
0.25
0.5
0.75
1.0
Fraction of cells
Epithelial
Immune
Stromal
Other
† Flow enrichment of immune cells
§ Suction blistering capturing mainly epidermis
h
Fraction of cells
KC
Cornified KC
Sweat gland
Fibroblast
Pericyte/SMC
VEC
LEC
Melanocyte
Schwann
Dendritic
Macrophage
Neutrophil
Mast cell
B cell
Plasma
T/NK
i
Number of cornified keratinocyte
Number of cells in total (×10³)
Healthy
Non-lesional
Lesional
Lesional Dupilumab (16 wk)
Lesional Dupilumab (1 yr)
j
Fraction of cells

### Figure S5

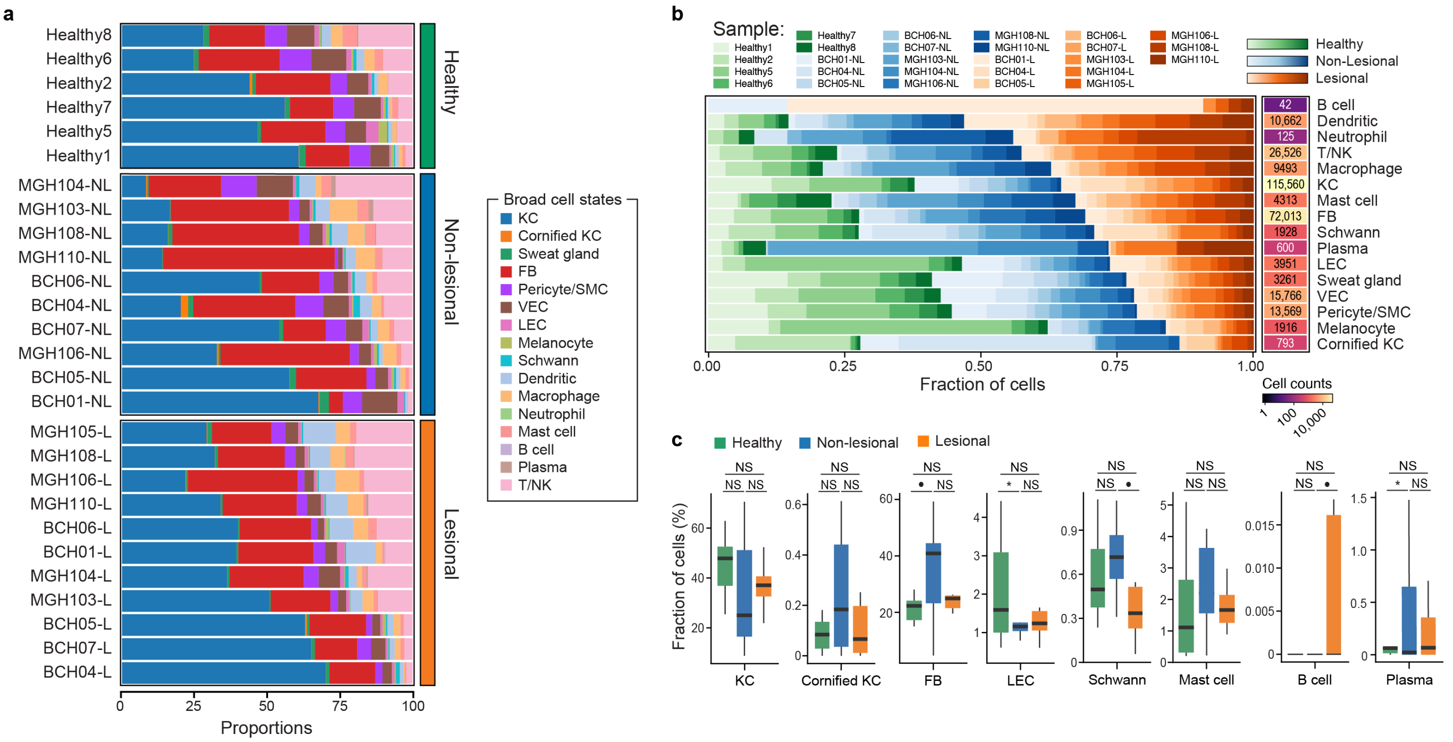
a
Healthy8
Healthy6
Healthy2
Healthy7
Healthy5
Healthy1
MGH104-NL
MGH103-NL
MGH108-NL
MGH110-NL
BCH06-NL
BCH04-NL
BCH07-NL
MGH106-NL
BCH05-NL
BCH01-NL
MGH105-L
MGH108-L
MGH106-L
MGH110-L
BCH06-L
BCH01-L
MGH104-L
MGH103-L
BCH05-L
BCH07-L
BCH04-L
Healthy
Non-lesional
Lesional
0
25
50
75
100
Proportions
Broad cell states
KC
Cornified KC
Sweat gland
FB
Pericyte/SMC
VEC
LEC
Melanocyte
Schwann
Dendritic
Macrophage
Neutrophil
Mast cell
B cell
Plasma
T/NK
b
Sample:
Healthy1
Healthy2
Healthy5
Healthy6
Healthy7
Healthy8
BCH01-NL
BCH04-NL
BCH05-NL
BCH06-NL
BCH07-NL
MGH103-NL
MGH104-NL
MGH106-NL
MGH108-NL
MGH110-NL
BCH01-L
BCH04-L
BCH05-L
BCH06-L
BCH07-L
MGH103-L
MGH104-L
MGH105-L
MGH106-L
MGH108-L
MGH110-L
Healthy
Non-Lesional
Lesional
42 B cell
10,662 Dendritic
125 Neutrophil
26,526 T/NK
9493 Macrophage
115,560 KC
4313 Mast cell
72,013 FB
1928 Schwann
600 Plasma
3951 LEC
3261 Sweat gland
15,766 VEC
13,569 Pericyte/SMC
1916 Melanocyte
793 Cornified KC
0.00
0.25
0.50
0.75
1.00
Fraction of cells
Cell counts
1
100
10,000
c
Healthy
Non-lesional
Lesional
Fraction of cells (%)
KC
Cornified KC
FB
LEC
Schwann
Mast cell
B cell
Plasma
NS
*

### Figure S6

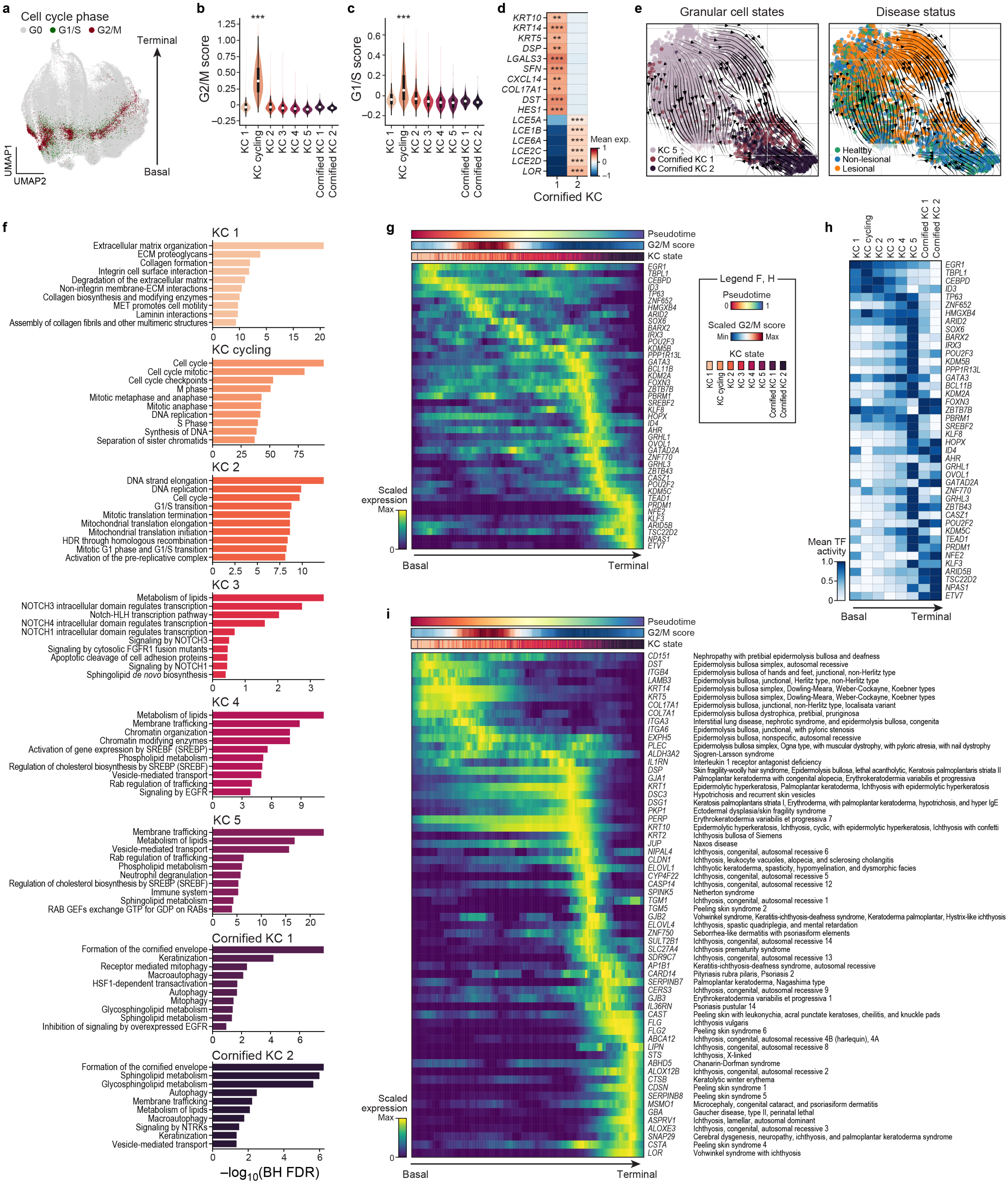

a
Cell cycle phase
G0
G1/S
G2/M
Terminal
Basal
UMAP1
UMAP2
b
G2/M score
1.25
1.00
0.75
0.50
0.25
0
−0.25
***
KC 1
KC cycling
KC 2
KC 3
KC 4
KC 5
Cornified KC 1
Cornified KC 2
c
G1/S score
0.6
0.4
0.2
0
−0.2
***
d
KRT10
KRT14
KRT5
DSP
LGALS3
SFN
CXCL14
COL17A1
DST
HES1
LCE5A
LCE1B
LCE6A
LCE2C
LCE2D
LOR
Mean exp.
1
0
−1
1
2
Cornified KC
e
Granular cell states
Disease status
KC 5
Cornified KC 1
Cornified KC 2
Healthy
Non-lesional
Lesional
f
KC 1
Extracellular matrix organization
ECM proteoglycans
Collagen formation
Integrin cell surface interaction
Degradation of the extracellular matrix
Non-integrin membrane-ECM interactions
Collagen biosynthesis and modifying enzymes
MET promotes cell motility
Laminin interactions
Assembly of collagen fibrils and other multimeric structures
KC cycling
Cell cycle
Cell cycle mitotic
Cell cycle checkpoints
M phase
Mitotic metaphase and anaphase
Mitotic anaphase
DNA replication
S Phase
Synthesis of DNA
Separation of sister chromatids
KC 2
DNA strand elongation
DNA replication
Cell cycle
G1/S transition
Mitotic translation termination
Mitochondrial translation elongation
Mitochondrial translation initiation
HDR through homologous recombination
Mitotic G1 phase and G1/S transition
Activation of the pre-replicative complex
KC 3
Metabolism of lipids
NOTCH3 intracellular domain regulates transcription
Notch-HLH transcription pathway
NOTCH4 intracellular domain regulates transcription
NOTCH1 intracellular domain regulates transcription
Signaling by NOTCH3
Signaling by cytosolic FGFR1 fusion mutants
Apoptotic cleavage of cell adhesion proteins
Signaling by NOTCH1
Sphingolipid de novo biosynthesis
KC 4
Metabolism of lipids
Membrane trafficking
Chromatin organization
Chromatin modifying enzymes
Activation of gene expression by SREBF (SREBP)
Phospholipid metabolism
Regulation of cholesterol biosynthesis by SREBP (SREBF)
Vesicle-mediated transport
Rab regulation of trafficking
Signaling by EGFR
KC 5
Membrane trafficking
Metabolism of lipids
Vesicle-mediated transport
Rab regulation of trafficking
Phospholipid metabolism
Neutrophil degranulation
Regulation of cholesterol biosynthesis by SREBP (SREBF)
Immune system
Sphingolipid metabolism
RAB GEFs exchange GTP for GDP on RABs
Cornified KC 1
Formation of the cornified envelope
Keratinization
Receptor mediated mitophagy
Macroautophagy
HSF1-dependent transactivation
Autophagy
Mitophagy
Glycosphingolipid metabolism
Sphingolipid metabolism
Inhibition of signaling by overexpressed EGFR
Cornified KC 2
Formation of the cornified envelope
Sphingolipid metabolism
Glycosphingolipid metabolism
Autophagy
Membrane trafficking
Metabolism of lipids
Macroautophagy
Signaling by NTRKs
Keratinization
Vesicle-mediated transport
−log10(BH FDR)
g
Pseudotime
G2/M score
KC state
Scaled expression
Max
0
Basal
Terminal
EGR1
TBPL1
CEBPD
ID3
TP63
ZNF652
HMGXB4
ARID2
SOX6
BARX2
IRX3
POU2F3
KDM5B
PPP1R13L
GATA3
BCL11B
KDM2A
FOXN3
ZBTB7B
PBRM1
SREBF2
KLF8
HOPX
ID4
AHR
GRHL1
OVOL1
GATAD2A
ZNF770
GRHL3
ZBTB43
CASZ1
POU2F2
KDM5C
TEAD1
PRDM1
NFE2
KLF3
ARID5B
TSC22D2
NPAS1
ETV7
Legend F, H
Pseudotime
0
1
Scaled G2/M score
Min
Max
KC state
h
Mean TF activity
1.0
0.5
0
Basal
Terminal
i
Pseudotime
G2/M score
KC state
Scaled expression
Max
0
Basal
Terminal
CD151 Nephropathy with pretibial epidermolysis bullosa and deafness
DST Epidermolysis bullosa simplex, autosomal recessive
ITGB4 Epidermolysis bullosa of hands and feet, junctional, non-Herlitz type
LAMB3 Epidermolysis bullosa, junctional, Herlitz type, non-Herlitz type
KRT14 Epidermolysis bullosa simplex, Dowling-Meara, Weber-Cockayne, Koebner types
KRT5 Epidermolysis bullosa simplex, Dowling-Meara, Weber-Cockayne, Koebner types
COL17A1 Epidermolysis bullosa, junctional, non-Herlitz type, localisata variant
COL7A1 Epidermolysis bullosa dystrophica, pretibial, pruriginosa
ITGA3 Interstitial lung disease, nephrotic syndrome, and epidermolysis bullosa, congenita
ITGA6 Epidermolysis bullosa, junctional, with pyloric stenosis
EXPH5 Epidermolysis bullosa, nonspecific, autosomal recessive
PLEC Epidermolysis bullosa simplex, Ogna type, with muscular dystrophy, with pyloric atresia, with nail dystrophy
ALDH3A2 Sjogren-Larsson syndrome
IL1RN Interleukin 1 receptor antagonist deficiency
DSP Skin fragility-woolly hair syndrome, Epidermolysis bullosa, lethal acantholytic, Keratosis palmoplantaris striata II
GJA1 Palmoplantar keratoderma with congenital alopecia, Erythrokeratodermia variabilis et progressiva
KRT1 Epidermolytic hyperkeratosis, Palmoplantar keratoderma, Ichthyosis with epidermolytic hyperkeratosis
DSC3 Hypotrichosis and recurrent skin vesicles
DSG1 Keratosis palmoplantaris striata I, Erythroderma, with palmoplantar keratoderma, hypotrichosis, and hyper IgE
PKP1 Ectodermal dysplasia/skin fragility syndrome
PERP Erythrokeratodermia variabilis et progressiva 7
KRT10 Epidermolytic hyperkeratosis, Ichthyosis, cyclic, with epidermolytic hyperkeratosis, Ichthyosis with confetti
KRT2 Ichthyosis bullosa of Siemens
JUP Naxos disease
NIPAL4 Ichthyosis, congenital, autosomal recessive 6
CLDN1 Ichthyosis, leukocyte vacuoles, alopecia, and sclerosing cholangitis
ELOVL1 Ichthyotic keratoderma, spasticity, hypomyelination, and dysmorphic facies
CYP4F22 Ichthyosis, congenital, autosomal recessive 5
CASP14 Ichthyosis, congenital, autosomal recessive 12
SPINK5 Netherton syndrome
TGM1 Ichthyosis, congenital, autosomal recessive 1
TGM5 Peeling skin syndrome 2
GJB2 Vohwinkel syndrome, Keratitis-ichthyosis-deafness syndrome, Keratoderma palmoplantar, Hystrix-like ichthyosis
ELOVL4 Ichthyosis, spastic quadriplegia, and mental retardation
ZNF750 Seborrhea-like dermatitis with psoriasiform elements
SULT2B1 Ichthyosis, congenital, autosomal recessive 14
SLC27A4 Ichthyosis prematurity syndrome
SDR9C7 Ichthyosis, congenital, autosomal recessive 13
AP1B1 Keratitis-ichthyosis-deafness syndrome, autosomal recessive
CARD14 Pityriasis rubra pilaris, Psoriasis 2
SERPINB7 Palmoplantar keratoderma, Nagashima type
CERS3 Ichthyosis, congenital, autosomal recessive 9
GJB3 Erythrokeratodermia variabilis et progressiva 1
IL36RN Psoriasis pustular 14
CAST Peeling skin with leukonychia, acral punctate keratoses, cheilitis, and knuckle pads
FLG Ichthyosis vulgaris
FLG2 Peeling skin syndrome 6
ABCA12 Ichthyosis, congenital, autosomal recessive 4B (harlequin), 4A
LIPN Ichthyosis, congenital, autosomal recessive 8
STS Ichthyosis, X-linked
ABHD5 Chanarin-Dorfman syndrome
ALOX12B Ichthyosis, congenital, autosomal recessive 2
CTSB Keratolytic winter erythema
CDSN Peeling skin syndrome 1
SERPINB8 Peeling skin syndrome 5
MSMO1 Microcephaly, congenital cataract, and psoriasiform dermatitis
GBA Gaucher disease, type II, perinatal lethal
ASPRV1 Ichthyosis, lamellar, autosomal dominant
ALOXE3 Ichthyosis, congenital, autosomal recessive 3
SNAP29 Cerebral dysgenesis, neuropathy, ichthyosis, and palmoplantar keratoderma syndrome
CSTA Peeling skin syndrome 4
LOR Vohwinkel syndrome with ichthyosis

### Figure S7

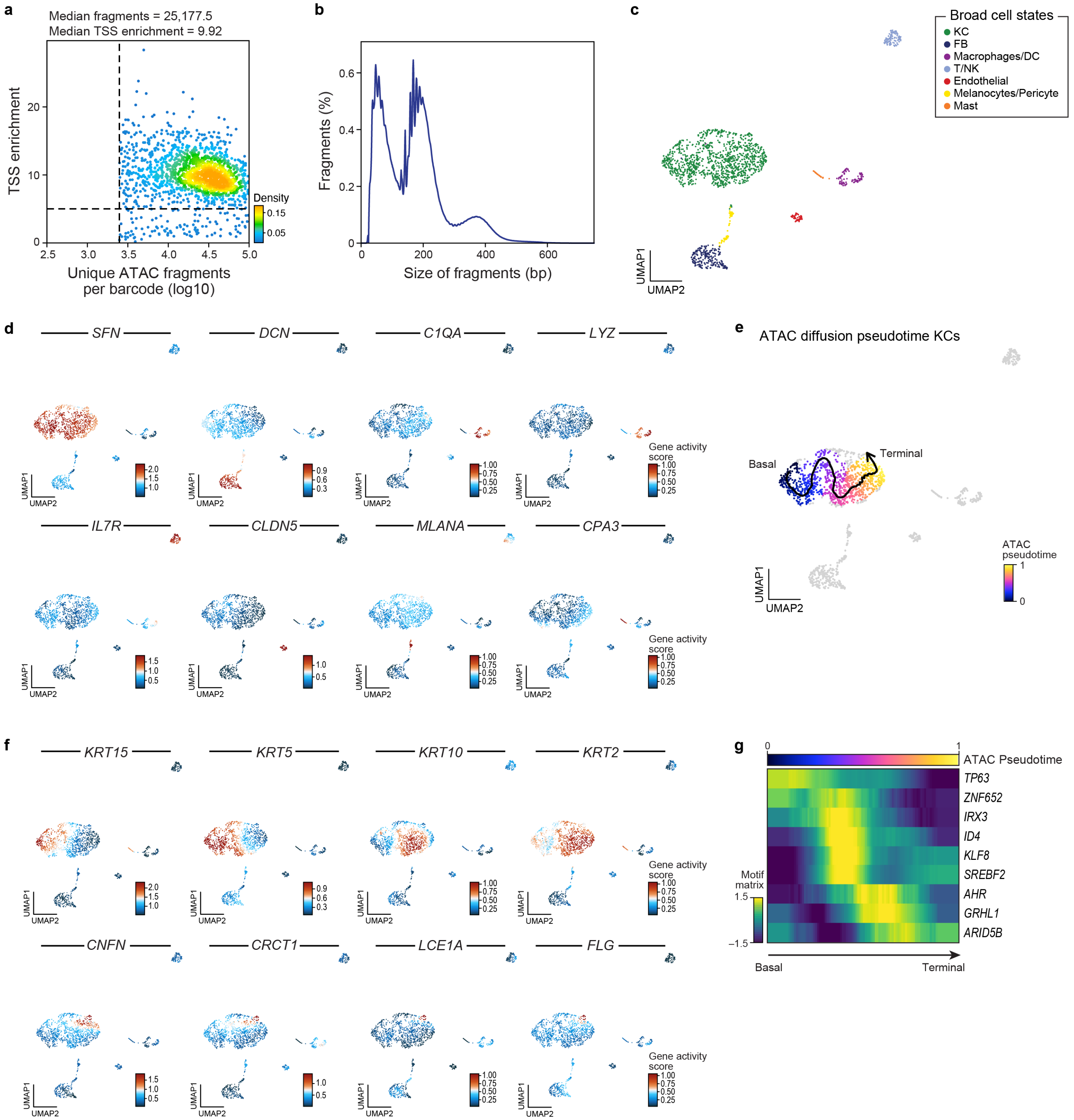
a
Median fragments = 25,177.5
Median TSS enrichment = 9.92
TSS enrichment
Unique ATAC fragments
per barcode (log10)
Density
b
Fragments (%)
Size of fragments (bp)
c
Broad cell states
KC
FB
Macrophages/DC
T/NK
Endothelial
Melanocytes/Pericyte
Mast
UMAP1
UMAP2
d
SFN
DCN
C1QA
LYZ
IL7R
CLDN5
MLANA
CPA3
Gene activity score
e
ATAC diffusion pseudotime KCs
Basal
Terminal
ATAC pseudotime
f
KRT15
KRT5
KRT10
KRT2
CNFN
CRCT1
LCE1A
FLG
g
ATAC Pseudotime
TP63
ZNF652
IRX3
ID4
KLF8
SREBF2
AHR
GRHL1
ARID5B
Motif matrix
Basal
Terminal

### Figure S8

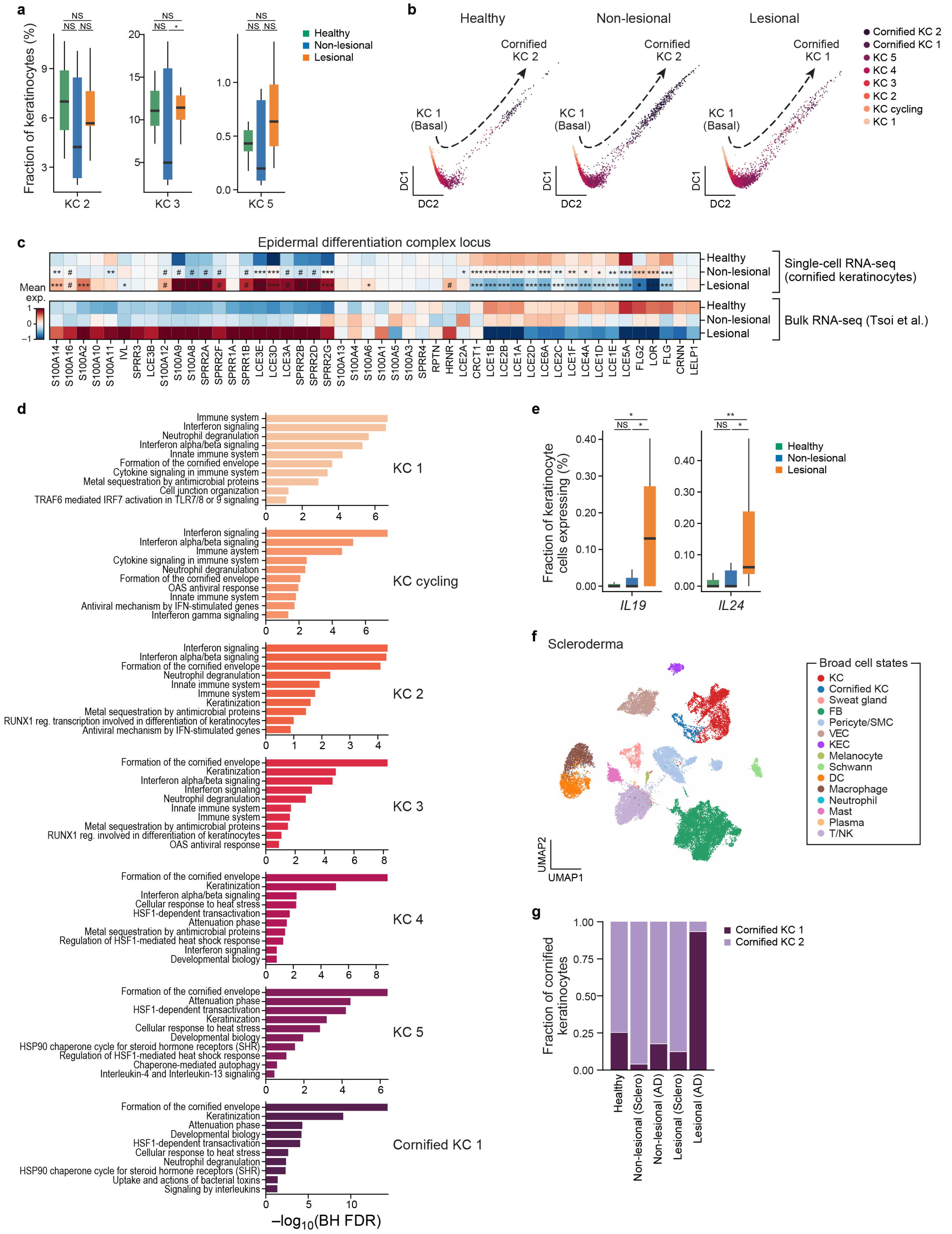

a
Fraction of keratinocytes (%)
KC 2
KC 3
KC 5
NS
Healthy
Non-lesional
Lesional
b
Healthy
Non-lesional
Lesional
Cornified KC 2
Cornified KC 1
KC 1 (Basal)
DC1
DC2
Cornified KC 2
Cornified KC 1
KC 5
KC 4
KC 3
KC 2
KC cycling
KC 1
c
Epidermal differentiation complex locus
Mean exp.
Healthy
Non-lesional
Lesional
Single-cell RNA-seq (cornified keratinocytes)
Bulk RNA-seq (Tsoi et al.)
S100A14 S100A16 S100A2 S100A10 S100A11 IVL SPRR3 LCE3B S100A12 S100A9 S100A8 SPRR2A SPRR2F SPRR1A SPRR1B LCE3E LCE3D LCE3A SPRR2B SPRR2D SPRR2G S100A13 S100A4 S100A6 S100A1 S100A5 S100A3 SPRR4 RPTN HRNR LCE2A CRCT1 LCE1B LCE2B LCE1A LCE2D LCE6A LCE2C LCE1F LCE4A LCE1D LCE1E LCE5A FLG2 LOR FLG CRNN LELP1
d
KC 1
Immune system
Interferon signaling
Neutrophil degranulation
Interferon alpha/beta signaling
Innate immune system
Formation of the cornified envelope
Cytokine signaling in immune system
Metal sequestration by antimicrobial proteins
Cell junction organization
TRAF6 mediated IRF7 activation in TLR7/8 or 9 signaling
KC cycling
Interferon signaling
Interferon alpha/beta signaling
Immune aystem
Cytokine signaling in immune system
Neutrophil degranulation
Formation of the cornified envelope
OAS antiviral response
Innate immune system
Antiviral mechanism by IFN-stimulated genes
Interferon gamma signaling
KC 2
Interferon signaling
Interferon alpha/beta signaling
Formation of the cornified envelope
Neutrophil degranulation
Innate immune system
Immune system
Keratinization
Metal sequestration by antimicrobial proteins
RUNX1 reg. transcription involved in differentiation of keratinocytes
Antiviral mechanism by IFN-stimulated genes
KC 3
Formation of the cornified envelope
Keratinization
Interferon alpha/beta signaling
Interferon signaling
Neutrophil degranulation
Innate immune system
Immune system
Metal sequestration by antimicrobial proteins
RUNX1 reg. involved in differentiation of keratinocytes
OAS antiviral response
KC 4
Formation of the cornified envelope
Keratinization
Interferon alpha/beta signaling
Cellular response to heat stress
HSF1-dependent transactivation
Attenuation phase
Metal sequestration by antimicrobial proteins
Regulation of HSF1-mediated heat shock response
Interferon signaling
Developmental biology
KC 5
Formation of the cornified envelope
Attenuation phase
HSF1-dependent transactivation
Keratinization
Cellular response to heat stress
Developmental biology
HSP90 chaperone cycle for steroid hormone receptors (SHR)
Regulation of HSF1-mediated heat shock response
Chaperone-mediated autophagy
Interleukin-4 and Interleukin-13 signaling
Cornified KC 1
Formation of the cornified envelope
Keratinization
Attenuation phase
Developmental biology
HSF1-dependent transactivation
Cellular response to heat stress
Neutrophil degranulation
HSP90 chaperone cycle for steroid hormone receptors (SHR)
Uptake and actions of bacterial toxins
Signaling by interleukins
$-\log_{10}$(BH FDR)
e
Fraction of keratinocyte cells expressing (%)
IL19
IL24
Healthy
Non-lesional
Lesional
f
Scleroderma
UMAP1
UMAP2
Broad cell states
KC
Cornified KC
Sweat gland
FB
Pericyte/SMC
VEC
KEC
Melanocyte
Schwann
DC
Macrophage
Neutrophil
Mast
Plasma
T/NK
g
Fraction of cornified keratinocytes
Cornified KC 1
Cornified KC 2
Healthy
Non-lesional (Sclero)
Non-lesional (AD)
Lesional (Sclero)
Lesional (AD)

### Figure S9

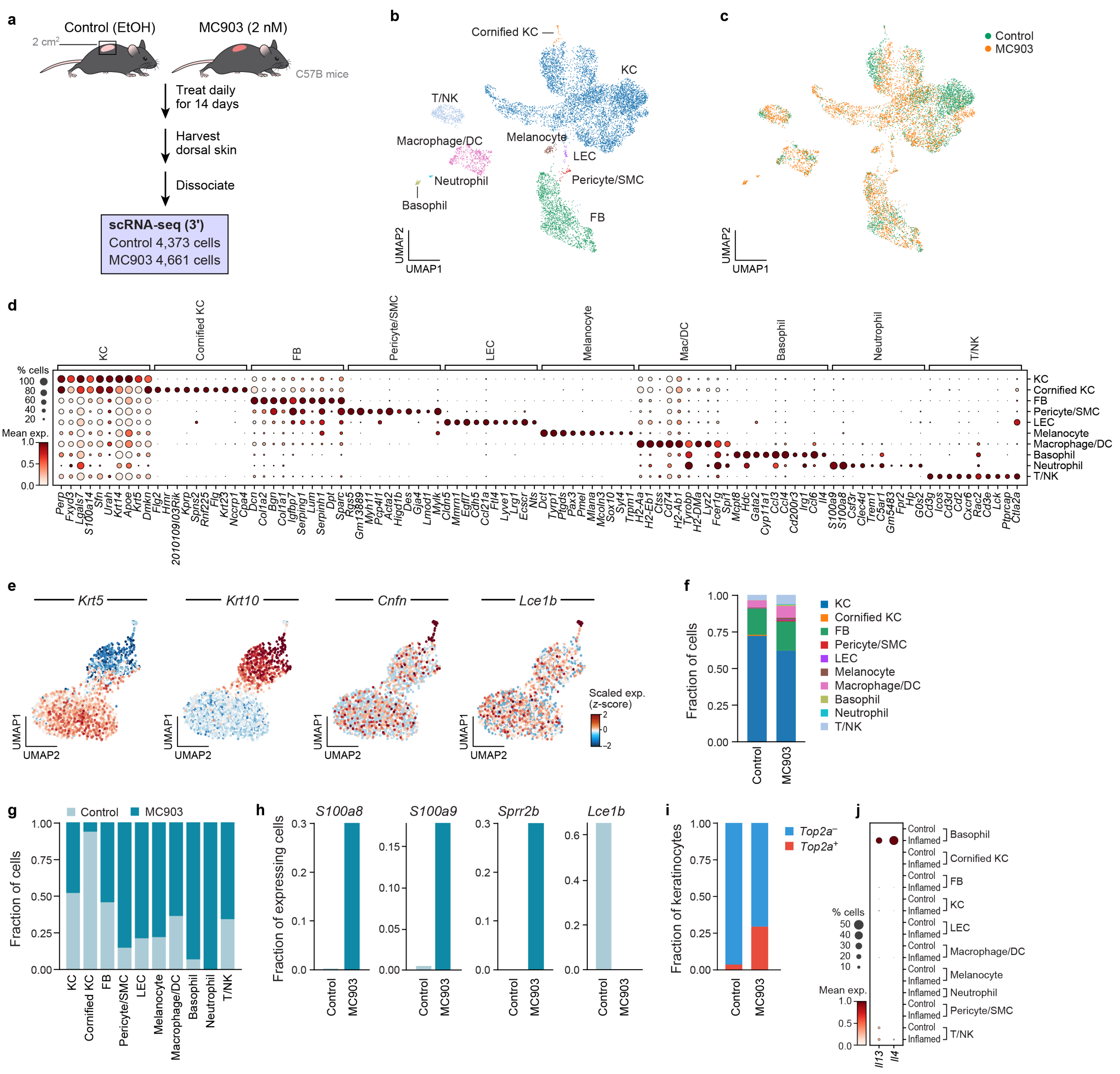
a
Control (EtOH)
MC903 (2 nM)
2 cm2
C57B mice
Treat daily for 14 days
Harvest dorsal skin
Dissociate
scRNA-seq (3')
Control 4,373 cells
MC903 4,661 cells
b
Cornified KC
KC
T/NK
Macrophage/DC
Melanocyte
LEC
Neutrophil
Pericyte/SMC
Basophil
FB
UMAP2
UMAP1
c
Control
MC903
d
KC
Cornified KC
FB
Pericyte/SMC
LEC
Melanocyte
Mac/DC
Basophil
Neutrophil
T/NK
% cells
Mean exp.
Macrophage/DC
e
Krt5
Krt10
Cnfn
Lce1b
Scaled exp. (z-score)
UMAP1
UMAP2
f
Fraction of cells
Control
MC903
g
Fraction of cells
h
S100a8
S100a9
Sprr2b
Lce1b
Fraction of expressing cells
i
Fraction of keratinocytes
Top2a−
Top2a+
j
Control
Inflamed
Il13
Il4

### Figure S10

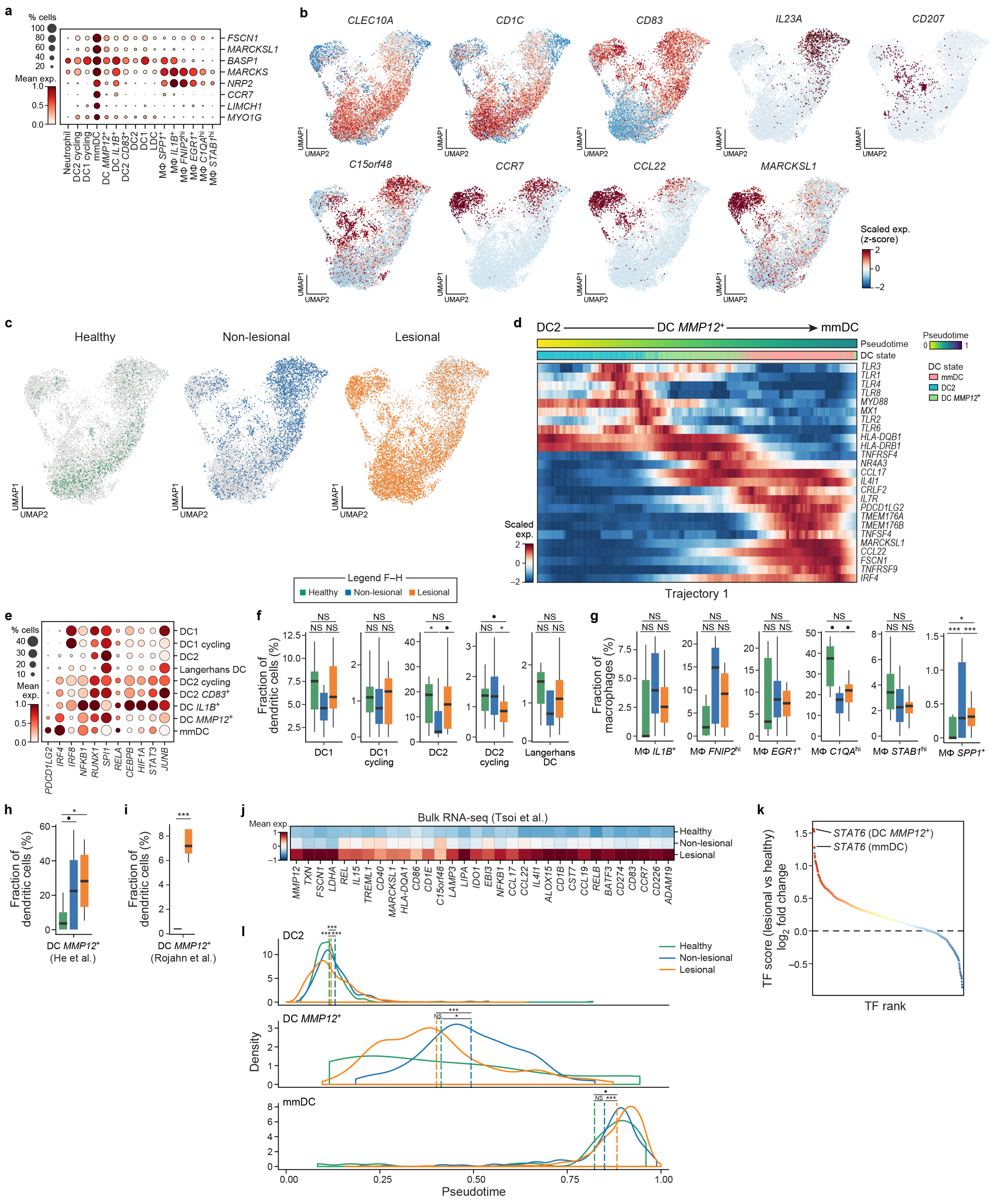
a
% cells
Mean exp.
FSCN1
MARCKSL1
BASP1
MARCKS
NRP2
CCR7
LIMCH1
MYO1G
Neutrophil
DC2 cycling
DC1 cycling
mmDC
DC MMP12+
DC IL1B+
DC2 CD83+
DC2
DC1
LDC
MΦ SPP1+
MΦ IL1B+
MΦ FNIP2hi
MΦ EGR1+
MΦ C1QAhi
MΦ STAB1hi
b
CLEC10A
CD1C
CD83
IL23A
CD207
C15orf48
CCR7
CCL22
MARCKSL1
UMAP1
UMAP2
Scaled exp. (z-score)
c
Healthy
Non-lesional
Lesional
d
DC2
DC MMP12+
mmDC
Pseudotime
DC state
Scaled exp.
Trajectory 1
Legend F–H
Healthy
Non-lesional
Lesional
e
DC1
DC1 cycling
DC2
Langerhans DC
DC2 cycling
DC2 CD83+
DC IL1B+
DC MMP12+
mmDC
PDCD1LG2
IRF4
IRF8
NFKB1
RUNX1
SPI1
RELA
CEBPB
HIF1A
STAT3
JUNB
f
Fraction of dendritic cells (%)
DC1
DC1 cycling
DC2
DC2 cycling
Langerhans DC
g
Fraction of macrophages (%)
MΦ IL1B+
MΦ FNIP2hi
MΦ EGR1+
MΦ C1QAhi
MΦ STAB1hi
MΦ SPP1+
h
Fraction of dendritic cells (%)
DC MMP12+
(He et al.)
i
DC MMP12+
(Rojahn et al.)
j
Bulk RNA-seq (Tsoi et al.)
Mean exp.
Healthy
Non-lesional
Lesional
MMP12
TXN
FSCN1
LDHA
REL
IL15
TREML1
CD40
MARCKSL1
HLA-DQA1
CD86
CD1E
C15orf48
LAMP3
LIPA
IDO1
EBI3
NFKB1
CCL17
CCL22
IL4I1
ALOX15
CD1B
CST7
CCL19
RELB
BATF3
CD274
CD83
CCR7
CD226
ADAM19
k
STAT6 (DC MMP12+)
STAT6 (mmDC)
TF score (lesional vs healthy) log2 fold change
TF rank
l
DC2
DC MMP12+
mmDC
Density
Pseudotime
Healthy
Non-lesional
Lesional

### Figure S11

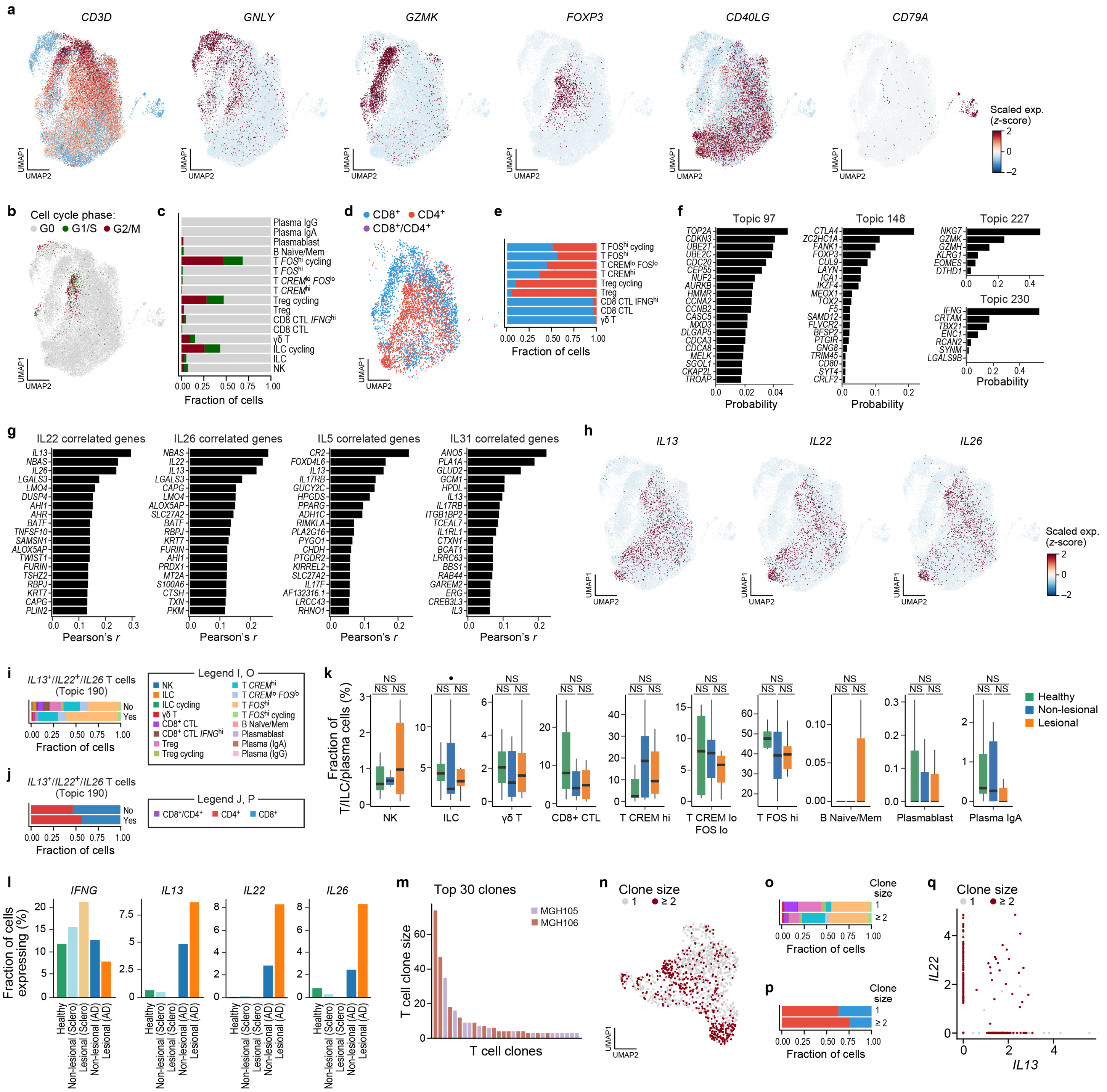
a
CD3D
GNLY
GZMK
FOXP3
CD40LG
CD79A
Scaled exp. (z-score)
b
Cell cycle phase:
G0 G1/S G2/M
c
Fraction of cells
d
CD8+ CD4+ CD8+/CD4+
e
Fraction of cells
f
Topic 97
Topic 148
Topic 227
Topic 230
Probability
g
IL22 correlated genes
IL26 correlated genes
IL5 correlated genes
IL31 correlated genes
Pearson's r
h
IL13
IL22
IL26
i
IL13+/IL22+/IL26 T cells (Topic 190)
Legend I, O
j
Legend J, P
k
Fraction of T/ILC/plasma cells (%)
Healthy
Non-lesional
Lesional
l
IFNG
IL13
IL22
IL26
Fraction of cells expressing (%)
m
Top 30 clones
MGH105
MGH106
T cell clone size
T cell clones
n
Clone size
o
p
q
Clone size
IL22
IL13

### Figure S12

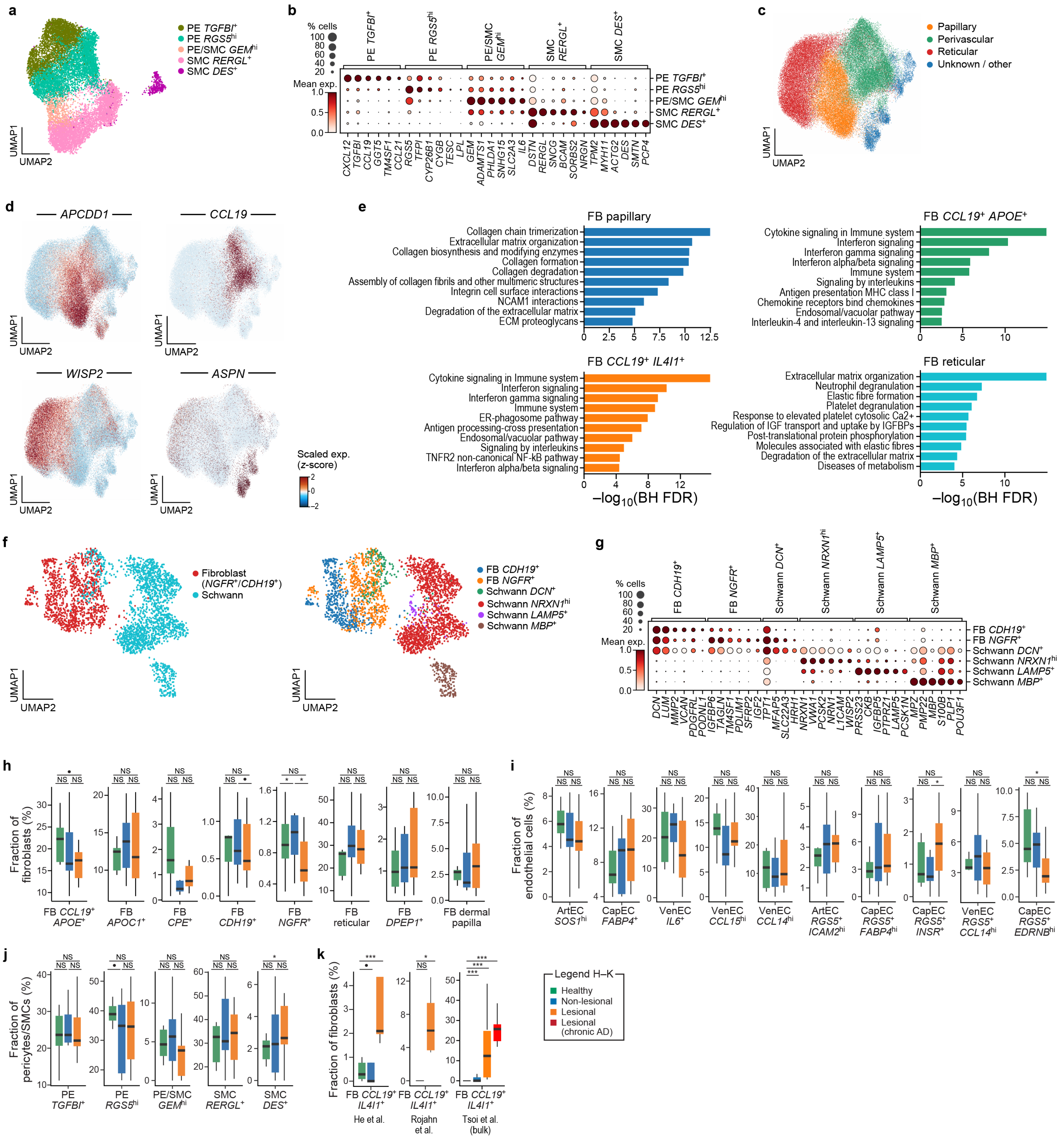

a
PE TGFBI+
PE RGS5hi
PE/SMC GEMhi
SMC RERGL+
SMC DES+
UMAP1
UMAP2
b
% cells
100
80
60
40
20
Mean exp.
1.0
0.5
0.0
PE TGFBI+
PE RGS5hi
PE/SMC GEMhi
SMC RERGL+
SMC DES+
CXCL12 TGFBI CCL19 GGT5 TM4SF1 CCL21 RGS5 TFPI CYP26B1 CYGB TESC LPL GEM ADAMTS1 PHLDA1 SNHG15 SLC2A3 IL6 DSTN RERGL SNCG BCAM SORBS2 NRGN TPM2 MYH11 ACTG2 DES SMTN PCP4
c
Papillary
Perivascular
Reticular
Unknown / other
UMAP1
UMAP2
d
APCDD1
CCL19
WISP2
ASPN
UMAP1
UMAP2
Scaled exp.
(z-score)
2
0
−2
e
FB papillary
Collagen chain trimerization
Extracellular matrix organization
Collagen biosynthesis and modifying enzymes
Collagen formation
Collagen degradation
Assembly of collagen fibrils and other multimeric structures
Integrin cell surface interactions
NCAM1 interactions
Degradation of the extracellular matrix
ECM proteoglycans
0 2.5 5 7.5 10 12.5
FB CCL19+ APOE+
Cytokine signaling in Immune system
Interferon signaling
Interferon gamma signaling
Interferon alpha/beta signaling
Immune system
Signaling by interleukins
Antigen presentation MHC class I
Chemokine receptors bind chemokines
Endosomal/vacuolar pathway
Interleukin-4 and interleukin-13 signaling
0 5 10
FB CCL19+ IL4I1+
Cytokine signaling in Immune system
Interferon signaling
Interferon gamma signaling
Immune system
ER-phagosome pathway
Antigen processing-cross presentation
Endosomal/vacuolar pathway
Signaling by interleukins
TNFR2 non-canonical NF-kB pathway
Interferon alpha/beta signaling
0 4 8 12
−log10(BH FDR)
FB reticular
Extracellular matrix organization
Neutrophil degranulation
Elastic fibre formation
Platelet degranulation
Response to elevated platelet cytosolic Ca2+
Regulation of IGF transport and uptake by IGFBPs
Post-translational protein phosphorylation
Molecules associated with elastic fibres
Degradation of the extracellular matrix
Diseases of metabolism
0 5 10
−log10(BH FDR)
f
Fibroblast
(NGFR+/CDH19+)
Schwann
FB CDH19+
FB NGFR+
Schwann DCN+
Schwann NRXN1hi
Schwann LAMP5+
Schwann MBP+
UMAP1
UMAP2
g
% cells
100
80
60
40
20
Mean exp.
1.0
0.5
0.0
FB CDH19+
FB NGFR+
Schwann DCN+
Schwann NRXN1hi
Schwann LAMP5+
Schwann MBP+
DCN LUM MMP2 VCAN PDGFRL PODNL1 IGFBP6 TAGLN TM4SF1 PDLIM1 SFRP2 IGF2 TPT1 MFAP5 SLC22A3 HRH1 NRXN1 VWA1 PCSK2 NRN1 L1CAM WISP2 PRSS23 CKB IGFBP5 PTPRZ1 LAMP5 PCSK1N MPZ PMP22 MBP S100B PLP1 POU3F1
h
Fraction of fibroblasts (%)
FB CCL19+ APOE+
FB APOC1+
FB CPE+
FB CDH19+
FB NGFR+
FB reticular
FB DPEP1+
FB dermal papilla
i
Fraction of endothelial cells (%)
ArtEC SOS1hi
CapEC FABP4+
VenEC IL6+
VenEC CCL15hi
VenEC CCL14hi
ArtEC RGS5+ ICAM2hi
CapEC RGS5+ FABP4hi
CapEC RGS5+ INSR+
VenEC RGS5+ CCL14hi
CapEC RGS5+ EDRNBhi
j
Fraction of pericytes/SMCs (%)
PE TGFBI+
PE RGS5hi
PE/SMC GEMhi
SMC RERGL+
SMC DES+
k
Fraction of fibroblasts (%)
FB CCL19+ IL4I1+
He et al.
FB CCL19+ IL4I1+
Rojahn et al.
FB CCL19+ IL4I1+
Tsoi et al. (bulk)
Legend H–K
Healthy
Non-lesional
Lesional
Lesional (chronic AD)

### Figure S13

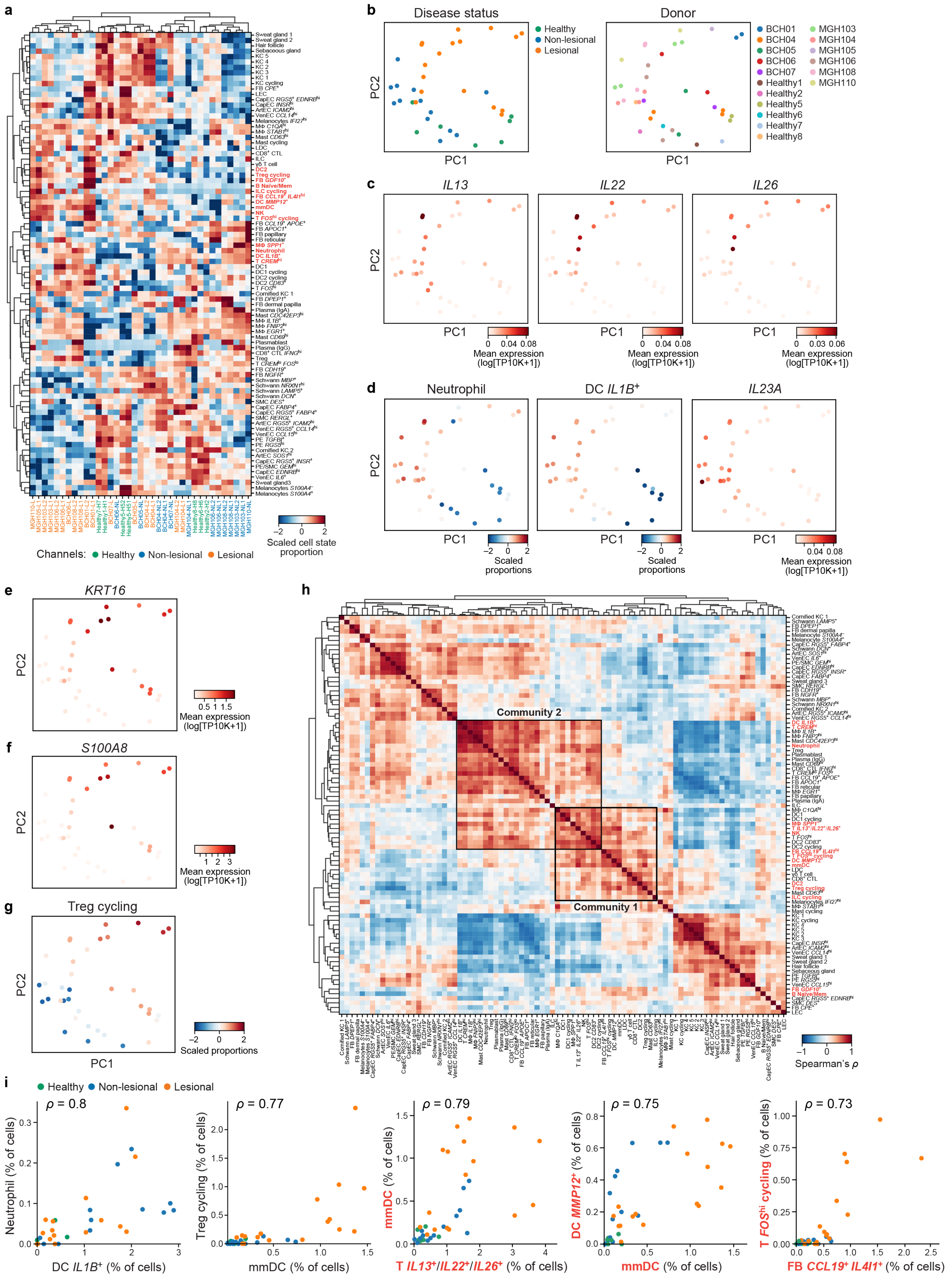
a
b
Disease status
Healthy
Non-lesional
Lesional
Donor
PC1
PC2
c
IL13
IL22
IL26
Mean expression (log[TP10K+1])
d
Neutrophil
DC IL1B+
IL23A
Scaled proportions
e
KRT16
f
S100A8
g
Treg cycling
h
Community 2
Community 1
Spearman's ρ
i
ρ = 0.8
ρ = 0.77
ρ = 0.79
ρ = 0.75
ρ = 0.73
Neutrophil (% of cells)
DC IL1B+ (% of cells)
Treg cycling (% of cells)
mmDC (% of cells)
T IL13+/IL22+/IL26+ (% of cells)
DC MMP12+ (% of cells)
T FOShi cycling (% of cells)
FB CCL19+ IL4I1+ (% of cells)
Scaled cell state proportion
Channels: Healthy Non-lesional Lesional

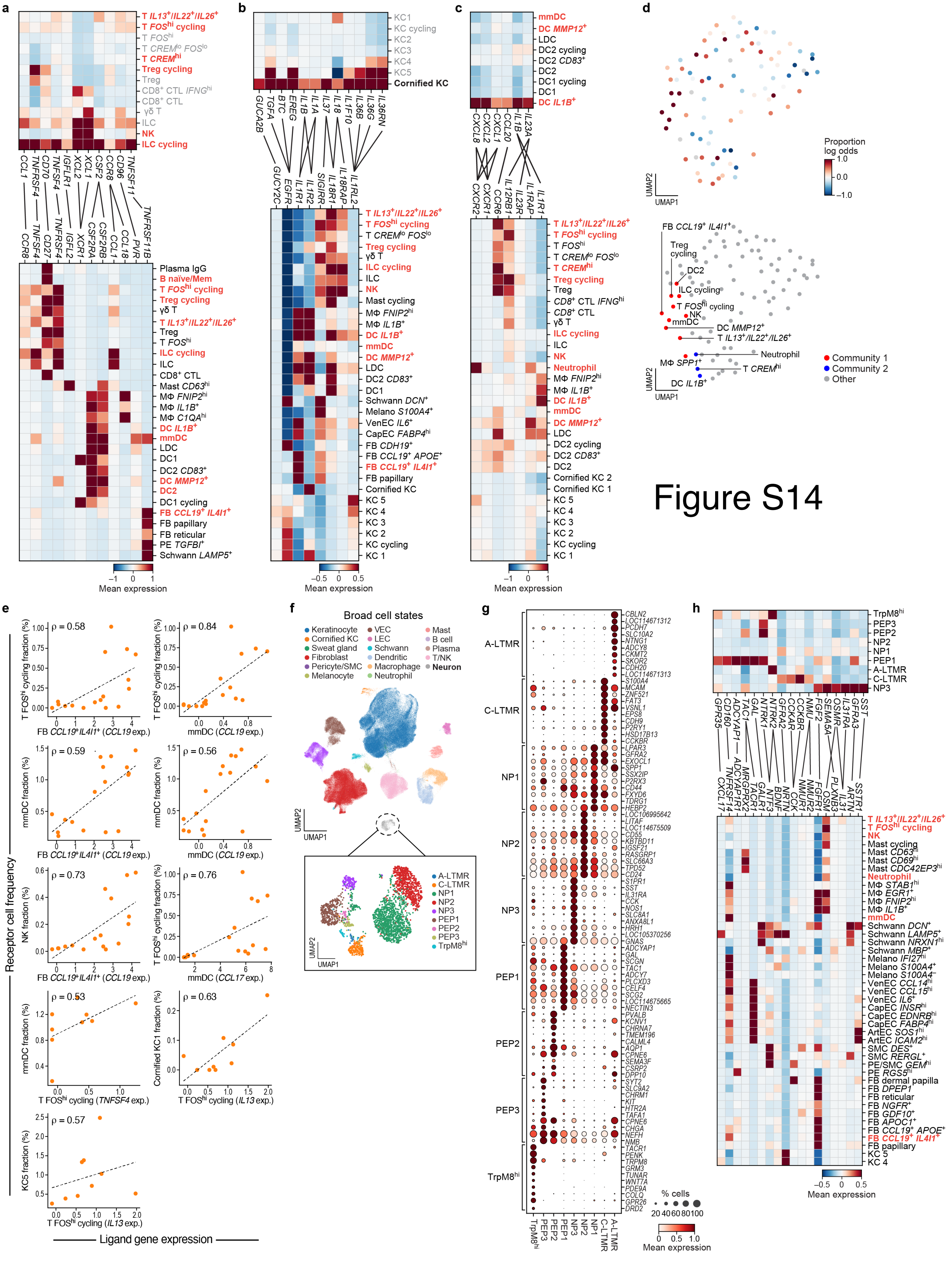
a
b
c
d
e
f
g
h
Mean expression
Proportion log odds
UMAP1
UMAP2
Community 1
Community 2
Other
Broad cell states
Keratinocyte
Cornified KC
Sweat gland
Fibroblast
Pericyte/SMC
Melanocyte
VEC
LEC
Schwann
Dendritic
Macrophage
Neutrophil
Mast
B cell
Plasma
T/NK
Neuron
A-LTMR
C-LTMR
NP1
NP2
NP3
PEP1
PEP2
PEP3
TrpM8hi
% cells
Receptor cell frequency
Ligand gene expression
ρ = 0.58
ρ = 0.84
ρ = 0.59
ρ = 0.56
ρ = 0.73
ρ = 0.76
ρ = 0.53
ρ = 0.63
ρ = 0.57

Figure S14

### Figure S15

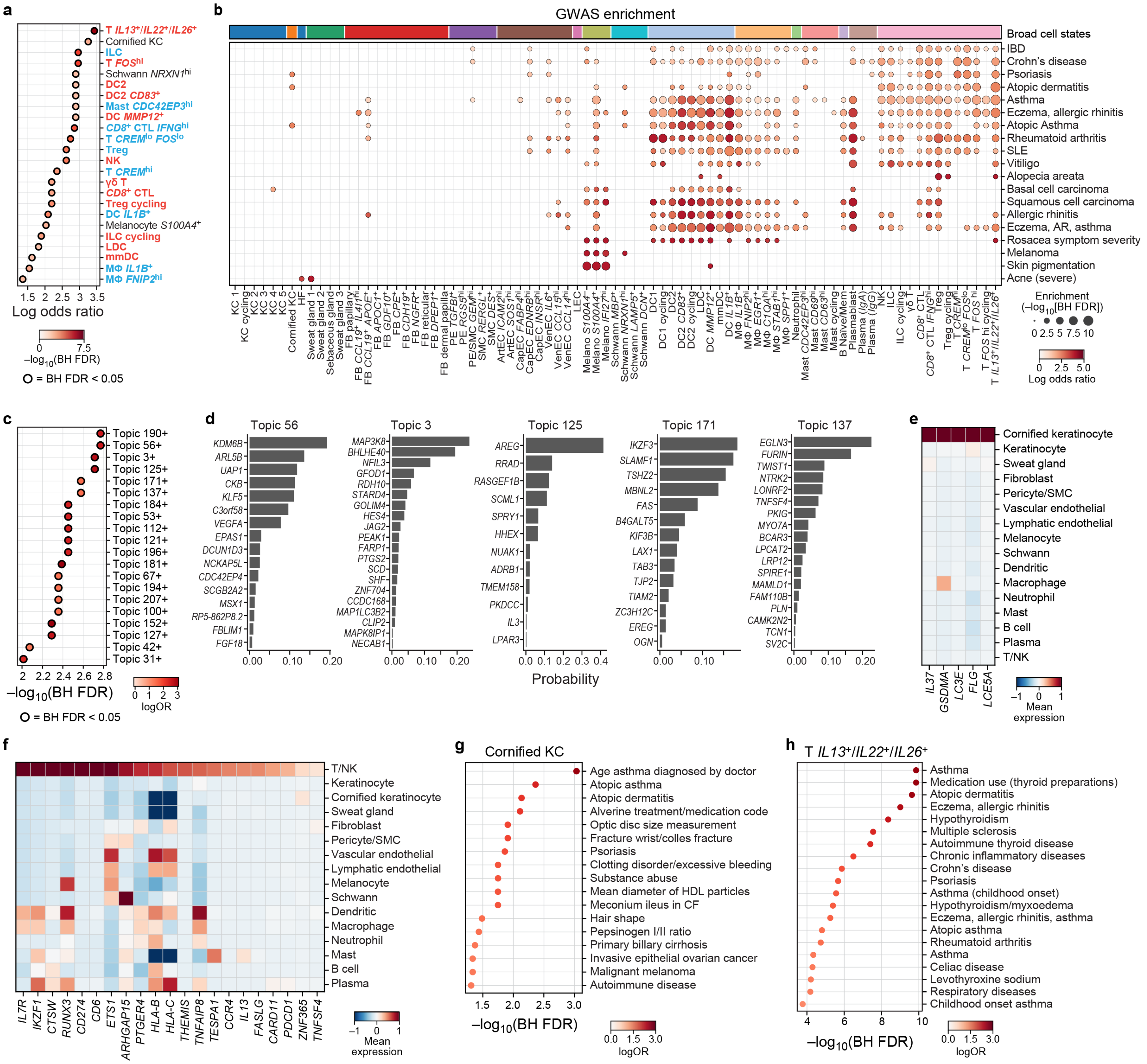

a
T IL13+/IL22+/IL26+
Cornified KC
ILC
T FOShi
Schwann NRXN1hi
DC2
DC2 CD83+
Mast CDC42EP3hi
DC MMP12+
CD8+ CTL IFNGhi
T CREMlo FOSlo
Treg
NK
T CREMhi
γδ T
CD8+ CTL
Treg cycling
DC IL1B+
Melanocyte S100A4+
ILC cycling
LDC
mmDC
MΦ IL1B+
MΦ FNIP2hi
Log odds ratio
–log10(BH FDR)
= BH FDR < 0.05
b
GWAS enrichment
Broad cell states
IBD
Crohn's disease
Psoriasis
Atopic dermatitis
Asthma
Eczema, allergic rhinitis
Atopic Asthma
Rheumatoid arthritis
SLE
Vitiligo
Alopecia areata
Basal cell carcinoma
Squamous cell carcinoma
Allergic rhinitis
Eczema, AR, asthma
Rosacea symptom severity
Melanoma
Skin pigmentation
Acne (severe)
Enrichment (–log10[BH FDR])
Log odds ratio
c
Topic 190+
Topic 56+
Topic 3+
Topic 125+
Topic 171+
Topic 137+
Topic 184+
Topic 53+
Topic 112+
Topic 121+
Topic 196+
Topic 181+
Topic 67+
Topic 194+
Topic 207+
Topic 100+
Topic 152+
Topic 127+
Topic 42+
Topic 31+
–log10(BH FDR)
logOR
= BH FDR < 0.05
d
Topic 56
Topic 3
Topic 125
Topic 171
Topic 137
Probability
e
Cornified keratinocyte
Keratinocyte
Sweat gland
Fibroblast
Pericyte/SMC
Vascular endothelial
Lymphatic endothelial
Melanocyte
Schwann
Dendritic
Macrophage
Neutrophil
Mast
B cell
Plasma
T/NK
IL37
GSDMA
LC3E
FLG
LCE5A
Mean expression
f
T/NK
Keratinocyte
Cornified keratinocyte
Sweat gland
Fibroblast
Pericyte/SMC
Vascular endothelial
Lymphatic endothelial
Melanocyte
Schwann
Dendritic
Macrophage
Neutrophil
Mast
B cell
Plasma
IL7R
IKZF1
CTSW
RUNX3
CD274
CD6
ETS1
ARHGAP15
PTGER4
HLA-B
HLA-C
THEMIS
TNFAIP8
TESPA1
CCR4
IL13
FASLG
CARD11
PDCD1
ZNF365
TNFSF4
Mean expression
g
Cornified KC
Age asthma diagnosed by doctor
Atopic asthma
Atopic dermatitis
Alverine treatment/medication code
Optic disc size measurement
Fracture wrist/colles fracture
Psoriasis
Clotting disorder/excessive bleeding
Substance abuse
Mean diameter of HDL particles
Meconium ileus in CF
Hair shape
Pepsinogen I/II ratio
Primary billary cirrhosis
Invasive epithelial ovarian cancer
Malignant melanoma
Autoimmune disease
–log10(BH FDR)
logOR
h
T IL13+/IL22+/IL26+
Asthma
Medication use (thyroid preparations)
Atopic dermatitis
Eczema, allergic rhinitis
Hypothyroidism
Multiple sclerosis
Autoimmune thyroid disease
Chronic inflammatory diseases
Crohn's disease
Psoriasis
Asthma (childhood onset)
Hypothyroidism/myxoedema
Eczema, allergic rhinitis, asthma
Atopic asthma
Rheumatoid arthritis
Asthma
Celiac disease
Levothyroxine sodium
Respiratory diseases
Childhood onset asthma
–log10(BH FDR)
logOR
